# Pleiotropy and epistasis within and between signaling pathways defines the genetic architecture of fungal virulence

**DOI:** 10.1101/2020.08.21.259861

**Authors:** Cullen Roth, Debra Murray, Alexandria Scott, Ci Fu, Anna F. Averette, Sheng Sun, Joseph Heitman, Paul M. Magwene

## Abstract

Cryptococcal disease is estimated to affect nearly a quarter of a million people annually. Environmental isolates of *Cryptococcus deneoformans*, which make up 15 to 30% of clinical infections in temperate climates such as Europe, vary in their pathogenicity, ranging from benign to hyper-virulent. Key traits that contribute to virulence, such as the production of the pigment melanin, an extracellular polysaccharide capsule, and the ability to grow at human body temperature have been identified, yet little is known about the genetic basis of variation in such traits. Here we investigate the genetic basis of melanization, capsule size, thermal tolerance, oxidative stress resistance, and antifungal drug sensitivity using quantitative trait locus (QTL) mapping in progeny derived from a cross between two divergent *C. deneoformans* strains. Using a “function-valued” QTL analysis framework that exploits both time-series information and growth differences across multiple environments, we identified QTL for each of these virulence traits and drug susceptibility. For three QTL we identified the underlying genes and nucleotide differences that govern variation in virulence traits. One of these genes, *RIC8*, which encodes a regulator of cAMP-PKA signaling, contributes to variation in four virulence traits: melanization, capsule size, thermal tolerance, and resistance to oxidative stress. Two major effect QTL for amphotericin B resistance map to the genes *SSK1* and *SSK2*, which encode key components of the HOG pathway, a fungal-specific signal transduction network that orchestrates cellular responses to osmotic and other stresses. We also discovered complex epistatic interactions within and between genes in the HOG and cAMP-PKA pathways that regulate antifungal drug resistance and resistance to oxidative stress. Our findings advance the understanding of virulence traits among diverse lineages of *Cryptococcus*, and highlight the role of genetic variation in key stress-responsive signaling pathways as a major contributor to phenotypic variation.

**Author summary:** Different environmental isolates (strains) of the same microbial species can vary greatly in their ability to cause disease, ranging from avirulent to hypervirulent. What makes some strains deadly pathogens, while others are relatively benign? This study describes the characterization of key genetic differences that underlie variation in traits thought to promote virulence in *Cryptococcus deneoformans*, a wide-spread opportunistic fungal pathogen. Using a combination of quantitative genetic and molecular genetic approaches we dissected the genetic architecture of virulence-related cellular traits (melanin production and the production of a polysaccharide capsule), physiological responses to stress (tolerance of thermal, oxidative, and osmotic stress), and sensitivity to multiple antifungal drugs. Strikingly we find that variation in most of these traits is governed by a small number of genetic differences that modify the function of two major cell signaling networks, cyclic AMP–Protein Kinase A (cAMP-PKA) signaling and a fungal specific MAP-kinase cascade called the high osmolarity glycerol (HOG) pathway. Similar to recent studies in a number of other fungal species, our findings point to an outsize role for a small number of highly pleiotropic signaling pathways in potentiating phenotypic variation both within and between fungal species.

## Introduction

Over the last two decades, fungal species have emerged as major threats and pathogens [1, 2], affecting endangered plant and animal species [3–7], reducing crop yields [8, 9], and causing human illness [10–13]. The propensity of fungal pathogens to cause disease is a complex outcome dependent on a variety of underlying physiological features that facilitate survival in stressful host niches, such as the ability to forage and acquire nutrients [14], to tolerate bombardment from reactive oxygen species [15], and to mount a successful defense against (or evade) the host immune system [16, 17]. Significant progress has been made with respect to understanding the cell and molecular biology of fungal pathogenesis; numerous virulence-related traits and key genes and pathways that regulate these traits have been identified for many fungal pathogens [18–29]. Similarly, the availability of low-cost, high-throughput genome sequencing has greatly advanced the understanding of genetic variation and population structure for many fungal pathogens [30–36]. However, despite advances in both the molecular genetics of fungal pathogenesis and the genomics of pathogenic species, for most fungal pathogens we have a limited understanding of the genetic changes between isolates that contribute to differences in virulence traits [37, 38].

Basidiomycete fungi of the genus *Cryptococcus* are important human pathogens, estimated to affect nearly a quarter of a million people worldwide annually [39]. The majority of cryptococcal infections occur in individuals with compromised or suppressed immune systems, such as those combating AIDS/HIV or organ transplant recipients, however infections in seemingly healthy people have also been reported [40–44]. If untreated, cryptococcal meningitis is uniformly fatal, and current estimates of mortality rates for individuals receiving treatment for cryptococcosis vary by region, from 10 – 30% in North America to as high as 50 – 70% in parts of sub-Saharan Africa [39, 45, 46]. Most cases of cryptococcosis are due to infections of *Cryptococcus neoformans*, but the sister species *Cryptococcus deneoformans* (formerly referred to as *C. neoformans* var. *neoformans* serotype D) is responsible for a significant number of clinical cases in temperate regions of the world, and mixed infections of both *C. neoformans* and *C. deneoformans* have been reported [47–50]. In addition to their clinical relevance, *Cryptococcus* species are attractive model organisms for studying traits associated with virulence due to their experimental tractability, including a well characterized sexual cycle featuring recombination and vegetative growth as haploid yeasts [19, 51, 52], methods for transformation and genetic engineering, [53–56], and a large panel of gene deletion strains [21].

Key *Cryptococcus* virulence traits include melanization, resistance to oxidative stress, formation of an extracellular capsule, and thermal tolerance [57–63]. *Cryptococcus* species are opportunistic rather than obligate pathogens, and not all *Cryptococcus* species or strains exhibit the full complement of traits that are thought to be required for pathogenesis in animal hosts [64]. Furthermore, many of these traits, such as the production of melanin and the polysaccharide capsule, are likely to impose a significant metabolic cost. Given this, there has been considerable interest in the selective forces that have contributed to the origin and maintenance of virulence traits. The “accidental pathogen hypothesis” suggests that these traits evolved due to interactions with microbial predators and physiological stresses within non-pathogenic niches; thus virulence-associated traits are likely to have dual roles in the natural environment and within animal hosts [1, 65–67]. For example, the biosynthesis of melanin, a hydrophobic high-molecular weight black or brown pigmented polymer, buffers *Cryptococcus* cells from thermal stress and protects cells against solar radiation [68, 69]. Within the host niche, this pigment prevents damage from reactive oxygen species [28, 57, 70]. The extracellular polysaccharide capsule is thought to protect cells from being phagocytosed by amoeboid protozoans, natural predators of *Cryptococcus* [71]. In the host environment, the capsule protects cells from the host immune response, including phagocytosis by macrophages and oxidative stress, and shed capsule material has a variety of activities on host immune cells [63, 72, 73]. Thermal tolerance, which could be selected for by extreme seasonal temperatures, allows for the infection of mammalian and avian hosts with high body temperatures [74–77]. *Cryptococcus* species most often associated with human disease, *C. neoformans*, *C. deneoformans*, and *C. gattii* display the highest thermal tolerance [64].

Another important trait, resistance to antifungal drugs, is not considered a virulence trait per se, because it is not necessary for establishing an initial infection within a host. However, antifungal resistance can lead to recurring disease and is thus a clinically relevant trait [62, 78, 79]. Amphotericin B is one of the few drugs effective in the treatment of cryptococcosis [80, 81], killing fungal cells by binding to and sequestering ergosterol from the bilipid membrane [82, 83]. While resistance to amphotericin B is rare [84], a recent examination of clinical isolates observed an increase in the amphotericin B minimum inhibitory concentrations compared to inhibitory concentrations taken from the same region of study ten years earlier [85]. Globally, there are an increasing number of reports documenting rises in antifungal resistance [86, 87] and understanding the genetic architecture of drug susceptibility is integral to combating this growing trend.

There is considerable variation in virulence-associated traits and antifungal drug resistance both within and between *Cryptococcus* species [88–97]. One of the most powerful approaches for dissecting the genetic basis of phenotypic variation is quantitative trait locus (QTL) mapping [98, 99]. QTL mapping has been employed extensively in the model yeast *Saccharomyces cerevisiae* [100–106] and has been used to explore the genetic basis of virulence-related traits for a number of fungal plant pathogens [107–109]. However, there have been relatively few QTL studies in human fungal pathogens – one in *Aspergillus* [110] and two previous QTL studies in *Cryptococcus* [111, 112]. In these important pathogens, QTL mapping has the potential to enhance our understanding of the genetic basis of virulence.

A common, though not universal, characteristic of many microbial QTL mapping studies is the use of microbial growth as a proxy for physiological responses to different environmental conditions or stresses [105]. However, the use of growth as a trait presents some important challenges. For example, two strains may exhibit drastically different lag times and exponential growth rates yet still reach the same final population density. Furthermore, classic mathematical models of microbial growth, such as the Gompertz equation or logistic growth models [113], often fail to capture the real world complexities of microbial growth [114]. Generally speaking, the population density of a microbial culture is a complex function dependent upon both time and the magnitude of exposure to a stress or environment. Such data are often termed longitudinal or function valued and a body of statistical methods have been developed for function-valued data in which the order and spacing of data is retained [115–121]. Studies which use frameworks for the analysis of function-valued data can be found across the fields of biology including ecology, developmental biology, and crop genetics [122–129]. With respect to QTL mapping, function-valued methods have been shown to increase the ability to detect QTL [122, 124, 130–136].

Here we describe the genetic architecture of six clinically important and complex phenotypic traits in *C. deneoformans*: melanization, capsule size, thermal tolerance, growth under oxidative stress, and resistance to the antifungal drugs amphotericin B and fludioxonil. Based on a mapping population derived from a cross between a laboratory strain (XL280; [137]) and an environmental isolate (431α; [138]), we employed genome-wide sequencing and function-valued QTL mapping to identify genetic differences that underlie variation in each of the above traits. We discovered a major QTL with highly pleiotropic effects on melanization, capsule size, high temperature growth, and resistance to oxidative stress. We identified a likely causal variant for this shared QTL, a premature stop codon in the gene *RIC8*, a component of the cyclic AMP-protein kinase A (cAMP-PKA) signaling pathway. Interestingly, allelic variation at *RIC8* has antagonistic effects with regard to virulence potential, increasing tolerance to high temperatures while decreasing melanization. We also identified two QTL that underlie amphotericin B susceptibility, and mapped the likely causal variants to the genes *SSK1* and *SSK2*, components of the high-osmolarity glycerol (HOG) pathway. Epistatic interactions within the HOG pathway, and between the HOG and cAMP-PKA pathways, also contribute to variability in drug resistance, thermal tolerance, and oxidative stress resistance. This study highlights the importance of genetic variation in key signal transduction pathways that regulate stress responses in *Cryptococcus* and other fungi, and illustrates the complex effects that such variants may have with respect to virulence potential.

## Results

### A high resolution genetic mapping population

We generated an F_1_ mapping population by crossing the *C. deneoformans* strains XL280**a** and XL280αSS [92, 137] with the environmental isolate, 431α [92, 139, 140] in α–α unisexual and **a**–α bisexual matings [92]. The haploid genomes of the parental strains and 101 segregants were sequenced at approximately 64*×* coverage. Following filtering, 92,103 sites were identified that differ between the parental strains, and genotypes at each of these variable sites were called for each segregant based on mapping to the XL280α reference genome [137]. Variable sites were collapsed into unique haploblocks based on genetic exchange events, resulting in 3,108 unique haploblocks. The average size of the haploblocks was 5.4 kb (approximately 1 cM; [141]) with a maximum and minimum size of 6.3 and 4.4 kb, respectively (S1 Fig). This set of 101 segregants, parental strains, and their genotypic states at each of the 3,108 haploblocks served as the mapping population for subsequent QTL analyses.

### QTL for melanization

Melanization is an important phenotype related to virulence, and a previous study that utilized the same set of progeny used here observed significant variation in melanin production [92]. Melanization was quantified from scanned images of colonies grown on L-DOPA plates by calculating the mean grayscale intensity (the amount of light reflected off of a colony) of each segregant. The parental strains differ in their production of melanin – the XL280**a** parental strain has an opaque, beige appearance and the 431α parental strain grew as a dark brown colony (Fig 1A). There is significant variation in the production of melanin among the segregants, with most progeny exhibiting melanization intermediate between the two parental phenotypes (Fig 1A). Less than 8% of progeny exhibited transgressive melanin phenotypes that were more pigmented than the 431α parental strain or lighter than the XL280**a** parental strain. QTL mapping of the melanin phenotype identified a single large peak on chromosome 14 (Fig 1B). Segregants with the XL280**a** genotype at this locus had lighter colonies (higher mean intensity), while segregants with the 431α allele produced darker colonies (lower mean intensity, Fig 1C and 1D). Based on the regression model used for mapping, this QTL explains 39% of the variation in melanization in this cross.

**Fig 1.**
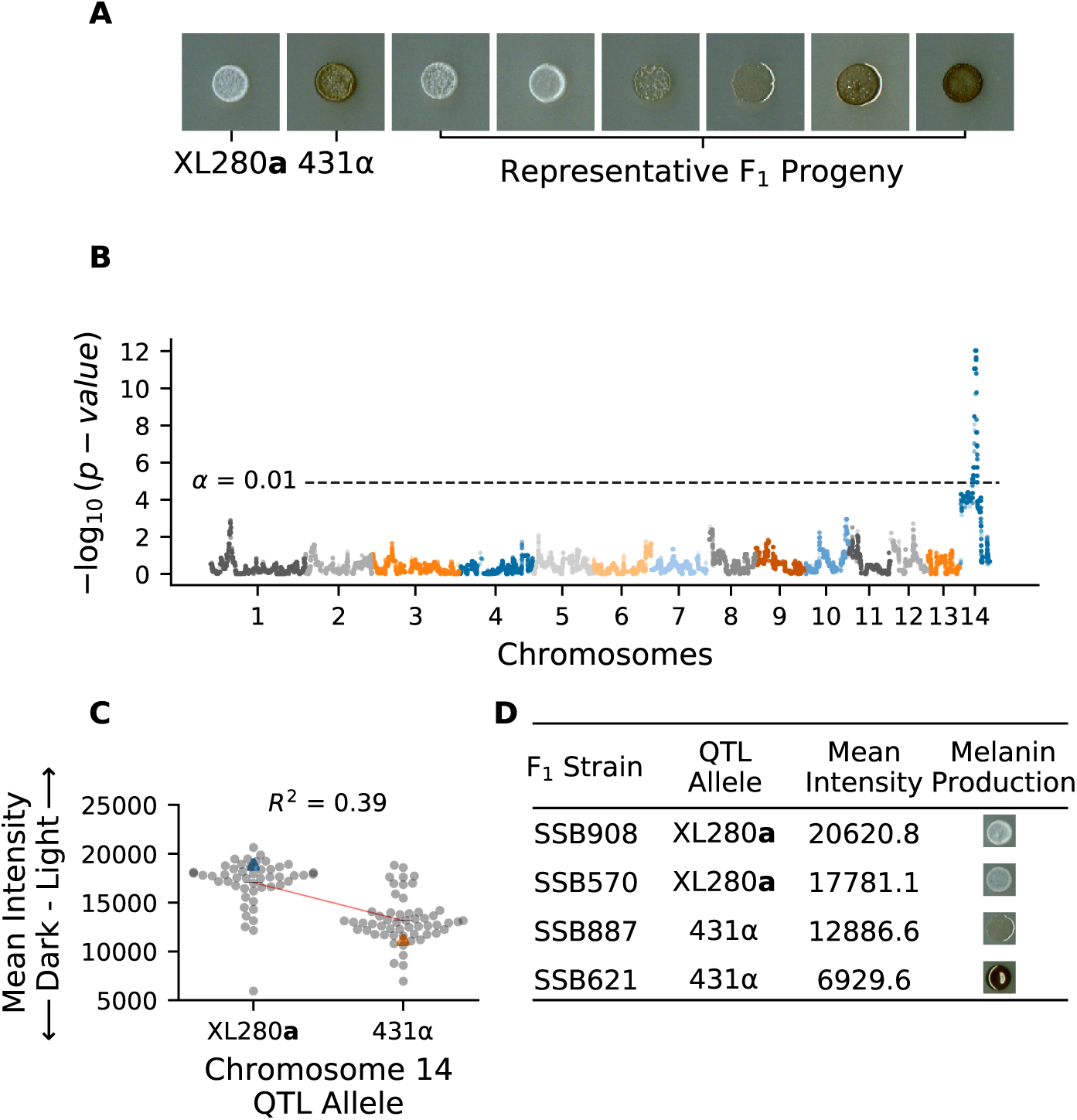
QTL analysis of variation in melanin production. **A**) Melanization phenotypes of parental strains – XL280**a** and 431α– and examplar phenotypes of their segregants. **B**) Manhattan plot of the association between genotype and melanin production. The x-axis represents chromosomal locations of haploblocks and the y-axis represents the strength of association between genotype and variation in melanization. The significance threshold (dashed horizontal line) was determined via permutation. **C**) Mean grayscale intensity (y-axis; arbitrary units) of segregants (gray dots) grown on L-DOPA plates as a function of genotype (x-axis) at the QTL peak on chromosome 14. Blue and orange triangles mark the parental phenotypes while black horizontal lines denote the phenotypic means by allele. The red line represents a regression model relating phenotype to genotype; this regression model explains 39% of the variation in melanin production. **D**) QTL allele, mean intensity, and melanin production of example F_1_ strains.

### QTL for variation in capsule diameter

The production of a polysaccharide capsule is another well-studied virulence trait in *Cryptococcus*. India ink stained cells from each segregant were imaged using brightfield microscopy, and cell body and capsule diameter were measured. There was a strong allometric relationship between capsule diameter and cell size (Fig 2A). To account for this “size effect” we regressed capsule diameter on the combined cell and capsule diameter, and used the residuals from this relationship as a measure of size-standardized capsule size (Fig 2B). A similar model, comparing cell diameter to the cell and capsule diameter was also calculated (S2 Fig).

**Fig 2.**
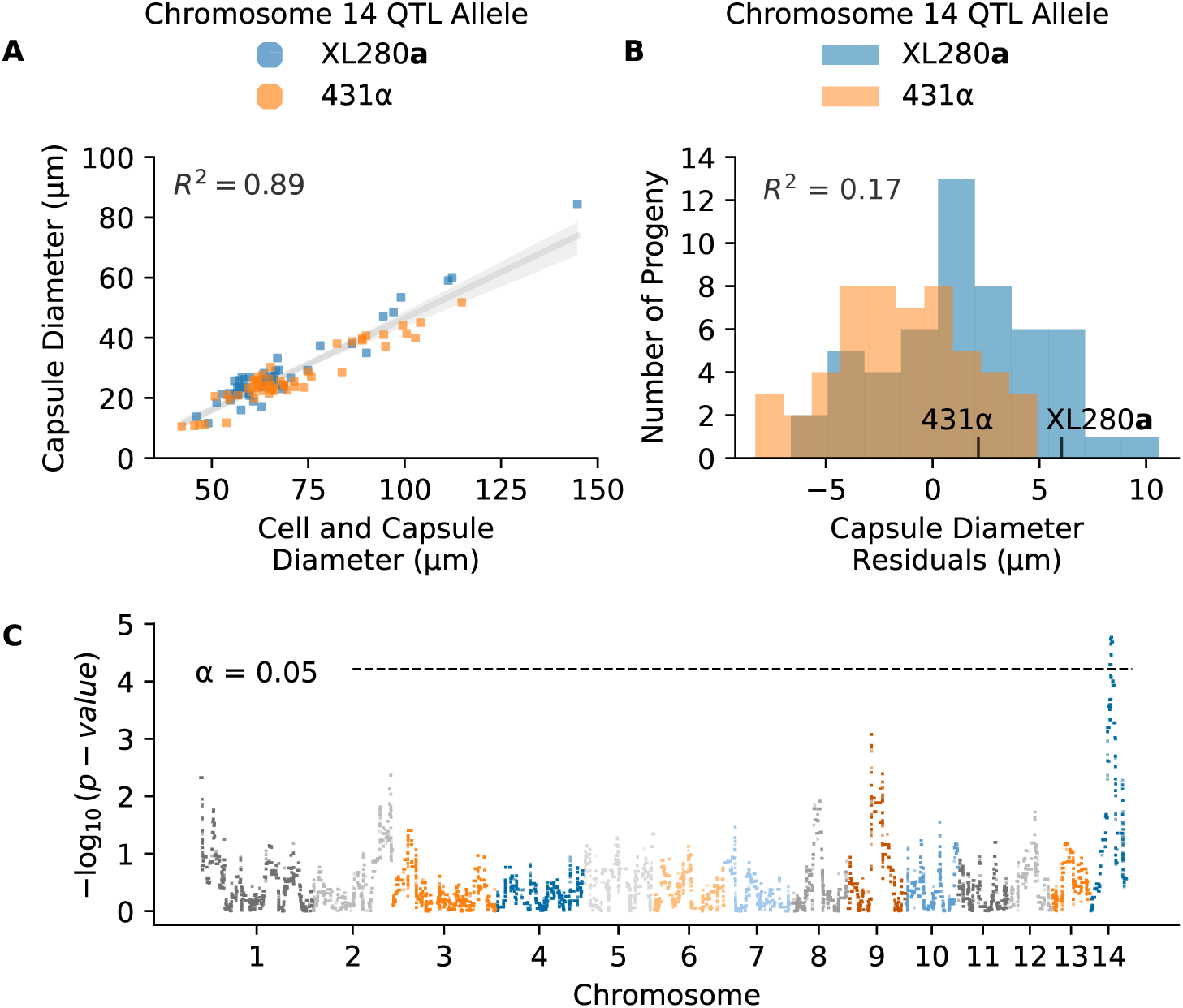
QTL analysis of variation in capsule diameter residuals. **A**) Measurements of the average cell and capsule diameters (x-axis) and the calculated capsule diameter (y-axis) per segregant colored by the chromosome 14 QTL allele in **C**. **B**) Histogram of capsule diameter residuals calculated from the linear regression model in **A**, separated by chromosome 14 QTL allele. **C**) Manhattan plot of the association between genotype and capsule diameter residuals. The x-axis represents chromosomal locations of haploblocks and the y-axis represents the strength of association between genotype and variation in capsule diameter residuals.

QTL mapping of the standardized capsule size identified a single significant peak on chromosome 14 (Fig 2C). Heritablity at this locus was estimated to be 17%. At the peak of this QTL, segregants with the XL280**a** genotype had larger (positive) capsule diameter residuals compared to sibling strains with the 431α allele (Fig 2B).

### Negative transgressive segregation in temperature and amphotericin B tolerance

Microbial stress responses are dependent on both the intensity of exposure and time since exposure. In order to capture both aspects of such responses to thermal stress and antifungal drugs we employed an automated phenotyping framework to measure microbial growth over time across multiple environmental conditions. For each of the segregants and parental strains, growth in liquid media was measured on an absorbance microplate reader for a total of eleven experimental conditions consisting of combinations of temperature (30, 37, and 39*^◦^*C) and amphotericin B (concentrations of 0, 0.075, 0.125, and 0.175 *µ*g/ml). These conditions were chosen to maximize the phenotypic variation within the mapping population. In each experimental condition, the optical density (OD_595_*_nm_*) was measured at 15-minute intervals for 72 hours. Each set of time series measurements was treated as a growth curve and four replicate growth curves were measured per segregant. After normalization and base-lining, total growth was estimated as the area under each growth curve. Fig 3 represents the median growth curve across replicates for each segregant at each combination of temperature and amphotericin B concentration.

**Fig 3.**
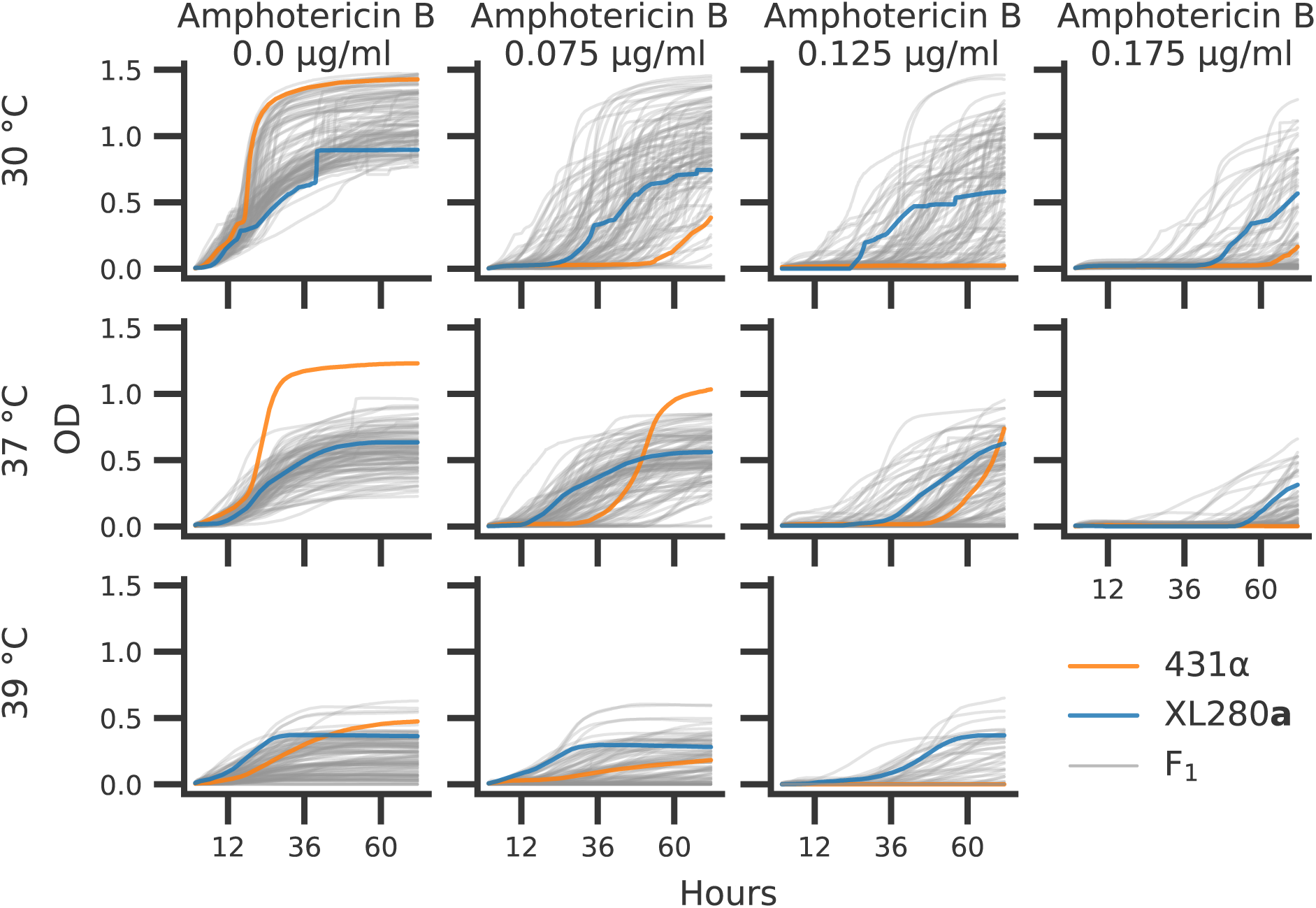
*C. deneoformans* growth curves. Parental strains and progeny were assayed for growth across combinations of temperatures (rows) 30*^◦^*, 37*^◦^*, 39*^◦^*C and concentrations of amphotericin B (columns) at 0.0, 0.075, 0.125, and 0.175 *µ*g/ml. Optical density (OD_595_*_nm_*, y-axis) was measured every 15 minutes for 72 hours (x-axis). The median optical density across replicates is shown for the parental strains, XL280**a** (blue curve) and 431α (orange curve), and the F_1_ segregants (grey curves).

There was significant variation in the growth trajectories across the eleven temperature by amphotericin B conditions. At the permissive conditions of 30*^◦^*C and no amphotericin B, most of the segregants growth curves fell near or between the parental growth curves. Conversely, at 37*^◦^*C without amphotericin B, the parental strain 431α outgrew the other parental strain, XL280**a**, as well as all of the segregants. In this high temperature condition 35% of segregants outgrew the XL280**a** parental strain. In most other combinations of temperature and amphotericin B stress, F_1_ progeny displayed negative transgressive segregation, with less total growth compared to the parental strains.

At 30*^◦^*C, across amphotericin B concentrations of 0.075, 0.125 and 0.175 *µ*g/ml, the 431α progenitor strain grew poorly and across these experimental conditions only 33, 24, and 19% of segregants (respectively) outgrew the XL280**a** parental strain. Surprisingly, 431α, when exposed to a combination of modest thermal stress (37*^◦^*C) and moderate amphotericin B concentrations (0.075 and 0.125 *µ*g/ml), grows better than when exposed to drug stress alone. At 37*^◦^*C in conditions of 0.075, 0.125 and 0.175 *µ*g/ml of amphotericin B, 44, 24, and 18% of segregants outgrow the XL280**a** progenitor strain, respectively. At 39*^◦^*C the parental strains had similar total growth with the 431α displaying a greater final OD. Across amphotericin B conditions at 39*^◦^*C, the XL280**a** parent outgrew the 431α strain, and only a modest number of offspring (*∼*7%) outgrew either parental strain. Taken as a whole, these data revealed a temporally dynamic and varying response to temperature and antifungal stress.

### Dynamic QTL underlying temperature stress and resistance to amphotericin B

A common approach to identify QTL associated with variation in microbial growth is to map the maximum growth rate or the population density at a specific time point and regress this value across variable genetic loci. This approach however fails to capture genotype-phenotype associations that change across time. Time varying traits are often referred to as function-valued [120]. Here, a function-valued, marker-regression approach was employed to quantify the relationship between genotype and growth phenotypes at each variable haploblock across the 72-hour time courses of each temperature and amphotericin B combination.

Temporally dependent QTL underlying variation across each of the eleven experimental combinations of temperature and concentrations of amphotericin B were identified with a temporal regression model. Following model fitting, the *−log*_10_ *p*-values (effect of a potential QTL) were calculated across time points (S3 Fig), and significance thresholds were estimated by permutation tests [142]. For nine of the eleven conditions, between one and three QTL (on different chromosomes) were identified across the time course (S4 Fig). Across the combinations of temperature and amphotericin B stress, taking the maximum association at each variable site across the 72-hour time course (Fig 4), a total of thirteen QTL above the thresholds of significance were identified across the eleven temperature and amphotericin B conditions (S5 Fig). Nearly all of the QTL identified showed temporally dependent behavior, with early time-series associations for some QTL and later associations for others (Fig 5A). Taking the maximum association across combinations of temperature and amphotericin B stress, and across time, four unique QTL on chromosomes 2, 11, 12, and 14 were identified (Fig 5B). Two of these QTL, on chromosomes 11 and 12 would not have been detected using the traditional marker-regression framework based on final growth (S6 Fig).

**Fig 4.**
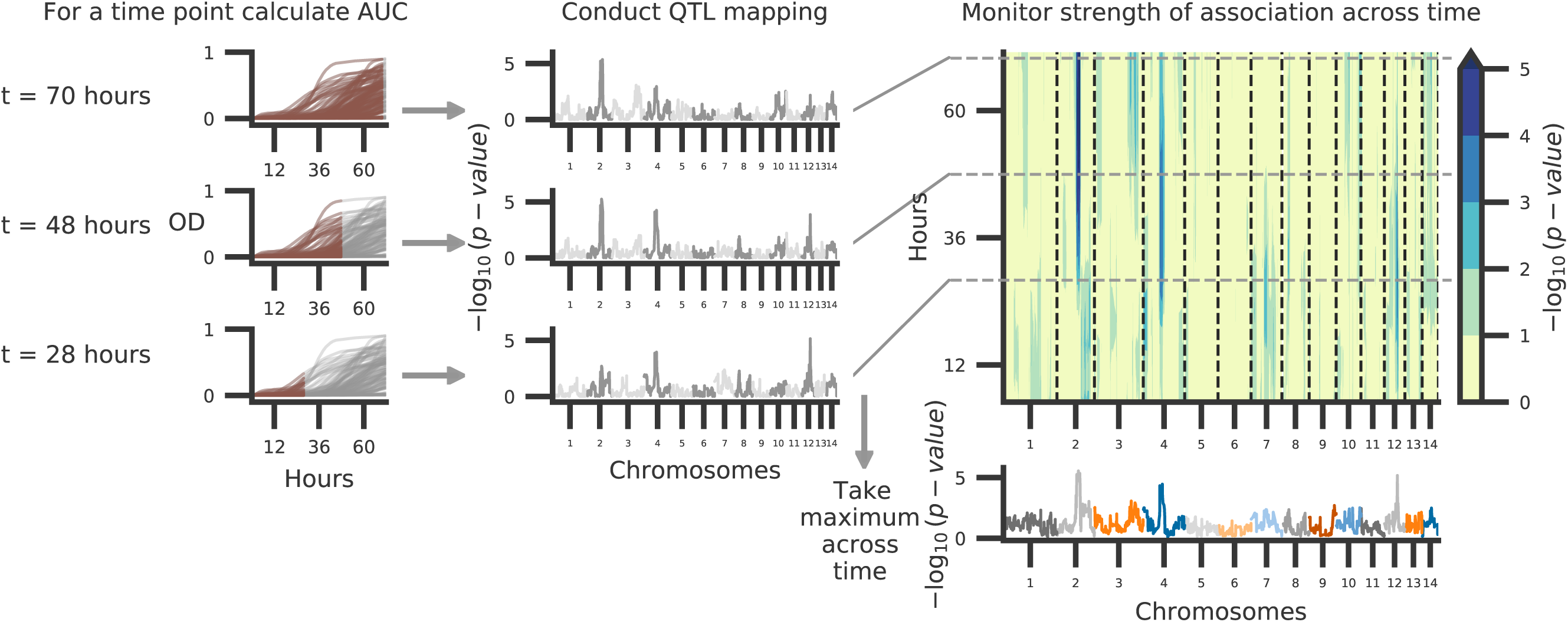
Schematic of temporal QTL mapping. Across experimental conditions, OD was sampled every 15 minutes for 72 hours. Across the 72-hour time course, the median (across replicates) area under the curve (AUC) is calculated per segregant and utilized for QTL mapping, regressing AUC across the 14 chromosomes represented by 3,108 haploblocks. This process is conducted per time point and examples of this analysis at 70, 48, and 28 hours from growth data collected at 37*^◦^*C with 0.125 *µ*g/ml of amphotericin B are depicted. The temporal trends in QTL may then be summarized by taking the maximum per haploblock across time.

**Fig 5.**
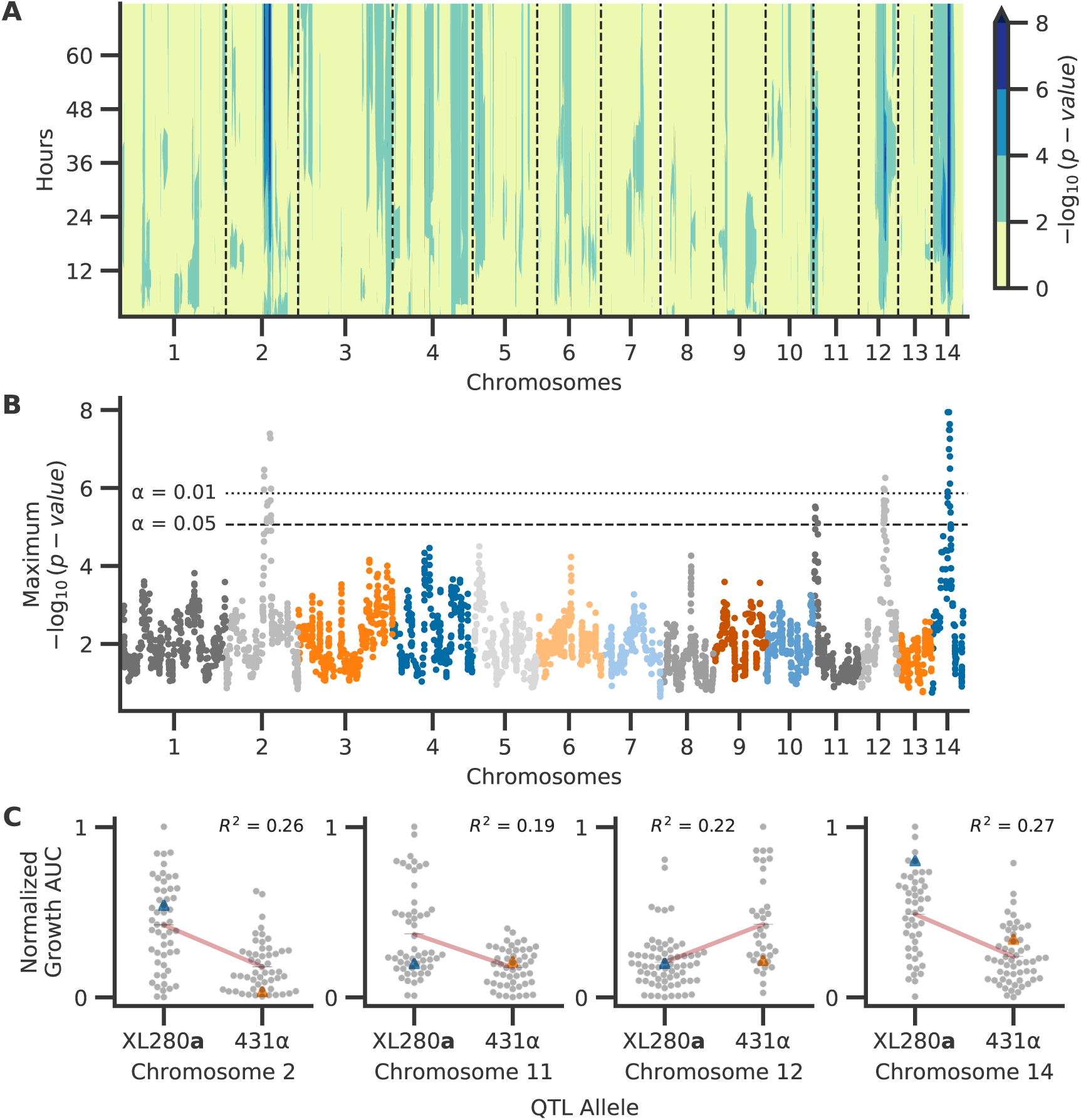
Associations between genotype and phenotype. Across the experimental combinations of temperature and amphotericin B concentrations in Fig 3, median growth AUCs were regressed onto haplotypes for each sample time point in 72-hour time courses. **A**) Temporal analysis of association between genotype and phenotype, collapsed across conditions. Across the experimental conditions, the maximum association across time (y-axis) per chromosome (x-axis) is shown. **B**) QTL collapsed across conditions and time. The x-axis represents chromosomal positions of haploblocks; the y-axis represents the maximum log_10_(p-value) for each haploblock across both time and conditions. The maximum significance thresholds (dotted and dashed horizontal lines) were determined via permutation. **C**) Normalized (max-min normalization) AUC of growth (y-axis) by parental allele (x-axis) at the QTL on chromosomes 2, 11, 12, and 14 (left to right, respectively). Blue and orange triangles represent the AUC values for parental strains, black horizontal lines denote phenotypic means by allele, and red lines indicate the best fit line from the regression used to detect QTL. The heritability of each QTL (annotated in black) is estimated by the coefficient of determination from the regression.

### QTL for thermal tolerance

A QTL on chromosome 14 was identified as having a significant association with growth across the three temperature conditions (30, 37, and 39*^◦^*C) with no drug and at 39*^◦^*C with amphotericin B (S4 Fig). The chromosome 14 QTL was strongest at 39*^◦^*C with no amphotericin B (S5 Fig). This QTL was thus classified as a high temperature growth QTL. At this locus, segregants possessing the XL280**a** haplotype exhibited greater thermal tolerance and outgrew siblings with the 431α haplotype (Fig 5C). This pattern was surprising given that the 431α parental strain is the more thermal tolerant of the parents at 37*^◦^*C. The maximum heritability, as estimated by the coefficient of determination from the linear regression QTL model, was approximately 27%. While this QTL had broad effects across time (Fig 5A, S3 Fig) the maximum association between genotype and phenotype was observed relatively early within the time course at 29 hours.

### A pleiotropic QTL governs melanization, capsule size, and thermal tolerance

QTL for melanization, capsule diameter, and thermal tolerance were mapped to an overlapping region on chromosome 14, suggesting the presence of allelic variation with pleiotropic effects (S7 Fig). Examining the relationships between all of the phenotypes assayed (S8 Fig), growth at 39*^◦^*C is strongly correlated with the capsule size (Spearman *ρ* = 0.43, *p*-value *<* 0.01) and melanization phenotypes (Spearman *ρ* = 0.51, *p*-value *<* 0.01). However, capsule size and melanization phenotypes displayed only a modest correlation (Spearman *ρ* = 0.19, *p*-value = 0.059). Segregants that displayed greater total growth at 39*^◦^*C had larger capsule diameter or lighter colonies but not necessarily both phenotypes (S7 Fig). Because the capsule diameter and melanization phenotypes are strongly correlated with thermal tolerance and because the shared QTL co-localized along chromosome 14, we treated the chromosome 14 locus as a pleiotropic QTL for subsequent analyses.

### Multiple QTL for amphotericin B sensitivity

The QTL on chromosome 2 reached or neared significance in five of the eleven combinations of temperature and amphotericin B concentrations. This QTL was not detected as significant in any of the conditions lacking amphotericin B, and the maximum association between genotype and phenotype was observed at 0.125 *µ*g/ml of amphotericin B (S5 Fig). Temporal analysis indicated that during growth in the presence of 0.125 *µ*g/ml amphotericin B, this QTL reached the threshold of significance in the middle of the 72-hour time course (approximately 36 hours) across multiple temperature conditions and reached its maximum at approximately 65 hours at 30*^◦^*C and 0.125 *µ*g/ml amphotericin B (Fig 5A, S3 Fig). This locus was thus designated as an amphotericin B sensitivity QTL. This QTL explains approximately 26% of the variance in growth at *∼*65 hours at 30*^◦^*C and 0.125 *µ*g/ml amphotericin B. Segregants with the 431α haplotype at this QTL were more susceptible to the fungicidal effects of amphotericin B (Fig 5C).

A second amphotericin B QTL was identified on chromosome 11. This QTL was maximally associated with growth at 37*^◦^*C with 0.175 *µ*g/ml of amphotericin B (S5 Fig). This second QTL explained 19% of the phenotypic variation as estimated by the regression model. At this locus, segregants with the XL280**a** haplotype outgrew their sibling progeny with the 431α haplotype (Fig 5C). The effect of this QTL was seen in the first two-thirds of the 72-hour time course, reaching a maximum at *∼*40 hours and trailing off thereafter (S3 Fig).

The QTL identified on chromosome 12 surpassed the significance threshold in three conditions of high temperature (37*^◦^* and 39*^◦^*C) and high amphotericin B concentration (0.125 and 0.175 *µ*g/ml, S5 Fig). This QTL was designated as a drug associated QTL as it only appeared significant in conditions with amphotericin B concentrations larger than 0.125 *µ*g/ml. At this QTL, segregants with the parental 431α allele outgrew progeny with the XL280**a** allele. Furthermore, of the QTL identified here, this was the only QTL that displayed a positive association with alleles from the 431α background (Fig 5C). At the highest concentration of amphotericin B, this QTL was maximal near the middle of the time course (*∼*36 hours) (Fig 5A, S3 Fig). The phenotypic heritability explained by this locus was estimated to be *∼*22%.

### Identifying candidate genes and nucleotide variants

For the four QTL detected in temperature and amphotericin B experiments, the regions containing candidate genes were determined by taking the maximum association for each haploblock across time, temperature, and amphotericin B concentration and calculating the left and right boundaries of haploblocks above the maximum significance threshold (across conditions). The open reading frames of genes within these regions were determined by realigning gene sequences from the JEC21α reference annotation to the XL280α reference. For all genes within the four candidate regions, we predicted potential changes in protein sequence due to the genetic variants between the XL280**a** and 431α parental strain (S9 Fig) and identified those with non-synonymous changes (S3 Table). Orthologous genes in the *C. neoformans* background were identified for genes with non-synonymous changes between the parental strains. Where available, gene deletion strains [21] were used in follow up temperature and amphotericin B growth assays (S10 – S13 Figs). Candidate genes and causal genetic variants were further narrowed down by consulting the previous literature, considering the severity of the non-synonymous genetic changes on protein length and function, and comparing the growth curve profiles from temperature and amphotericin B experiments on *C. neoformans* deletion strains. Using this approach, we identified candidate quantitative trait genes (QTGs) and their associate quantitative trait nucleotides (QTNs) for three of the four QTL identified above (discussed below). For the chromosome 11 QTL we were unable to predict a candidate QTG as several significant non-synonmous changes are observed in genes with unknown function and none of the phenotypes of *C. neoformans* deletion mutants for this region were consistent with our QTL mapping results. Although our analyses focused on coding variants within each QTL region, a summary of non-coding and synonymous variants is included in S3 Table.

### *RIC8* is a candidate QTG for the pleiotropic QTL on chromosome 14

The pleiotropic chromosome 14 QTL contributing to variation in melanization, capsule diameter, and high temperature growth spanned approximately 69 kb and was located between the coordinates 354,000 to 423,000 bp. There are 29 genes within this QTL region, 17 of which are estimated to have genetic variants that lead to non-synonymous changes between the parental backgrounds (S3 Table). A single-nucleotide polymorphism (SNP) identified in the second to last exon of the gene *RIC8* (*CNN01270*) is predicted to cause a premature stop-gain in the XL280**a** background when compared to the JEC21α reference strain (Fig 6). Additional non-synonymous changes in the *RIC8* gene were identified in the 431α strain compared to both the XL280**a** parental strain and the JEC21α reference strain and include an in-frame codon deletion and a predicted shift in the stop codon (S14 Fig).

**Fig 6.**
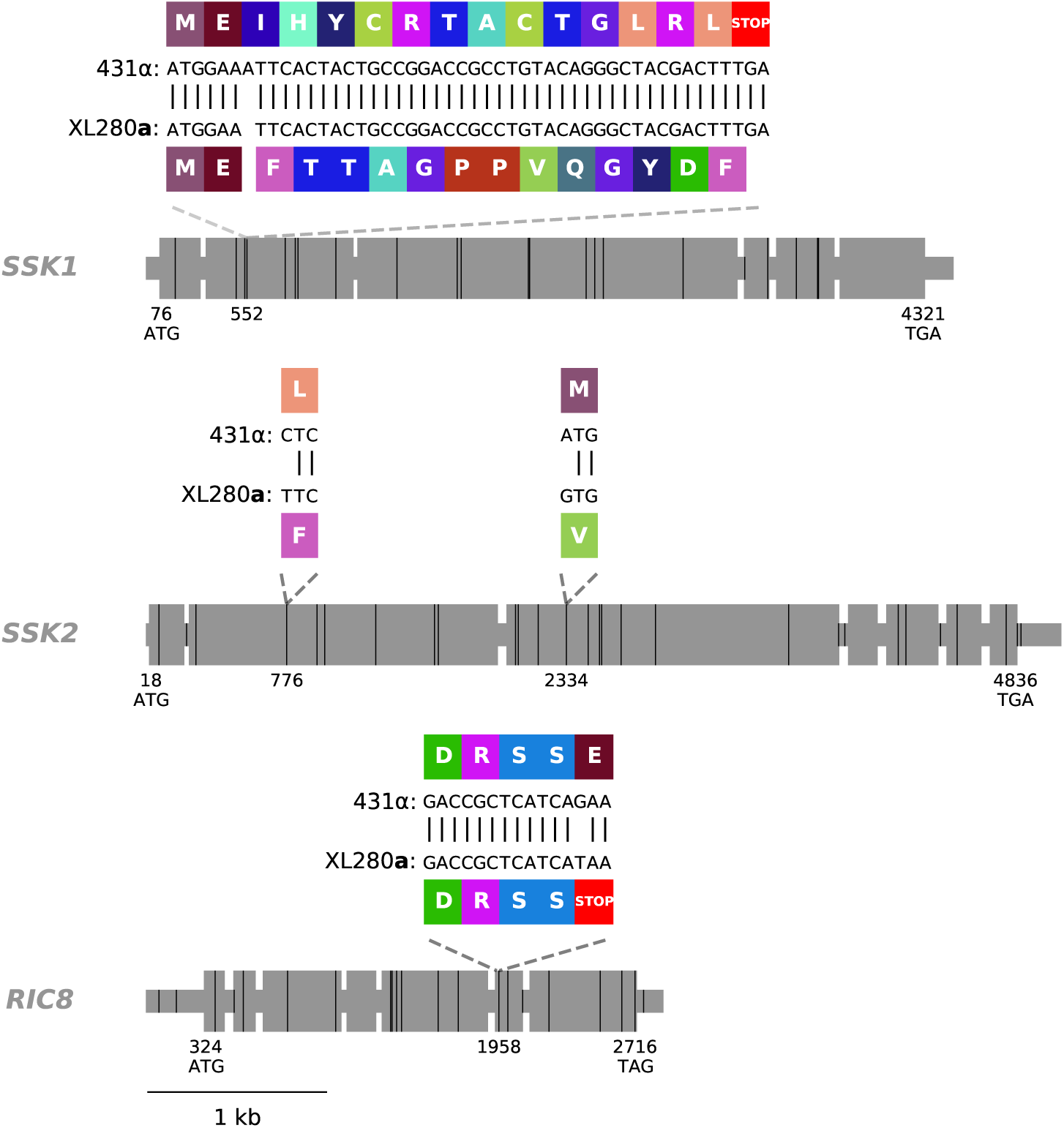
*SSK1*, *SSK2*, and *RIC8* gene models. Exons are shown as large grey rectangles, while the introns, 5’ UTR, and 3’ UTR are shown as grey, horizontal lines. The positions of the predicted start and stop codons are annotated along the bottom of the gene bodies and the positions of genetic differences between 431α and XL280**a** are marked by black, vertical lines. Within the second exon of *SSK1*, an insertion of a single nucleotide, present in the 431α parental strain is predicted to cause a frame shift that leads to a downstream early stop-gain. Within the second and third exons of *SSK2*, two SNPs are annotated that lead to non-synonymous changes previously identified by Bahn et al. [143]. Within the second-to-last exon of *RIC8*, a SNP is present in the XL280**a** parental strain that is predicted to cause a premature stop. The local, predicted translations of the regions near these non-synonymous, genetic variants and associated amino acids are annotated in colored rectangles.

Ric8 is a guanine nucleotide exchange factor for Gpa1, the G*_α_* activator of the cAMP-PKA pathway in *Cryptococcus* [144]. In *C. neoformans*, *ric8*Δ strains have been previously demonstrated to exhibit melanization and capsule defects [144]. We confirmed the melanization defect using a *ric8*Δ strain from the *C. neoformans* deletion collection (Fig 7A, S15 Fig). To test the effect of *ric8* mutations on thermal tolerance, growth of the *ric8*Δ strain was profiled at 37*^◦^* and 39*^◦^*C. At these elevated temperatures, the *ric8*Δ strain exhibited a slower initial growth rate than the wild-type control strain, but then reached a higher maximum density, with the result being higher total growth (Fig 7B, S15 Fig).

**Fig 7.**
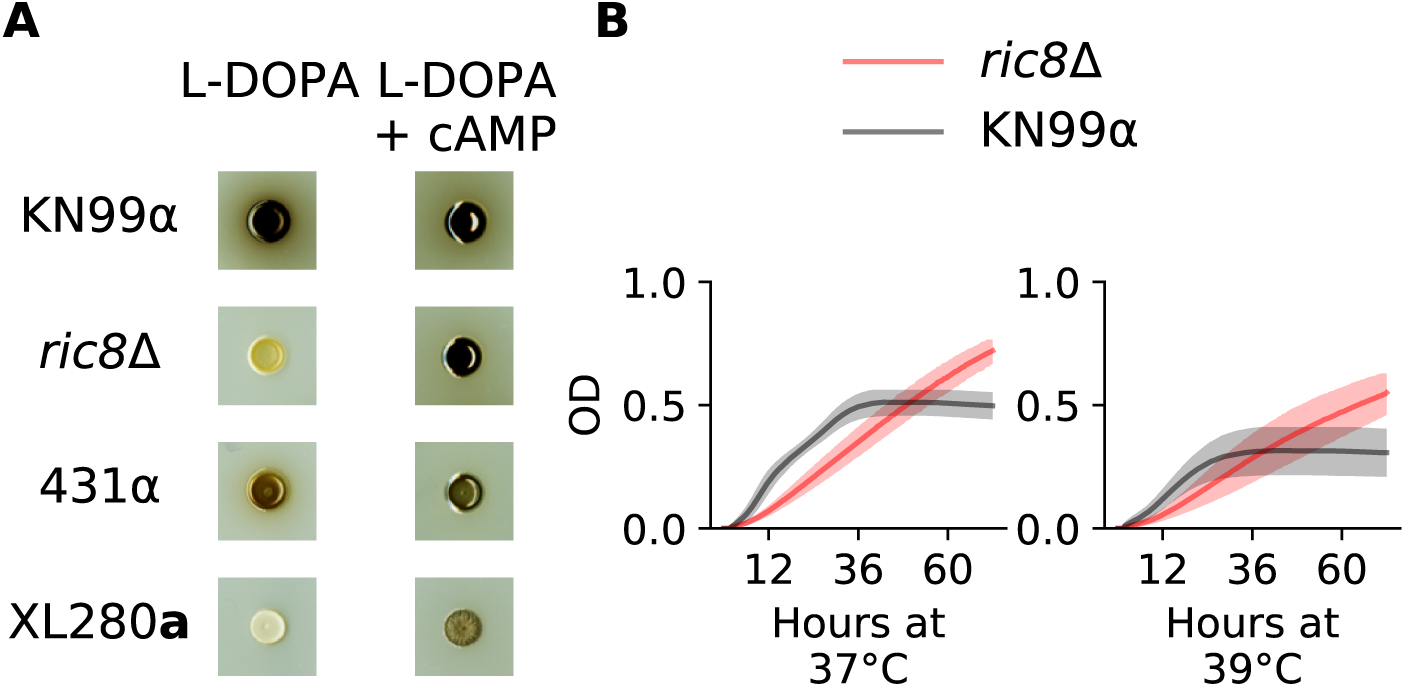
Melanin and high-temperature phenotypes of *RIC8*. **A**) The *C. neoformans* strain, KN99α, the *ric8*Δ deletion strain, and *C. deneoformans* parental strains, 431α and XL280**a** were grown on plates with L-DOPA and L-DOPA + cAMP. Both the *ric8*Δ strain and XL280**a** demonstrated large increases in the production of melanin when grown in the presence of exogenous cAMP. **B**) Growth in liquid culture (OD_595_, y-axis) of KN99α (black) and the corresponding *ric8*Δ strain (red) under conditions of heat stress (37*^◦^*C and 39*^◦^*C) across 72 hours (x-axis).

Ric8 loss-of-function mutants are predicted to have lower levels of cAMP signaling [144]. Consistent with the finding of Gong et al. [144], the addition of exogenous cAMP to L-DOPA plates restored melanization in the *C. neoformans ric8*Δ strain. The parental strain XL280**a**, bearing the predicted *ric8* loss-of-function allele, also exhibited increased melanization when grown on plates with L-DOPA + cAMP (Fig 7A). The 431α parent exhibited only modest changes in melanization in the presence of cAMP, suggesting that cAMP-PKA signaling is already active in this background. Because the melanization and thermal tolerance phenotypes of the *ric8*Δ strain were consistent with the effects predicted from the QTL mapping, as were the predicted effects of chemical manipulation of the XL280**a** background, the *RIC8* allele identified in the XL280**a** background (*RIC8* ^XL280**a**^) was labeled as a likely QTN for melanization, high temperature growth, and capsule size.

### *SSK1* is a candidate QTG for amphotericin B sensitivity

The QTL peak on chromosome 2 spanned approximately 154-kb and was located between coordinates 847,000 and 1,001,000 bp. There are 43 genes within this peak, and 18 of these genes were predicted to have non-synonymous changes between the parental strains (S3 Table). Of these 18 genes, *SSK1* (*CNB03090*) exhibits the most dramatic difference between the two parental strains. The 431α parental haplotype includes a single base-pair insertion within the second exon that is predicted to cause a frame shift, leading to a premature stop-gain (Fig 6). Because this stop-gain was predicted to truncate more than three-quarters of the Ssk1 protein sequence, the *SSK1* ^431α^ variant was categorized as a likely loss-of-function allele.

To provide an independent test of the phenotypic effect of *SSK1* loss-of-function mutations, we phenotyped the *ssk1*Δ strain from the *C. neoformans* gene deletion collection [21]. The *ssk1*Δ strain in the H99α *C. neoformans* strain background exhibited an amphotericin B sensitive phenotype, consistent with the phenotype of segregants bearing the *SSK1*^431α^ predicted loss-of-function allele (S16 Fig). Additional *ssk1*Δ strains were constructed in the *C. deneoformans*, XL280**a** and 431α parental backgrounds (S1 Table) and phenotyped for amphotericin B sensitivity (S16 Fig). In the 431α strain background, the *ssk1*Δ knockout strain exhibited an amphotericin B sensitive phenotype, as expected. However, relative to the wild type XL280**a** strain, none of the XL280**a** *ssk1*Δ strains exhibited an amphotericin B sensitive phenotype. We hypothesized this may be due to additional undiscovered allelic variants in this background that also contribute to amphotericin B resistance.

### The centromere hinders fine mapping of chromosome 2 QTL

A fine-mapping procedure was conducted to narrow down the QTL peak on chromosome 2. Specifically, intergenic regions were identified that flank the QTL on chromosome 2 and within these regions, *NAT* and *NEO* markers were transformed into the XL280**a** and 431α parental strains (respectively). From this procedure, one and three transformants were generated in the XL280**a** and 431α parental strain backgrounds, respectively (S1 Table). Three **a**–α bisexual crosses were conducted using these marked parental strains, and a large pool of segregants was generated using a random sporulation protocol. From this pool of segregants 173 *NAT^R^ NEO^R^* segregants, with recombination events within the QTL on chromosome 2 between the two flanking markers, were selected.

Examining the allele frequencies of these progeny, a bias in the *SSK1* allele was observed – only 10% of the population possessed the *SSK1* allele from the 431α parental strain. This was disappointing given the limits on statistical power needed for additional QTL mapping. In this species, centromeres are flanked by crossover hot- and cold-spots [141]. We hypothesized that the proximity of the *SSK1* locus to the centromere on chromosome 2 led to a repression of recombination near the left flanking *NAT* marker, leading to the deviation from the expected 50:50% allele frequencies (S17 Fig).

### *SSK2* is also a candidate QTG for amphotericin B sensitivity

The chromosome 12 QTL spans *∼*62 kb and is centered between coordinates 554,000 and 616,000 bp. There are 25 genes within this region, 15 of which are predicted to contain non-synonymous changes between the parental strains. Two genes within this region contain a stoploss and stop-gain, but are hypothetical and of unknown function (S3 Table). Furthermore, deletion strains of these unknown genes in the H99α strain background did not display an amphotericin B sensitive phenotype (S12 Fig).

Among the other candidate genes within this QTL is *SSK2* (*CNL05560*), a MAP kinase of the HOG pathway [143]. By comparing the *SSK2* genotypes of the XL280**a** and 431α parent strains, three SNPs were identified that are predicted to cause non-synonymous amino acid differences between the two parental backgrounds (S14 Fig). Two of these non-synonymous SNPs and their associated amino acid changes were previously identified by Bahn et al.[143] and shown to underlie differences in high temperature growth, fludioxonil sensitivity, and osmotic stress responses of *C. deneoformans* strains (Fig 6).

### QTL mapping of HOG-related phenotypes

Our initial studies of amphotericin B susceptibility implicated two key genes – *SSK1* and *SSK2* in the HOG pathway, a signaling network that plays a central role in the regulation of cellular responses to osmostress in fungi. Consequently, we predicted that segregants in this study might show variation for additional HOG pathway related phenotypes attributable to one or both of these loci. Thus, we undertook additional analyses of HOG-related phenotypes including resistance to salt stress, resistance to the antifungal drug fludioxonil, and oxidative stress tolerance

### QTL for osmotic stress response

The segregants from this cross were assayed for variation of growth in response to osmotic stress. High resolution images of colonies grown with 1M NaCl were made using transmissive imaging (S18 Fig) and the mean grayscale intensity of each colony was used as a proxy for population density. The two parental strains did not vary greatly in their response to salt stress but there was significant growth variation among the segregants (S18 Fig).

A salt tolerance QTL, explaining *∼*18% of the phenotypic variance, was identified on chromosome 10 (S18 Fig). Segregants with the XL280**a** allele at the peak of this QTL outgrew sibling segregants with the 431α allele (S18 Fig). While there are 17 genes within this QTL, none were identifiable as obvious candidate genes for follow up experimentation (S3 Table).

### Fludioxonil resistance is governed by an epistatic interaction between *SSK1* and *SSK2*

Fludioxonil is an agricultural antifungal drug whose mode of action is thought to be hyper-activation of the HOG pathway, leading to physiological effects such as glycerol accumulation and increased turgor pressure [145]. Resistance to fludioxonil has been shown to occur primarily through mutations that ameliorate or decrease HOG signaling. While resistance to fludioxonil is rare in most fungal species due to the negative pleiotropic consequences of HOG pathway loss-of-function mutations [145], *Cryptococcus* is unusual in that many strains of both *C. deneoformans* and *C. neoformans* exhibit resistance to this drug. Bahn et al.[143] demonstrated that variation in sensitivity to fludioxonil among *Cryptococcus* lineages correlates with Hog1 phosphorylation levels which are in turn correlated with two different allelic states observed at *SSK2*. The allelic states identified by Bahn et al. are the same *SSK2* alleles identified as segregating in the cross considered here, with the *SSK2* ^XL280**a**^ allele predicted to correlate with resistance to fludioxonil and the *SSK2* ^431α^ variant predicted to be sensitive.

Surprisingly, when exposed to fludioxonil (100 *µ*g/ml) the two parental strains both exhibited resistance. We reasoned that the resistance seen in the 431α parental strain was due to an epistatic interaction involving the *SSK1* loss-of-function allele identified in this background. Following this logic, we predicted that recombinant segregants with the *SSK1* ^XL280**a**^ *SSK2* ^431α^ genotype would exhibit sensitivity to fludioxonil. Consistent with this prediction, 14 of the 20 segregants with the *SSK1* ^XL280**a**^ *SSK2* ^431α^ genotype were fludioxonil sensitive. Segregants with any of the other of three possible allele combinations at these two loci were fludioxonil resistant (Fig 8).

**Fig 8.**
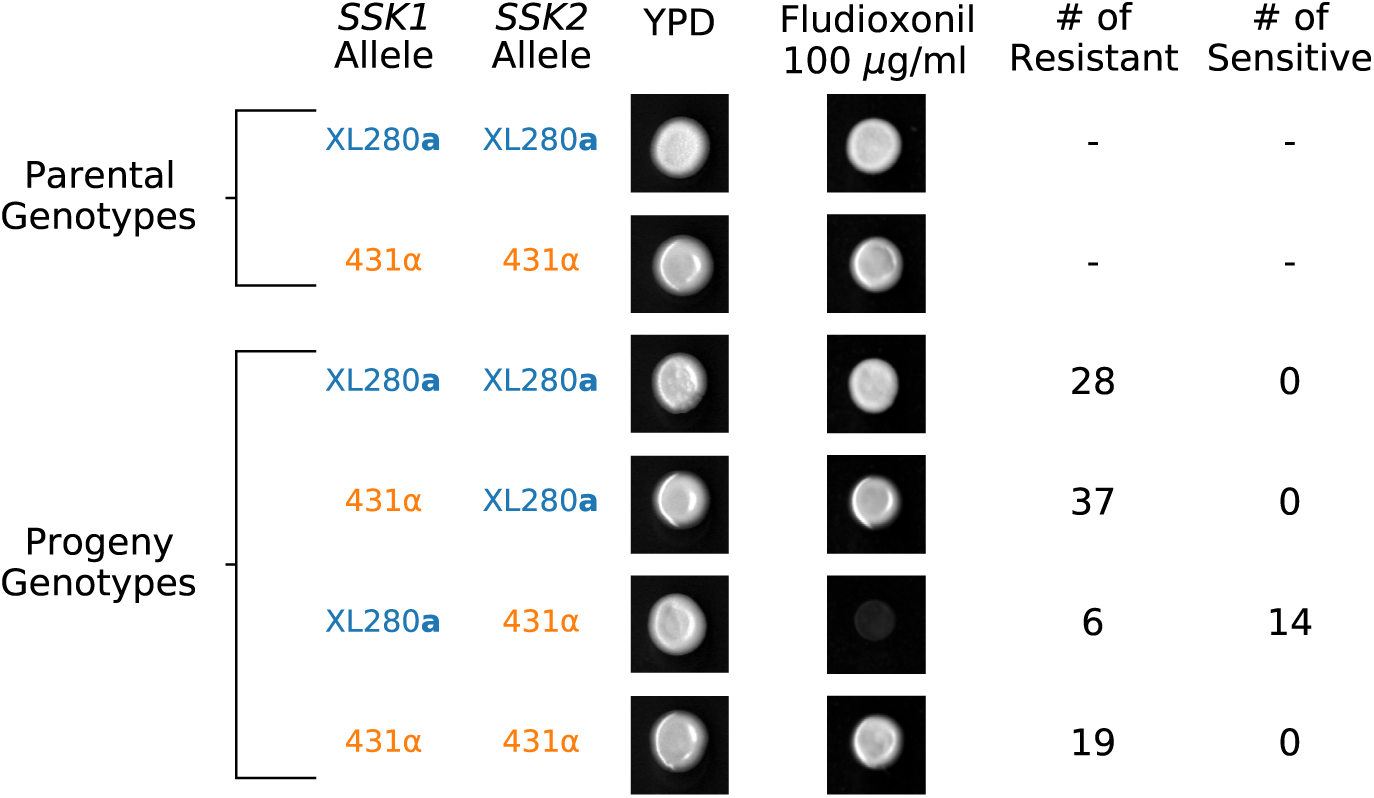
*SSK1* and *SSK2* govern fludioxonil sensitivity. The relationship between *SSK1* and *SSK2* genotypes and fludioxonil sensitivity. Only progeny with the *SSK1*^XL280**a**^ *SSK2*^431α^ genotype are sensitive to fludioxonil. Resistant *SSK1*^XL280**a**^ *SSK2* ^431α^ segregants suggest additional higherorder epistatic interaction involving *SSK1* and *SSK2*.

To provide further evidence for an epistatic interaction between *SSK1* and *SSK2* we assayed the additional set of fine-mapped offspring from **a**–α bisexual crosses between the XL280**a** strain and the three 431α strains (S1 Table) for fludioxonil resistance. In this larger set of progeny, only those segregants that were the genetic mosaics of the XL280**a** and 431α strains, possessing the *SSK1* ^XL280**a**^ *SSK2* ^431α^ genotype, displayed sensitivity to fludioxonil (S19 Fig). These data supported our hypothesis that the *SSK1* ^431α^ allele observed in the 431α parental strain is indeed a naturally occurring loss-of-function mutation and in this isolate the *SSK1*^431α^ has an epistatic effect with *SSK2* ^431α^, rescuing an otherwise fludioxonil-sensitive *SSK2* phenotype. Our findings also point to even higher order genetic interactions – a small number of segregants among those with the mosaic *SSK1* ^XL280**a**^ *SSK2* ^431α^ genotype were resistant, indicating the presence of additional loci that interact epistatically with *SSK1* and *SSK2* to mediate fludioxonil resistance.

Of the multiple traits examined in this study, fludioxonil sensitivity was the only phenotype to exhibit distinct distributions between segregants derived from α–α unisexual versus **a**–α bisexual matings. In the primary mapping population, the majority of fludioxonil sensitive progeny were derived from the unisexual cross. This is due to an allelic bias at the *SSK2* locus, wherein bisexually derived segregants preferentially inherit the *SSK2* ^XL280**a**^ allele. However, in the fine-mapped offspring, all of which are derived from **a**–α bisexual matings, there is a bias *in favor* of the *SSK2* ^431α^ allele. Despite these opposing allelic biases, for both unisexually and bisexually derived offspring, the only segregants exhibiting fludioxonil sensitivity are those with the allelic combination *_SSK1_* XL280**a** *_SSK2_* 431α.

### *SSK2* and *RIC8* are QTGs underlying oxidative stress tolerance

Resistance to oxidative stress is another virulence related trait in *Cryptococcus*, that is associated with HOG signaling [143]. Segregants were grown on media containing 5 mM of H_2_O_2_, and colony growth was quantified from high-resolution images by two independent observers using ordinal scoring (Fig 9A). There was significant variation in response to H_2_O_2_ across the segregants. The XL280**a** parental strain displayed higher tolerance of H_2_O_2_ (on average) than the 431α parental strain, and a portion of the progeny (less than 25%) displayed no growth and complete sensitivity to H_2_O_2_ (Fig 9B).

**Fig 9.**
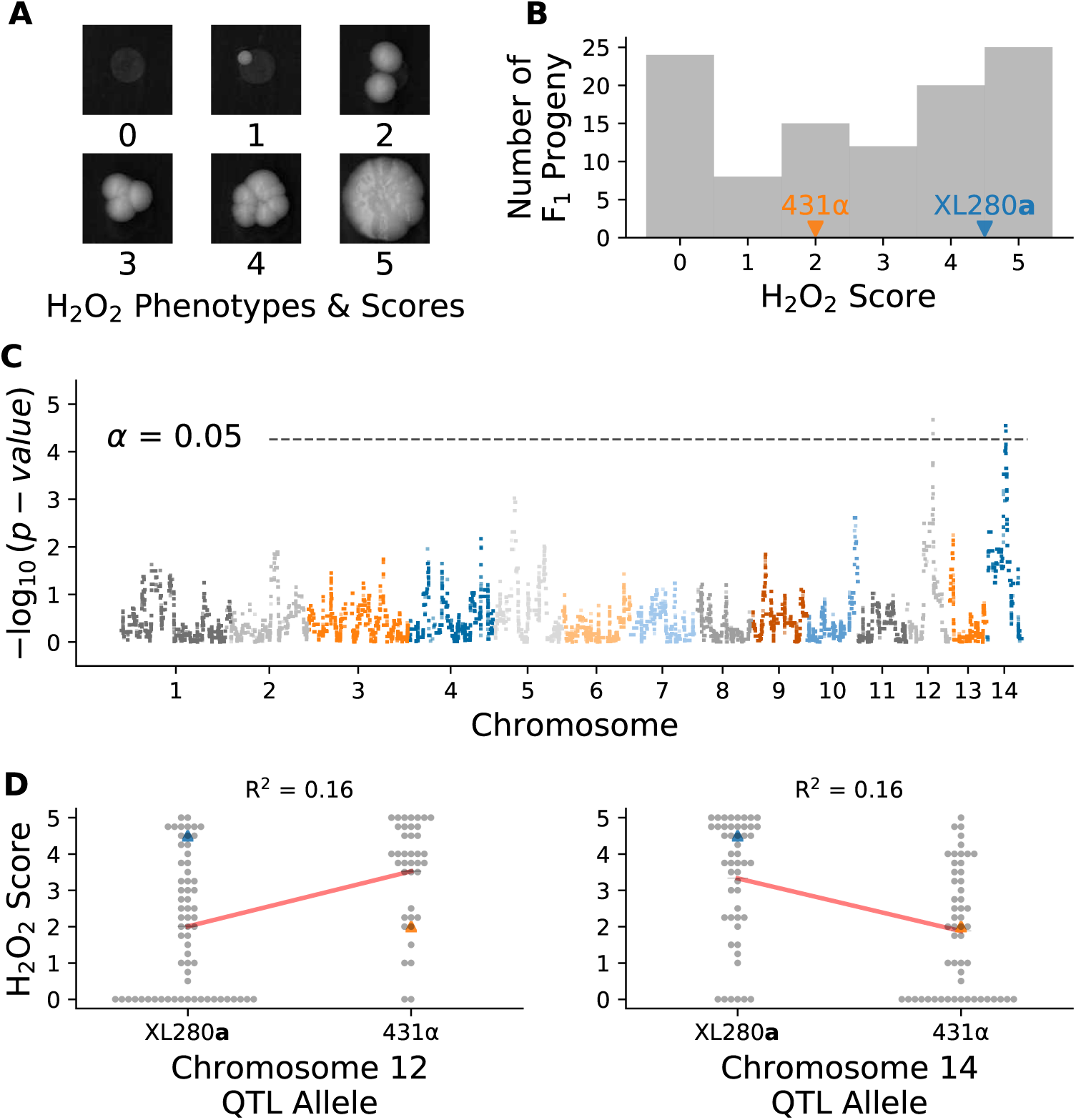
Variation in hydrogen peroxide sensitivity. QTL analysis of variation in response to hydrogen peroxide. **A**) Example growth phenotypes and scores of *C. deneoformans* segregants grown on media with hydrogen peroxide. **B**) Histogram of hydrogen peroxide growth scores. Growth scores of parental strains are marked by vertical lines. **C**) Manhattan plot of the association between genotype and growth in response to hydrogen peroxide. The x-axis represents chromosomal locations of haploblocks and the y-axis represents the strength in association between genotype and variation in growth score. **D**) Hydrogen peroxide growth scores (y-axis) as a function of allele at the peak of chromosome 12 (left) and chromosome 14 (right) QTL. Blue and orange triangles mark the parental phenotypes, black horizontal lines denote the average phenotypes of segregants by allele, and red lines are regression models relating the genotype to phenotype. The heritablity at each locus is estimated from these models and annotated in black.

Two QTL associated with hydrogen peroxide growth were detected (Fig 9C), on chromosomes 12 and chromosome 14, each with modest heritablity (*∼*16% at each QTL); (Fig 9D). These two QTL overlapped with the previously identified QTL governing amphotericin B sensitivity on chromosome 12 and the pleiotropic QTL underlying variation in melanization, capsule size, and thermal tolerance on chromosome 14. Joint analysis of the H_2_O_2_ growth scores and growth at 37*^◦^*C with *µ*g/ml of amphotericin B revealed a strong correlation between these phenotypes (Spearman *ρ* = 0.5, *p*-value < 7^-8^). Segregants with the 431α allele at the chromosome 12 QTL peak display greater tolerance to H_2_O_2_ and resistance to amphotericin B (S20 Fig). Similarly, for the virulence-related phenotypes associated with the chromosome 14 QTL, the H_2_O_2_ phenotype was strongly correlated with thermal tolerance at 39*^◦^*C (Spearman *ρ* = 0.53, *p*-value < 9^-9^) and negatively correlated with melanization (Spearman *ρ* = −0.59, *p*-value < 5^-11^). There was no significant correlation between H_2_O_2_ growth and the capsule phenotype (Spearman *ρ* = 0.15, *p*-value = 0.13). Given the location of H_2_O_2_ QTLs, and the general trends of phenotypic correlations observed, we predict that *SSK2* (chromosome 12) and *RIC8* (chromosome 14) are the underlying QTGs for this trait.

At the predicted QTN for *SSK2* and *RIC8* the marginal variance explained was approximately 13% and 14% (respectively). A linear model based solely on additive effects of these two loci explains approximately 21% of phenotypic variance, while models that include an interaction term between these loci explains 26.4% of the variance (ANOVA, *p*-value *<* 1.0*^−^*^6^). Segregants with the *SSK2* ^XL280**a**^ *RIC8*^431α^ genotype displayed the greatest average sensitivity to H_2_O_2_ while other allelic combinations of *SSK2* and *RIC8* exhibited similar average H_2_O_2_ resistance (S21 Fig).

Re-examination of thermal tolerance phenotypes (Fig 3) with respect to two-locus *SSK2 RIC8* genotypes, suggests that epistasis between these two loci may also influence this trait. Across the temperatures used in growth curve assays (30, 37, 39*^◦^*C), segregants bearing the *SSK2* ^XL280**a**^ *RIC8* ^431α^ genotype displayed the poorest overall growth (S21 Fig). This growth defect was most pronounced at 39*^◦^*C (ANOVA, *R*^2^ = 0.332, *p*-value *<* 9*^−^*^9^).

### Three-way epistasis contributes to hydrogen peroxide resistance

Since *SSK2* was found to interact epistatically with both *SSK1* (fludioxonil resistance) and *RIC8* (oxidative stress resistance and thermal tolerance), we hypothesized that higher-order interactions involving all three loci might contribute to phenotypic variation in one or more of these traits. To test for three-way epistasis, we employed an approach proposed by Hu et al.[146], which uses a statistic called “information gain” (IG), which is based on information-theoretic mutual information measures [147]. Hu et al.’s IG statistic provides a measure of synergistic interaction between three loci with respect to a phenotype of interest, after subtracting the information inherent in single locus effects and synergies between pairs of loci. Since the IG statistic requires discrete data, we limited our analysis of three-way epistasis to a transformed H_2_O_2_ resistance phenotype, classifying each segregant as either sensitive, intermediate, or resistant. Applying the IG method to H_2_O_2_ resistance, we find evidence for single locus effects (IG(*SSK1*) = 4.6%, *p*-value = 0.005; IG(*SSK2*) = 6.0%, *p*-value = 0.002; IG(*RIC8*) = 5.4%, *p*-value = 0.006) as well as a three-way synergy between *SSK1*, *SSK2*, and *RIC8* (IG(*SSK1*,*SSK2*,*RIC8*) = 7.0%, *p*-value = 0.004), but no significant pairwise synergies.

Fig 10A illustrates the distributions of H_2_O_2_ growth scores for each of the eight possible genotypic combinations of *SSK1*, *SSK2*, and *RIC8*. The mapping between the three-locus genotypes and H_2_O_2_ resistance can be summarized as follows. Segregants with the two-locus genotypic combination *SSK2* ^XL280**a**^ *RIC8* ^431α^ exhibit the lowest average H_2_O_2_ resistance. Conversely, segregants with the opposite genotypic combination *SSK2* ^431α^ *RIC8* ^XL280**a**^ exhibit high average H_2_O_2_ resistance. The phenotype of segregants with the other two-locus combinations of *SSK2* and *RIC8* (i.e. *SSK2* ^XL280**a**^ *RIC8* ^XL280**a**^ and *SSK2* ^431α^ *RIC8* ^431α^) depends on their genotypic state at *SSK1* – those with the *SSK1*^XL280**a**^ allele have high average resistance, while those with the *SSK1*^431α^ allele exhibit intermediate average resistance.

**Fig 10.**
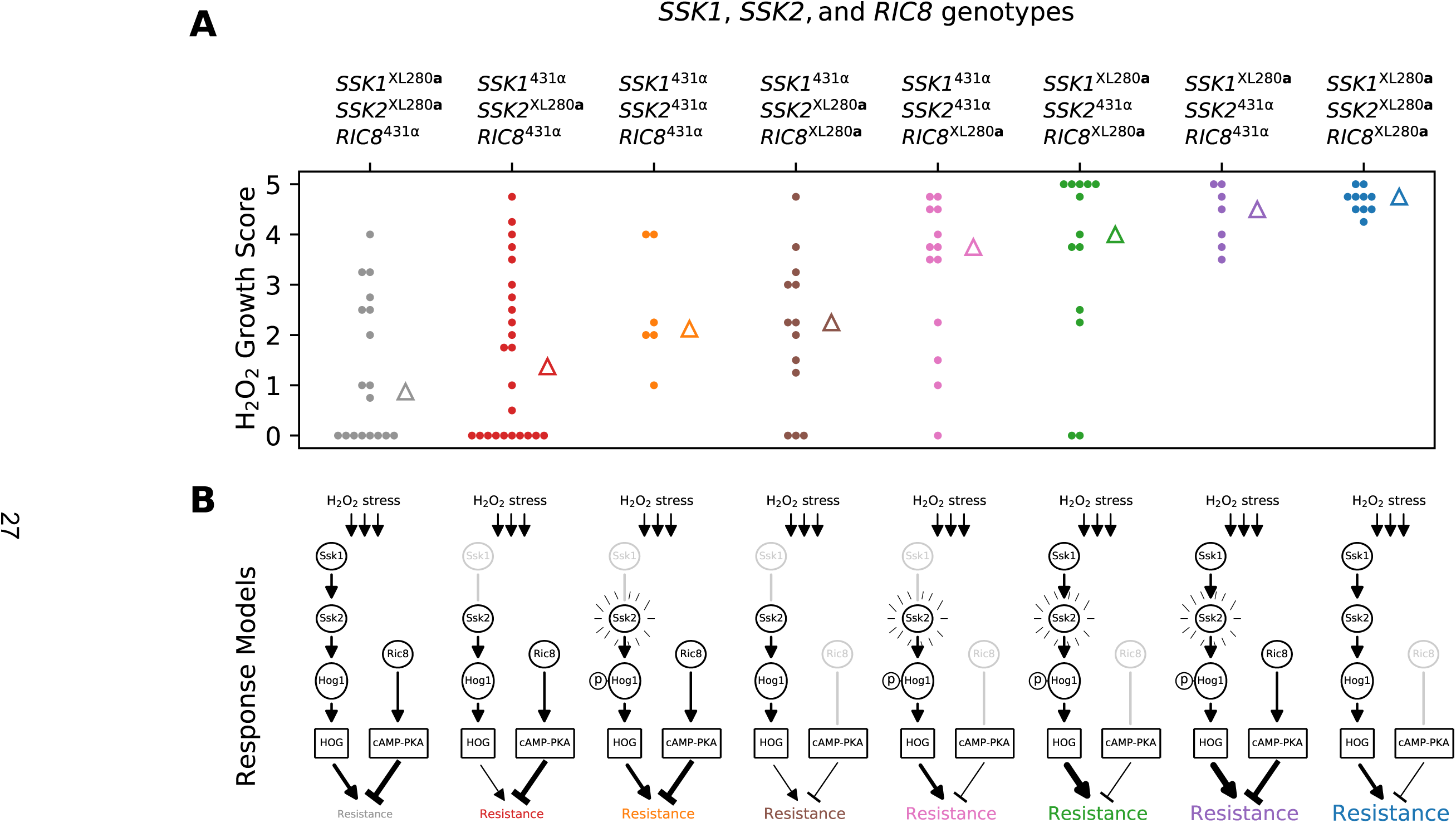
Three-way epistasis underlies H_2_O_2_ resistance. Three-way epistasis between *SSK1*, *SSK2*, and *RIC8* governs resistance to H_2_O_2_ stress. **A**) Distributions of growth scores under H_2_O_2_ stress (y-axis) for segregants with different three locus genotypes at *SSK1*, *SSK2*, and *RIC8* (x-axis and color). Triangles mark the median scores per genotype. **B**) Proposed models for HOG and cAMP-PKA signaling for each genotypic combination in **A**. The *SSK1* ^431α^ and *RIC8* ^XL280**a**^ alleles are predicted to be non-functional and associated with reduced signaling. The *SSK2* ^431α^ allele has been previously associated with increased basal levels of Hog1 phosphorylation. Proposed levels of HOG and cAMP-PKA signaling are denoted by edge thickness of expression and repression arrows.

### The *RIC8*^XL280a^ allele likely arose during laboratory passage

Allelic variation at the *RIC8* gene, which is predicted to affect cAMP-PKA signaling, was associated with phenotypic variation for a large number of virulence traits. Given its prevalence across our mapping experiments, we sought to identify the ancestral source of the premature stop-gain allele observed in the XL280 backgrounds. The strains XL280**a** and XL280αSS are laboratory generated strains [137], and analysis of ancestral, progenitor strains used in their construction revealed that the *RIC8* stop-gain allele is also found within the strain B3502 [148]. However, this allele was not present in either progenitor strains used to construct B3502. We therefore concluded that the premature stop-gain in the *RIC8*^XL280**a**^ allele is inherited from the strain B3502 and is a *de novo* mutation generated during its construction. The findings of prior studies that have used the B3502 background [e.g. 52, 70, 143, 149–151] should be evaluated in light of potential abrogation of cAMP-PKA signaling.

## Discussion

The work presented here is the highest resolution QTL mapping study to date in a human fungal pathogen [79, 110, 111]. We mapped QTL for multiple virulence-associated traits as well as resistance to two widely used antifungal drugs. By exploiting the detailed SNP information that whole-genome sequence data provides, we subsequently identified specific genes (Quantitative Trait Gene; QTG) and nucleotide variants (Quantitative Trait Nucleotide; QTN) that are likely to underlie those QTL. Of particular note is the fact that the three QTG we identified are regulators of signaling pathways – the cAMP-PKA pathway and the HOG pathway – that are important for fungal adaptation to extracellular stresses [152, 153] and have been shown to be integral to virulence in *Cryptococcus* [24, 154]. Both of these pathways regulate multiple physiological and morphological traits in *Cryptococcus* as well as other fungi including *S. cerevisiae* [155], *Candida albicans* [26, 156], and *Candida auris* [157].

### Allelic variation in cAMP-PKA signaling has highly pleiotropic effects on virulence traits

QTL for melanization, thermal tolerance, capsule size, and growth under oxidative stress (H_2_O_2_) all mapped to the same approximate region on chromosome 14. We identified a likely QTN for this pleiotropic QTL – a SNP that leads to a premature stop-gain within the penultimate exon of the gene *RIC8*. This genetic variant is present in the XL280**a** parental strain. Ric8 is a guanine nucleotide exchange factor for the G*_α_* protein Gpa1, which activates the cAMP-PKA signaling pathway in *C. neoformans* [144, 158]. Based on the location of this variant, we predict that the XL280**a** allele results in a loss-of-function, perhaps partial, of *RIC8*, and hence reduces signalling through the cAMP-PKA pathway.

In our mapping population the *RIC8* ^XL280**a**^ allele was associated with decreased melanisation but increased thermal tolerance, capsule size, and H_2_O_2_ resistance. The associations we observed between phenotypes and *RIC8* genotypes in our *C. deneoformans* mapping population show a mixture of agreement and disagreement with prior studies of cAMP-PKA signaling in *Cryptococcus*, most of which have been conducted in *C. neoformans*. For example, Gong et al. [144] showed that in *C. neoformans*, *ric8*Δ mutants exhibit a loss of melanization and reduced capsule size, and that both of these phenotypes could be rescued by cAMP supplementation. We found that addition of exogenous cAMP to the growth medium increased melanization of the XL280**a** strain, consistent with the prediction that the *RIC8* ^XL280**a**^ is associated with reduced cAMP signaling. However, contrary to that prior study, the predicted loss-of-function *RIC8* ^XL280**a**^ allele was associated with *increased* relative capsule size.

Gong et al. [144] did not examine thermal tolerance in their study, but we assayed growth of a *C. neoformans ric8*Δ strain at high temperatures, and found that this strain exhibited slower initial growth rates relative to the wild-type background, but came to a higher overall population density. This parallels the thermal tolerance phenotype we observed for the XL280**a** background. A similar inverse relationship between thermal tolerance and cAMP-PKA signaling has been observed in *S. cerevisiae*; Li et al. [159] found that hyperactivation of cAMP-PKA signaling reduces resistance to acute heat stress while PKA inhibition increased resistance.

Considering the multivariate relationships among these four traits, we find that: a) melanisation is negatively correlated with thermal tolerance and H_2_O_2_ resistance; b) capsule size is positively correlated with thermal tolerance but weakly correlated with melanization and H_2_O_2_ resistance; and c) H_2_O_2_ resistance is positively correlated with thermal tolerance. While melanization is thought to be protective against reactive oxygen species [160], Jacobson et al. [70] observed no significant protection against H_2_O_2_ in *C. deneoformans* strains; thus observing a negative relationship between these two phenotypes is not unexpected. However, the inverse relationship between melanization and thermal tolerance is somewhat surprising, as prior studies have demonstrated a positive relationship between the production of melanin and the ability to grow at high temperatures [68].

A likely explanation for the mix of similar and dissimilar correlates with *RIC8* genotypes relative to earlier work, is divergence in cAMP-PKA signaling between *C. neoformans* and *C. deneoformans* strains. Hicks et al. and Hicks and Heitman [161, 162] showed that mutations of the PKA catalytic subunits, *PKA1* and *PKA2*, have distinctly different effects on melanization, capsule formation, and mating in *C. neoformans*, *C. deneoformans* and the more distantly related species *C. gattii*. Genetic variation that affects cAMP-PKA signaling in particular is an increasingly common theme in studies of fungal quantitative genetics and experimental evolution [163–166]. A recent comparative study of cAMP-PKA signaling in *S. cerevisiae* and related yeasts hypothesized that this pathway is likely to be a hotspot for functional variation and evolutionary adaptation [167]. We predict that segregating genetic variation in cAMP-PKA signaling may be particularly relevant for natural variation in virulence-related traits not only in *Cryptococcus* but in other pathogenic fungi as well.

### HOG pathway variants moderate resistance to the antifungal drug amphotericin B

Major QTL for amphotericin B sensitivity were found on chromosomes 2 and 12. We identified candidate QTG that underlie these loci – *SSK1* on chromosome 2 and *SSK2* on chromosome 12 both of which are components of the HOG signalling pathway. The HOG pathway is known to regulate the production of ergosterol, the target of amphotericin B [168].

At the chromosome 2 locus we discovered a single nucleotide insertion in the 431α parental strain background that leads to an early stop-gain in the gene *SSK1*, the response regulator of the HOG pathway. Given the location of the identified variant, it is likely that this results in a complete loss of function for the gene. *C. neoformans ssk1*Δ mutant strains exhibit increased sensitivity to amphotericin B compared to wild-type strains [148, 168], a pattern we observed in segregants with the *SSK1* ^431α^ allele. Interestingly, the *ssk1*Δ strains in the XL280**a** background we generated did not display an amphotericin B sensitive phenotype. We hypothesize that this was due to additional genetic variants in this background that also contribute to amphotericin B resistance.

In *C. neoformans*, but *not C. deneoformans* strains, Ssk1 has also been shown to govern capsule elaboration and melanization [169]. Consistent with this earlier study, allelic variation at *SSK1* is not associated with variation in either capsule or melanin phenotypes in our mapping population, suggesting that HOG signaling has diverged between *Cryptococcus* species.

The likely causal variants for amphotericin B sensitivity that mapped to chromosome 12 are alleles of the gene *SSK2*, the MAPKKK of the HOG pathway. The variant sites within *SSK2* observed here have been previously described by Bahn et al. [143]. In their study, Bahn et al. [143] showed that the allele present in the 431α parental strain (*SSK2* ^431α^) is associated with increased basal levels of Hog1 phosphorylation and in a follow-up study, Ko et al. [168] demonstrated that *C. deneoformans* strains with this allele are more resistant to higher concentrations of amphotericin B. Across *C. neoformans* and *C. deneoformans* strains, the amount of pre-phosphorylated Hog1 is known to vary [169], and Bahn et al. [143] hypothesized that this allowed some strains to rapidly respond to extracellular stresses. Interestingly in our study, *SSK2* ^431α^ was the only allele from the 431α parental strain associated with increased fitness.

### Epistatic interactions within and between signaling pathways

Ssk1 and Ssk2 are both members of the HOG pathway, and Ssk1 physically interacts with and regulates Ssk2’s kinase activity [24, 153, 170]. This naturally led us to explore genetic interactions between the allelic states we observed for *SSK1* and *SSK2*. Additionally, crosstalk between HOG and cAMP-PKA signaling pathways has been documented in both *S. cerevisiae* and *C. neoformans* [169, 171–174], thus motivating an exploration of genetic interactions between *RIC8* and HOG-pathway alleles.

We found that an epistatic interaction between *SSK1* and *SSK2* affected sensitivity to fludiox-onil, an agricultural antifungal drug. Fludioxonil’s mode of action is thought to be hyperactivation of the HOG pathway [175] and resistance to fludioxonil occurs primarily through HOG pathway loss-of-function mutations [145, 176]. Bahn et al. [143] showed that strains with the *SSK2* ^431α^ genotype were sensitive to fludioxonil, presumably due to hyperactive HOG signaling associated with pre-phosphorylated Hog1. Surprisingly, the parental strain 431α, which bears the predicted sensitive *SSK2* allele, was fludioxonil resistant. Analysis of offspring from our cross revealed that the unexpected resistance in 431α is mediated by allelic variation at *SSK1* – segregants with the sensitive *SSK2* ^431α^ allele as well as the loss-of-function *SSK1* ^431α^ allele exhibit fludioxonil resistance, while the pairing of the sensitive *SSK2* ^431α^ allele with the functional *SSK1*^XL280**a**^ allele results in sensitivity to fludioxonil. All other allelic combinations at these two loci are fludioxonil resistant. This thus represents a compelling example of within-pathway epistasis.

Our analysis of hydrogen peroxide resistance suggests complex cross-pathway epistasis between the HOG and cAMP-PKA signaling networks. We propose that the three-way epistatic interaction between *SSK1*, *SSK2*, and *RIC8* can be rationalized in terms of the relative balance between HOG signaling and cAMP-PKA signaling (Fig 10B). Segregants with the genotype *SSK2* ^XL280**a**^ *RIC8* ^431α^ exhibit the greatest average sensitivity to hydrogen peroxide stress. This genotypic combination is predicted to have weak or intermediate levels of HOG signaling but normal cAMP-PKA signaling. Conversely, segregants with the genotype *SSK2* ^431α^ *RIC8* ^XL280**a**^ are predicted to have intermediate or high HOG signaling activity (associated with pre-phosphorylation of Hog1), but weak cAMP-PKA signaling (due to Ric8 loss-of-function alleles), resulting in H_2_O_2_ resistance. The two genotype combinations are thus consistent with the observed marginal effects of *SSK2* and *RIC8*. The phenotypes of the other genotypic combinations of *SSK2* and *RIC8* require consideration of *SSK1* allelic state. For example, intermediate resistance phenotypes are associated with either intermediate HOG signaling (*SSK1* ^431α^ *SSK2* ^431α^) combined with normal cAMP-signaling (*RIC8* ^431α^) *or* weak HOG signaling (*SSK1* ^431α^ *SSK2* ^XL280**a**^) coupled with weak cAMP-signaling (*RIC8* ^XL280**a**^).

There is significant experimental evidence consistent with a model of opposing effects of HOG and cAMP-PKA signaling for a variety of stress responsive phenotypes in both *Cryptococcus* and other fungi. For example, Maeng et al. [177] report that *C. neoformans ras1*6. mutants show increased resistance to H_2_O_2_ stress but decreased resistance to diamide, while *hog1*6. mutants are H_2_O_2_ sensitive and diamide resistant. *C. neoformans hog1*6. mutants exhibit an increase in capsule size [169], another stress responsive phenotype, while cAMP related mutations such as *gpa1*6., *pka1*6., *cac1*6., and *ric8*6. have reduced or absent capsule [144, 178–180]. Gutin et al. [173] report evidence of crosstalk between cAMP-PKA and HOG signaling with respect to the general stress response in yeast, with cAMP-PKA activity associated with the repression of key stress responsive genes while HOG activity is associated with their activation.

### The importance of time for the study of microbial growth phenotypes

A critical feature of our study is the inclusion of temporal data and the application of function-valued analytical approaches for key growth traits. While there are both experimental [181] and statistical [134, 135] challenges associated with the collection of high-resolution time series growth phenotypes and functional valued QTL mapping, such data are much more information rich than conventional end-point growth or estimates of maximum growth rates. For example, several of the QTL associations we identified are temporally variable and would likely not have been detected without the inclusion of a time axis in our analyses.

The importance of temporal information for microbial growth phenotypes is not limited to QTL mapping, but is equally relevant to genetic approaches based on mutational analysis. For example, we found that when exposed to heat stress *C. neoformans ric8*Δ mutants exhibit slower maximum growth rates compared to an isogenic wild-type strain, but reach a higher final population density (Fig 7B). The determination of whether the mutant grows better or worse than wild-type thus depends critically on when during growth this question is posed and whether the investigator considers maximal growth rate or final population density to be a better reflection of thermal tolerance. Only by including the time axis and using a function valued approach can we appreciate the complexity of growth patterns inherent in this comparison.

While not explored in this report, we hypothesize that the relative timing of QTL or mutational associations with microbial growth phenotypes may reflect key transitions in physiological state within microbial populations. Temporal QTL/mutational information could thus be used to inform the construction of models relating the dynamical behavior of gene networks to distinct physiological states.

### Comparison to previous QTL studies in *Cryptococcus* and other fungi

There are commonalities between the results presented here and a previous QTL mapping study of *C. deneoformans*. For example, Lin et al. [111] examined variation in thermal tolerance and melanization within *C. deneoformans* progeny and identified a pleiotropic QTL on chromosome 7 that contributes towards both of these aforementioned traits. This QTL was narrowed down to allelic differences in *MAC1*, a copper homeostasis transcription factor. Similar to this past study, we also identified pleiotropic QTGs contributing to more than one virulence-related trait examined here (including thermal tolerance and melanization), namely the genes *RIC8* and *SSK2*. While our study did not share any of the previously implicated QTGs [79, 111], across several studies – not just those using QTL mapping strategies – observing pleiotropic effects of genes and pathways connected to virulence and virulence-associated traits seems to be a unifying phenomena [108, 182–187].

### The genetic complexity of virulence phenotypes

While the effects of each of the QTL we identified are relatively large, explaining on average 24% of the phenotypic variance in amphotericin B susceptibility at *SSK1* and *SSK2*, *∼*33% in thermal tolerance between *SSK2* and *RIC8*, 17% in capsule size, and 39% in melanization at *RIC8*, and 25% of variation in resistance to H_2_O_2_ between *SSK2* and *RIC8*, there are still large portions of unexplained phenotypic variation. Coupled with the observation of transgressive phenotypes in several of our experiments, these data may suggest the presence of many unidentified QTL and undiscovered epistatic interactions. Our analysis also focused primarily on the analysis of non-synonymous coding variants, though non-coding variation (e.g. [188, 189]) and synonymous variation (e.g. [190]) have both been shown to be important for the genetic architecture of complex traits in fungi and as well as other eukaryotes. Additional functional analyses, larger mapping populations and higher order models that test for multiple [191] and interacting loci [192] may help to detect these elusive QTL in future studies.

### Implications for the study of fungal virulence

Virulence is a complex outcome, an emergent property, that is determined by the combined effects of numerous morphological, physiological, metabolic, and molecular features of pathogens and their hosts [1]. The QTL and associated candidate genes and variants we have identified emphasize the genetic and functional complexity of virulence traits. For example, the *RIC8*^XL280**a**^ allele we identified is associated with decreased melanization but increased thermal tolerance, oxidative stress resistance, and capsule size. Despite low levels of melanin, and the loss of a key activator of the cAMP-PKA pathway, the strain XL280 is still virulent in an inhalation infection model of murine cryptococcosis [137, 193]. One must therefore exercise caution when trying to predict the likely effects of natural variation on virulence potential. This is likely to apply equally to engineered genetic manipulations or the effects of drugs that target particular pathways.

Our findings in the present study may also have implications for clinical treatment of cryptococcal disease. Outright resistance to antifungals such as amphotericin B is rare in *Cryptococcus* species [84, 194], but variance in the minimum inhibitory concentrations of antifungals, including amphotericin B, have been observed within species. Such variation could lead to recurring instances of disease within patients [49, 78, 195, 196]. A recent survey of antifungal susceptibility in clinical isolates of *C. neoformans* saw increases across a ten-year period in the minimum inhibitory concentration for both fluconazole and amphotericin B [85]. The QTN (at *SSK1* and *SSK2*) identified in the environmental strain 431α are implicated in sensitivity and increased resistance (respectively) to amphotericin B and provide concrete examples of the types of natural genetic variants that are present within *Cryptococcus* that might underlie differences in response to clinical treatment, depending on the particular lineage(s) that a patient is infected with. Similarly, in the case of infections by multiple *Cryptococcus* strains [48], naturally occurring alleles that decrease sensitivity to antifungals are likely targets for selection.

Current global trends point to both a warming planet and an increase rate of resistance to antifungal drugs [1, 197]. Studies like the one presented here, that focus on standing genetic variation within species, may help to predict the complex and evolving landscape of fungal virulence, providing insights into both lineages and genetic variants that are likely to be favored or disfavored as environments and clinical treatment change.

## Materials and methods

### Parental strains, laboratory crosses, and isolation of F_1_ progeny

As described in Sun et al. [92] the parental strains 431α, XL280αSS, and XL280**a** were used in α–α unisexual and **a**–α bisexual crosses (S1 Table). The parental strain 431α is a natural *C. deneoformans* isolate with the *MAT* α allele [92, 138]. The parental strain XL280αSS is an XL280 strain with an inserted *NAT* resistance marker in the *URA5* gene [141] and is congenic to the parental strain XL280**a** with the exceptions of the *URA5* gene, *NAT* resistance marker, the *MAT* locus, and a partial duplication of the left arm of chromosome 10 [92, 137, 141]. Due to the insertion of the *NAT* in the *URA5* gene of the XL280αSS strain, a wild type XL280α strain was used in phenotyping experiments. Because the strains XL280α and XL280**a** are congenic with the exception of the *MAT* locus, throughout the manuscript only the XL280**a** strain is referred to when referencing the XL280 background.

As described in [92], both **a**–α bisexual (XL280**a** *×* 431α) and α–α unisexual (XL280αSS *×* 431α) matings were carried out and progeny were isolated, yielding 261 and 156 progeny respectively. Parental strains and segregants were maintained in 35% glycerol frozen stocks at −80*^◦^*C and subcultured from freezer stock to YPD media for experimentation. Between segregants derived unisexually verses bisexual, no significant effect was observed in any of the phenotypes examined here – except for fludioxonil sensitivity (see results) – and their phenotypic values were pooled into a single mapping population for use in QTL mapping.

### Sequencing, aligning, and variant calling

In total, 127 segregants, which included 63 from the α–α unisexual, 61 from the **a**–α bisexual crosses, and the 3 parental strains, XL280**a**, XL280αSS, and 431α, were sequenced as previously described [92, 141]. Raw reads were aligned to an XL280α *C. deneoformans* reference genome [137] using BWA (v0.7.12-r1039, [198]). Variant calling was carried out using SAMtools (v0.1.19-96b5f2294a, [199]) and FreeBayes (v1.2.0, [200]) resulting in 449,197 bi-allelic single nucleotide polymorphisms (SNPs) and 1,500 bi-allelic insertions and deletions (INDELs) between the parental strains, segregating within the F_1_ segregants.

### Segregant filtering and marker creation

Each of the 127 segregants were filtered to remove those which exhibited aneuploidy, clonality, or lack of recombination as described in Roth et al. [141]. After applying these filtering criteria, 104 segregants – composed of 55 progeny from α–α unisexual crosses, 46 progeny from **a**–α bisexual crosses, and the three progenitor strains – were retained for further analysis.

The 449,197 bi-allelic variants sites were filtered on call rate, read depth, allelic read depth ratio, minor allele frequency, and quality scores. Across the 104 segregants, SNP and INDEL sites were required to have 100% call rate, greater than 10*×* coverage in read depth, an allelic read depth ratio of 80% (for example, if a SNP site has 10 reads mapping over it, 8 of the 10 reads must support the existence of the SNP), a minor allele frequency of 20%, and a log_10_ quality score, normalized by read depth, of greater than or equal to 0.75. A maximum log_10_ read depth of 4.1 was set to filter out SNPs in regions with repetitive elements. Finally, bi-allelic SNP and INDEL sites within 5 kb of centromeres and of the ends of the chromosomes were removed as these regions are difficult to sequence, resulting in 92,103 genetic variants. These 92,103 genetic variants were then grouped into haploblocks (“haplotype blocks”) based on variants in perfect linkage, in order to reduce the number of markers for analysis in genotype-phenotype association tests [201]. This was done for each chromosome such that every haploblock had at least one segregant with a genotype change between contiguous haploblocks, resulting in 3,108 sites across the segregants. The average size of haploblocks was 5.4 kb with a minimum size of 4.4 kb and maximum size of 6.3 kb.

### Quantitative growth assays

Quantitative growth assays were measured using absorbance microplate readers (Tecan Sunrise). Initially, segregants were arrayed in U-bottom 96 well plates containing 100 *µ*l of liquid YPD, incubated for two days at 30*^◦^*C, and after the addition of glycerol, preserved as frozen stocks at −80*^◦^*C. Plates were stored as frozen stocks and used to start each assay. After two days of growth on YPD solid agar plates, segregants were pinned into 150 *µ*l liquid YPD and grown on a plate shaker for two days at 30*^◦^*C. Subsequently for each segregant, 1 *×* 10^5^ cells were transferred into 150 *µ*l BD Difco yeast nitrogen base (YNB) buffered to pH 7.0 with 0.165 M MOPS (morpholine-propanesulfonic acid) buffer (Sigma-Aldrich). For drug treatments, amphotericin B (Sigma-Aldrich) was added to YNB from a stock solution of 100 *µ*g/ml for final drug concentrations of 0.075, 0.125, and 0.175 *µ*g/ml. Cells were grown in microplate readers for three days at either 30*^◦^*, 37*^◦^*, or 39*^◦^*C with no drug and assayed for all possible combinations of temperature by amphotericin B concentration except for 39*^◦^*C and 0.175 *µ*g/ml because no strains grew at this combination of temperature and drug stress. Assays were replicated four times. To monitor growth, optical density measurements were made at a wavelength of 595 nm (OD_595_) every 15 minutes. To prevent fogging in the machines, plate lids were pre-treated with a solution of 0.05% Triton X-100 in 20% EtOH.

### Plate based assays

For plate based assays using solid agar media, segregants were pinned from liquid YPD media to the appropriate, freshly prepared assay media. To minimize edge effects, no segregants were arrayed in the outer rows and columns of the plates. Instead, a control strain was grown in these positions, either H99α or JEC21α.

To assess melanin production, segregants were grown on chemically defined minimal medium containing L-DOPA (7.6 mM L-asparagine monohydrate, 5.6 mM glucose, 10 mM MgSO4, 0.5 mM 3,4-dihydroxy-L-phenylalanine, 0.3 mM thiamine–HCl, and 20 nM biotin) and incubated at 30*^◦^*C for three days in the dark. After three days plates were scanned on an Epson Expression 10000 XL Flatbed Scanner in reflective mode (scanned from below) at 300 dpi. The grayscale intensity of each colony, as measured using ImageJ, was used as a proxy for melanization, and the mean across three replicates of these values was utilized in statistical association tests. The average Spearman rank correlation coefficient between replicates values was approximately 0.98.

To assay for hydrogen peroxide (H_2_O_2_) sensitivity, segregants were pinned onto YPD plates supplemented with 5mM H_2_O_2_ and incubated at 30*^◦^*C. After five days, plates were scanned in the same manner as above. Across four replicates, colony growth was scored manually by two individuals on a scale from 0 to 5. The median score per segregant was used for data analysis. Between replicates, the average Spearman rank correlation coefficient was 0.86.

To examine sensitivity to fludioxonil, segregants were similarly pinned onto YPD plates supplemented with 100 *µ*g/ml fludioxonil and incubated at 30*^◦^*C for five days. Scanned images of colonies were scored by two observers on a binary scale (growth or no growth) and this value was used in analysis. This assay was replicated in an additional, larger set of segregants generated using the same parental backgrounds (see below).

To assess the response to osmotic stress in this mapping population, colonies were pinned to YP medium containing 1.0 M NaCl, incubated for three days, and then scanned in transmissive mode. The average grayscale density across three replicates of each colony was used as the growth phenotype. Across replicates the average Spearman rank coefficient was 0.80.

### Capsule induction and imaging

Overnight cell cultures were resuspended in 9.0 ml CO_2_-independent medium (Gibco) and incubated for three days at 37*^◦^*C with shaking at 150 rpm. After incubation, cells were washed and then stained with India ink [202]. Each strain was imaged at least three times on an EVOS M5000 Cell Imaging System using a 40X objective. Using ImageJ, total area of the capsule plus cell and cell body only were measured and used to quantify capsule size for approximately 30 cells per segregant. The mean of these values was used in QTL mapping.

### Growth curve base-lining and parameter estimation

For each growth experiment, a blank optical density was calculated from the average optical density of wells containing no cells, and this value was subtracted from each well on a per plate basis. The first two time points, representing the first fifteen minutes of data collection, were dropped from analysis.The next five time points were used to baseline the data by calculating the average optical density of these points (the first 1.5 hours) and subtracting this from the remaining sampled time points. These five baseline points were then set to zero. After blank correction and baselining, individual growth curves were filtered using a median filter [203] with a moving, symmetric window of 25 time points, padding the beginning and end of the time courses with zeros or the final OD, respectively. After base-lining and median filtering, we estimated the area under the growth curve at each time point *n* as 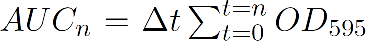 with Δ*t* = .25 hours (or 15 minutes). For each segregant, the median of the *AUC_n_* across replicates with respect to time was used in QTL mapping across the experimental conditions. One replicate at the condition of 30*^◦^*C and 0 *µ*g/ml of amphotericin B was dropped from analysis for 60 of the segregants due to poor initial growth seen in pre-culture plate. At 30*^◦^*C, in the absence of drug, the average Spearman rank correlation coefficient between replicates was 0.82.

### QTL mapping

For each plate based assay and the 3,108 haploblock test sites, a marker regression frame work was used to associate genotypes to phenotypes. The genotype of each haploblock was coded as zero if inherited from the XL280**a** (or XL280αSS) parental strain, and one if from the parental 431α strain. The model used in statistical association tests can be summarized as *y* = *µ* + *β***I***_c_* + *E* where *E* is the error term, *µ* is the average phenotype (i.e. mean intensity, growth score, capsule or cell size), **I***_c_* is an indicator variable for genotype, and *β* is the coefficient depicting the effect of having the genotype of XL280**a** (or XL280αSS) or genotype of 431α at the given haploblock. An estimation of this effect, *β*, is given by 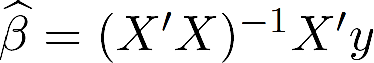 [204]. The *−log*_10_(p-value) for each unique haploblock, for a given experiment and phenotype was used as the measure of association. The 95% confidence intervals for melanization, capsule size, and thermal tolerance were calculated as described in Visscher et al. [205], sampling a 1,000 times with replacement and taking the mean location per maximum haploblock.

For experiments that generated colony growth curves, a function-valued, marker-regression approach was employed to quantify the relationship between genotype and growth phenotypes for each variable haploblock across the 72-hour time course. For these experiments, the area under the growth curve (*AUC*) was calculated at 15-minute intervals and used as the growth phenotype. As described above, the usual marker-regression model is *y* = *µ* + *β***I***_c_* + *E*, with *y* = *AUC*. Across time, the *AUC_t_* can be calculated for a given time point within the 72-hour time course and treated as separate phenotypes (Fig 4). This marker-regression model may then be extended for the functional phenotype dependent on time, *y*(*t*), where, *y*(*t*) = *µ*(*t*) + *β*(*t*)**I***_c_* + *E*(*t*). An estimate of the QTL effect across time is then given by 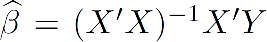 where *Y* is a matrix of segregant phenotypes with columns that represent the multiple time points [134, 206].

### Permutation tests

Permutation tests, as described in Churchill and Doerge [142], were conducted to establish significance thresholds for the *−log*_10_(p-value) from QTL mapping. The number of permutations for all phenotypes analyzed here was 10,000. The same set of random reassignments of genotype to phenotype were used for the eleven temperature by amphotericin B experimental conditions to conserve correlation structure between the experimental conditions. Each growth curve was treated as a single phenotypic measure during permutations to preserve autocorrelation across time points, and the significance thresholds for the maximum and mean associations between genotype and phenotype (with respect to time) were calculated [134]. The 95^th^ and 99^th^ percentile of the permuted distribution of genotype-phenotype associations was used to estimate thresholds for significance.

### Analyzing three-way epistasis

We used an information-theoretic approach proposed by Hu et al. [146] to evaluate models of three-way genetic epistasis for the association of hydrogen peroxide resistance with genotypic variation at *SSK1*, *SSK2*, and *RIC8*. Hu et al.’s method is based on a measure of the relative amount of “information gain” (IG) attributable to synergistic epistatic interactions, quantified in terms of standard information theoretic measures of mutual information. To apply this approach we discretized each segregant’s H_2_O_2_ response as sensitive, intermediate, or resistant based on the observed growth scores, and estimated Hu et al.’s IG statistic using mutual information functions implemented in the Python package scikit-learn [207]. We used permutation tests to simulate null distributions and estimate *p*-values for the IG statistic. For single locus effects, we used permutations which randomized the relationship between phenotype and genotype at each of the three loci, while maintaining the genotypic covariance between loci. For second- and third-order effects we permuted genotypes of samples within each phenotypic class, preserving the independent main effects while randomizing any non-linear interactions, as recommended in Hu et al. [146]. A thousand permutations were used to simulate the distributions of both main and higher order effects. We report normalized information gain, expressed as a percentage of the Shannon entropy of the phenotypic distribution.

### Annotation realignment and genetic variant effect prediction

To predict the effects of genetic variants identified between the XL280**a** (or XL280αSS) and 431α parental strains, annotated gene features were derived from the *C. deneoformans* reference strain, JEC21α [208]. These sequences were then aligned via the blast-like alignment tool (BLAT, [209]) to the XL280α reference genome [137]. Per gene, alignments were filtered for sequence identity of 95% and at most two mismatches between the JEC21α and XL280α genomes. There are a total of 5,210 annotated features in the JEC21α genome annotation, of which 4,800 mapped perfectly and uniquely to the XL280α genome. After mapping orthologous genes, the effects of genetic differences between the XL280**a** (or XL280α) and the 431α backgrounds were imputed with respect to the predicted exonic and intronic regions.

### Gene disruption

TRACE (Transient CRISPR-Cas9 Coupled with Electroporation [56]) was used to genetically disrupt *SSK1* in the parental strains. Deletion constructs were assembled with two-step PCR using homologous arms one kb in length and a *NAT* marker, a dominant drug-resistance marker conferring resistance to nourseothricin. Single-guide RNAs were designed with Eukaryotic Pathogen CRISPR guide RNA/DNA Design Tool (http://grna.ctegd.uga.edu/) using default parameters, and the gRNA scaffold was amplified from pDD162. To generate complete gRNAs, a one-step overhang PCR was used to amplify the construct from sgRNA, the scaffold, and the U6 promoter (JEC21α), following TRACE protocols. Cas9 was amplified from pXL1-Cas9-HygB.

Parental strains were transformed with the amplified constructs following the protocol for electroporation in Fan and Lin [56], except competent yeast cells were washed and resuspended in 1M sorbitol before transformation. Electroporated cells recovered for two hours in YPD before being plated onto YPD supplemented with nourseothricin (YPD+*NAT*) selective media. After restreaking, transformants were screened with internal *SSK1* primers. Colonies that were capable of growing on YPD+*NAT* selective media were subsequently screened with external primers for product size, primers that spanned across the gene boundaries, and primers for detecting the presence of Cas9. One *ssk1* transformant for 431α and three *ssk1* transformants for XL280**a** were identified. The inserted deletion construct for each of these transformants was sequenced in full for confirmation. Primers are listed in S2 Table. To assess gene disruption effects, growth curves for the *ssk1* strains were measured at 30*^◦^*C in 0.125 *µ*g/ml amphotericin B, the condition with the largest strength in association at the chromosome 2 QTL which contains the gene *SSK1*.

### Additional fine-mapped crosses, segregant isolation, and sequencing

To further investigate the identified QTL on chromosome 2 (approximately 150-kb wide), fine-mapping techniques were applied to generate additional progeny. Parental strains were transformed as described previously [56] using the *NAT* and *NEO* selectable markers inserted at intergenic regions flanking the QTL. The chosen intergenic regions were between genes CNB02680 and CNB02690 at approximately 797,055 – 797,281 kb on the left and between genes CNB03490 and CNB03500 at approximately 1,047,138 – 1,047,346 kb on the right of the chromosome 2 QTL for the *NAT* and *NEO* markers respectively. Transformants were screened as described previously and one strain transformed from the XL280 strain background, with *NAT* cassette and three strains transformed from the 431α strain background, each with the *NEO* cassette, were identified (S1 Table). Southern blot probing for the selectable markers was used to determine that only one copy was inserted in the genome for each transformant.

To generate recombinant progeny from the transformed parental strains, spores from mass matings were purified through Percoll gradient centrifugation [22, 210]. Purified, recombinant spores were selected for by growing progeny on YPD+*NAT* +*NEO*, and *NAT^R^ NEO^R^* segregants were verified as recombinant by colony PCR. In total, 192 progeny were sequenced on a NovaSeq 6000 (SP) flow cell (150 bp PE). Sequencing data were analyzed in the same manner as described previously. Segregants were filtered to remove clones and progeny with diploid or aneuploid genomes and 173 segregants were retained for analysis.

### Data availability and software

Raw sequence reads generated from samples utilized in this study are available on NCBI’s sequence read archive under BioProject identification number PRJNA420966, with individual accession numbers SRR6352893 – SRR6352999, SRR10810110 – SRR10810130, and SRR10861770 - SRR10861961. The generated variant call file from the aligned sequenced reads along with the software developed for both analysis and figure generation are publicly available on GitHub: https://github.com/magwenelab/crypto-QTL-paper.

## Acknowledgements

This work was supported by NIH grant RO1 AI133654 and the GCB Summer Scholars program in Genome Sciences and Medicine (NIH R25) awarded to AS. We would also like to thank our collaborators for their comments on the manuscript: Magwene lab members Thomas Sauters and Lydia Hendrick, members of the Heitman lab, Shelby Priest and Dr. Marcia David Palma (Ph.D) along with Dr. David Tobin (Ph.D) and Tobin lab member, Jared Brewer.

## Author contributions

PM, JH, SS, CF, DM, and CR designed experiments. SS, AA, and CF provided strains and materials. CF, AS, and DM conducted experiments and generated data. PM and CR analyzed the data and CR created figures. PM, DM, and CR wrote the manuscript. JH, SS, AA, and CF provided edits to the manuscript.

## Supplemental information

**S1 Table.** Genotypes of parental and transformant strains. QTL-L and QTL-R refer to intergenic regions on chromosome 2 at 797,055 – 797,281 bp, between genes *CNB02680* and *CNB02690* and at 1,047,138 – 1,047,346 bp, between genes *CNB03490* and *CNB03500*, respectively.

| Strain | Name | Genotype | Source |
| --- | --- | --- | --- |
| SSA837 | 431α | wild type | [92] |
| SSB830 | XL280a | wild type | [211] |
| SSA853 | XL280αSS | *MAT*α, *ura5::NAT* | [92, 141] |
| PMY2408 | CF1705 | *MAT*α, QTL-R::*NEO* (431α) | this study |
| PMY2420 | CF1706 | *MAT*α, QTL-R::*NEO* (431α) | this study |
| PMY2432 | CF1707 | *MAT*α, QTL-R::*NEO* (431α) | this study |
| PMY2444 | CF1730 | *MATa*, QTL-L::*NAT* (XL280a) | this study |
| PMY2552 | 1a-12 | *MAT*α, *SSK1::NAT* (431α) | this study |
| PMY2553 | 4b-2 | *MATa*, *SSK1::NAT* (XL280a) | this study |
| PMY2554 | 5a-8 | *MATa*, *SSK1::NAT* (XL280a) | this study |
| PMY2555 | 5b-1 | *MATa*, *SSK1::NAT* (XL280a) | this study |

**S2 Table. Primer sequences used in this study.**

**S3 Table. Predicted ORF and summaries of genetic variants within QTL regions.** For each gene, the number of genetic variants within and upstream of a predicted gene is provided. For each gene the upstream distance from the 5’ UTR was taken as the intergenic distance (maximum 500 bp) between flanking genes on the same strand. For those genes containing genetic variants the predicted protein length of the reference strain JEC21α, parental strain XL280a, and the parental strain 431α is listed. The number of predicted stop-codons in the parental strains, non-synonymous changes, and variants within UTRs, exons and introns are also listed. Gene names are given in the *c. deneoformans* reference strain JEC21α background [208]. The position and strand are relative to the XL280α strain [137].

---

**S1 Fig.**
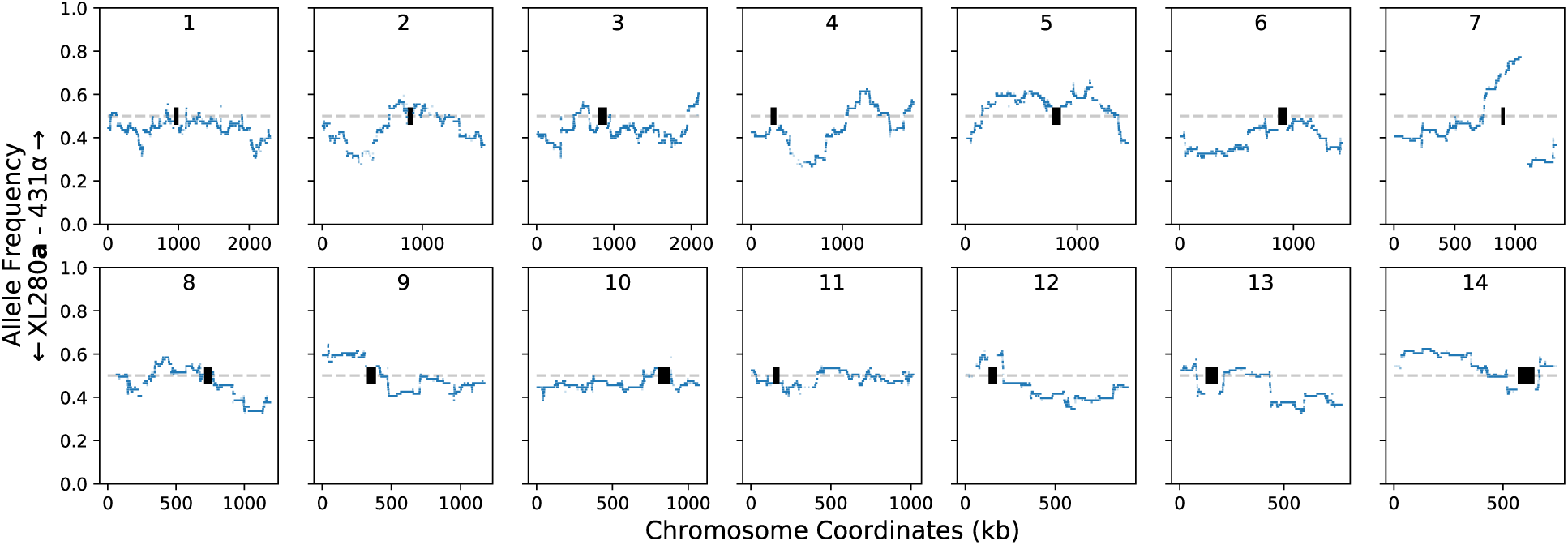
Genome-wide haplotype allele frequencies. *C. deneoformans* strains XL280a and XL280αSS were crossed with 431α in a–α bisexual and α–α unisexual matings, generating 101 segregants. Between the parental strains there are 92,103 bi-allelic genetic variants (see methods) and these genetic variants are collapsed across the segregants, based on genetic exchange events, generating 3,108 unique haplotypes across the genome. The allele frequencies of these haplotypes (blue dots) per chromosome are shown for each of the 14 chromosomes (numbers denote chromosome). A horizontal, grey dashed line marks an allele frequency of 0.5. Centromere locations are marked by black rectangles. The bias present on the right of chromosome 7 is due to selectable genetic markers used to generate progeny from the α–α unisexual cross [141].

**S2 Fig.**
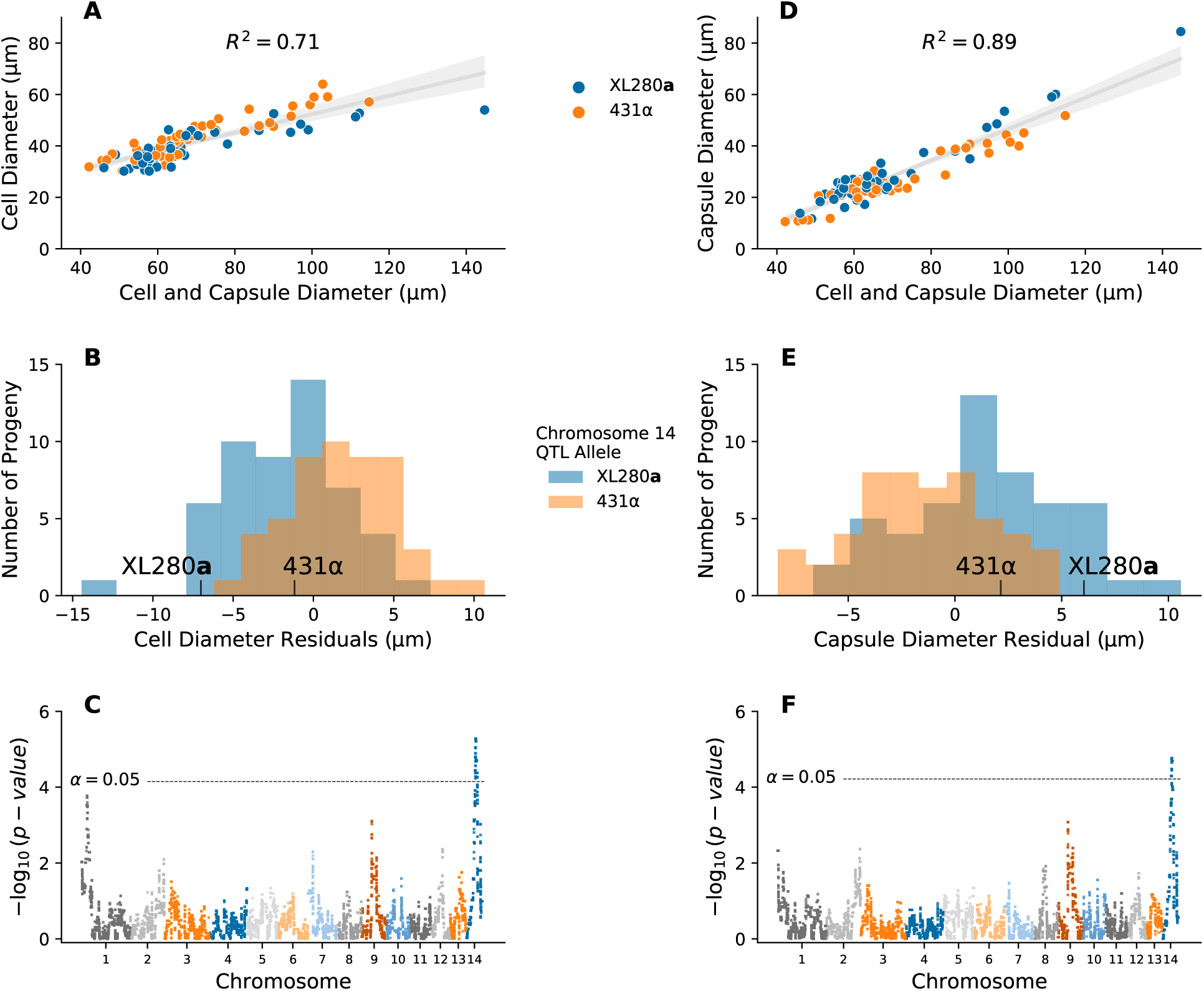
QTL analysis of variation in cell and capsule size. **A** – **C**) Analysis of variation in cell diameter (y-axis) as a function of cell and capsule diameter (x-axis, **A**), a histogram of the cell diameter residuals used in QTL mapping (**B**), and associated Manhattan plot (**C**). **D** – **F**) Analysis of variation in capsule diameter (y-axis) as a function of cell and capsule diameter (x-axis, **D**), a histogram of the capsule diameter residuals used in QTL mapping (**E**), and temporaassociated Manhattan plot (**F**). Grey lines and shaded regions in A and D represent regression models and associated 95% confidence intervals. The variation explained by these models is annotated within each plot. For both cell and capsule diameter residuals a QTL is detected on chromosome 14. Dotted horizontal lines represent significance thresholds from permutation tests. Progeny cell and capsule diameter and cell and capsule diameter residual values are colored by the chromosome 14 QTL allele.

**S3 Fig.**
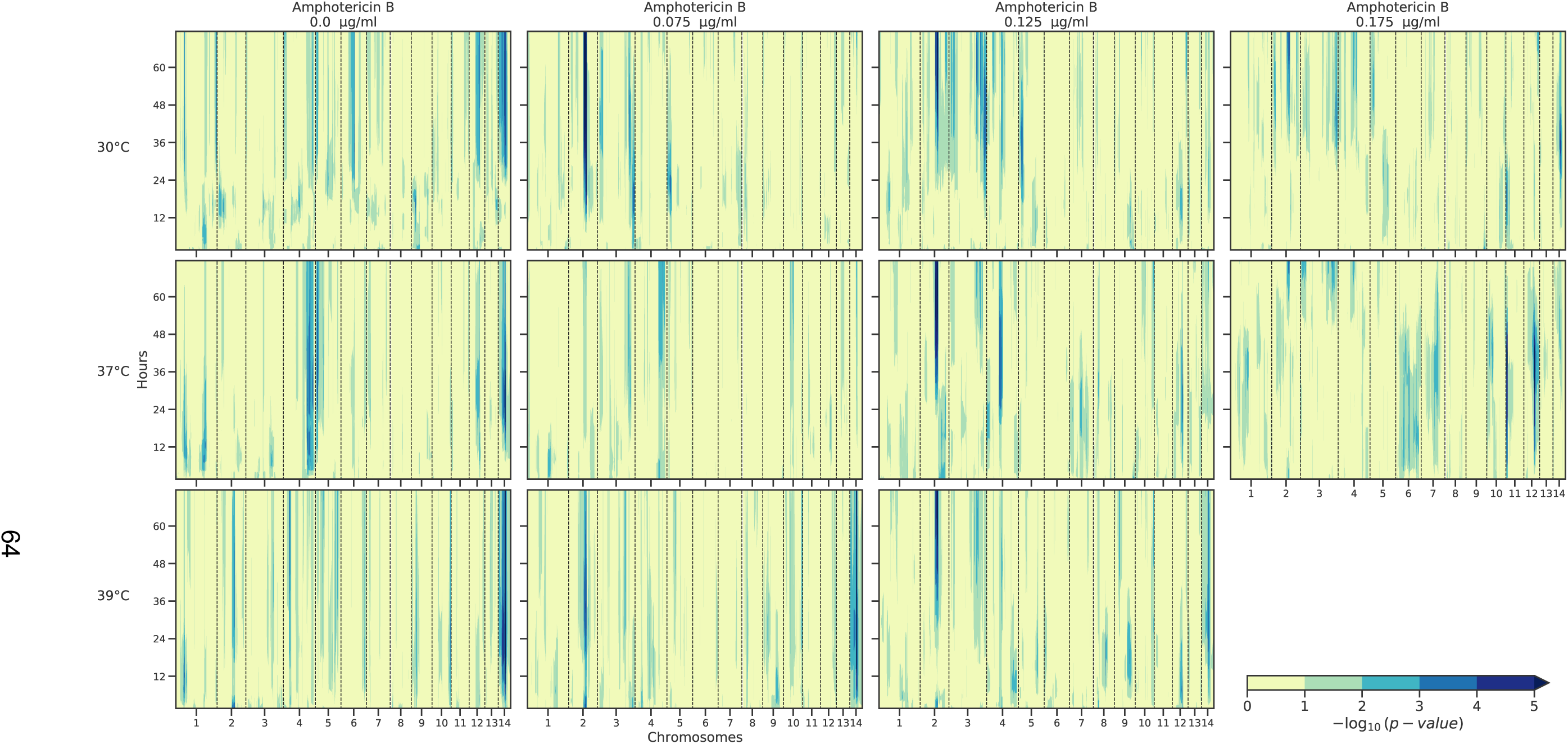
Genome-wide temporal Manhattan plot. Genome-wide Manhattan heat maps of association between genotype and phenotype across 72 hour for combinations of temperature (rows) and amphotericin B (columns) concentrations in Fig 3. Across combinations of temperature and amphotericin B stress, the median growth AUC of segregants, calculated every 15 minutes for each 72-hour time course, was regressed onto the parental genotypes of XL280**a** and 431α. The yellow to blue colors depict the strength in association (as measured by the *log*_10_(*p value*) from the linear regression) between the growth AUC values and 3,108 bi-allelic haploblocks across segregants (x-axis) along the 72-hour time course (y-axis).

**S4 Fig.**
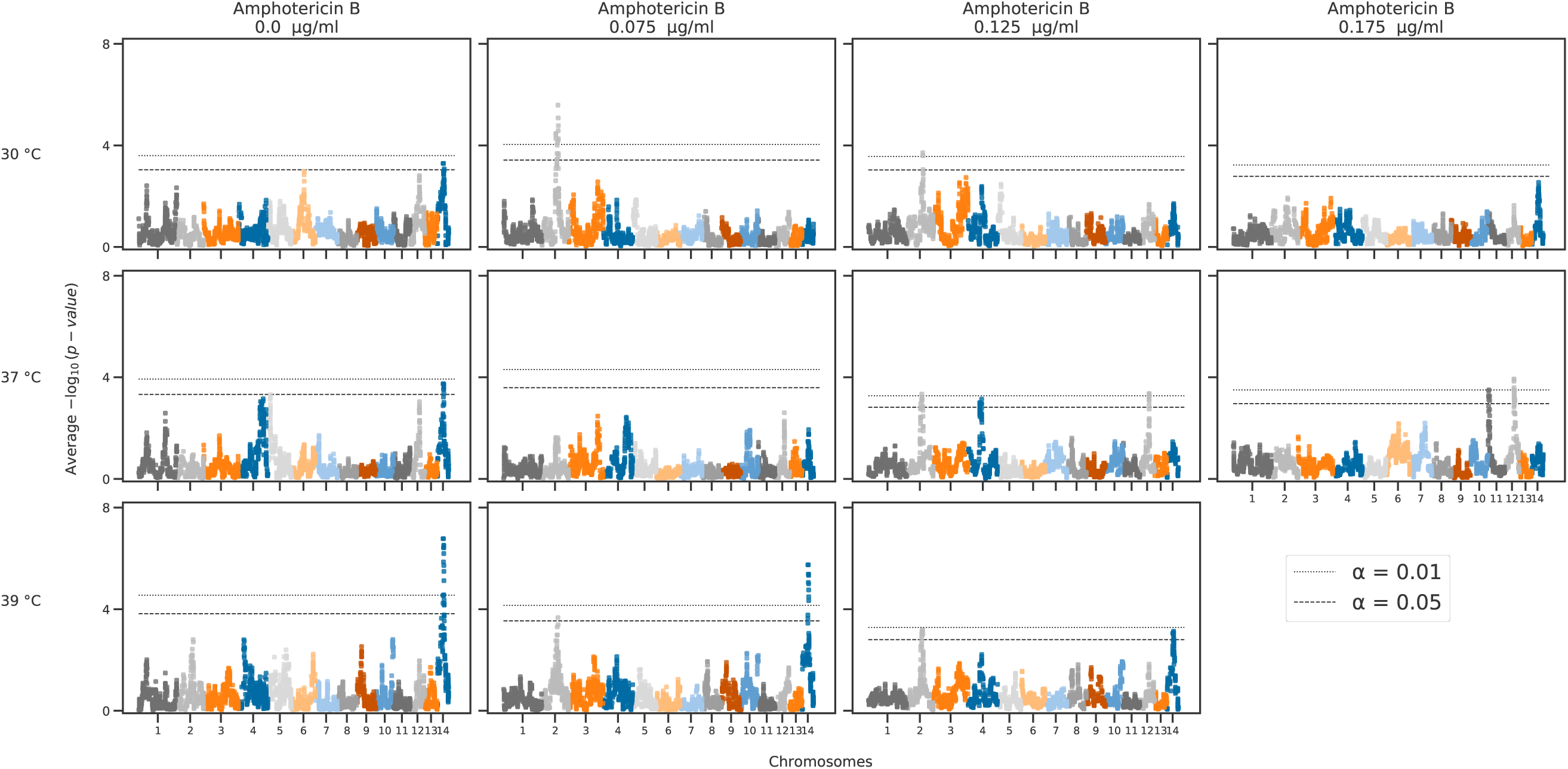
Genome-wide temporal average Manhattan plot. Genome-wide Manhattan plots of average association across time between genotype and phenotype for combinations of temperature (rows) and amphotericin B (columns) stress. For each experimental condition in Fig 3, the median growth AUC of segregants across the 72-hour time course was regressed onto the parental genotypes of XL280a and 431α. The x-axis represents positions along chromosomes (separated by colors) of 3,108 bi-allelic genetic variant sites, collapsed into haploblocks across segregants, and the y-axis is the average association across the 72-hour time course between genotype and the growth AUC values. Significance thresholds (horizontal dashed and dotted lines) were determined via permutation.

**S5 Fig.**
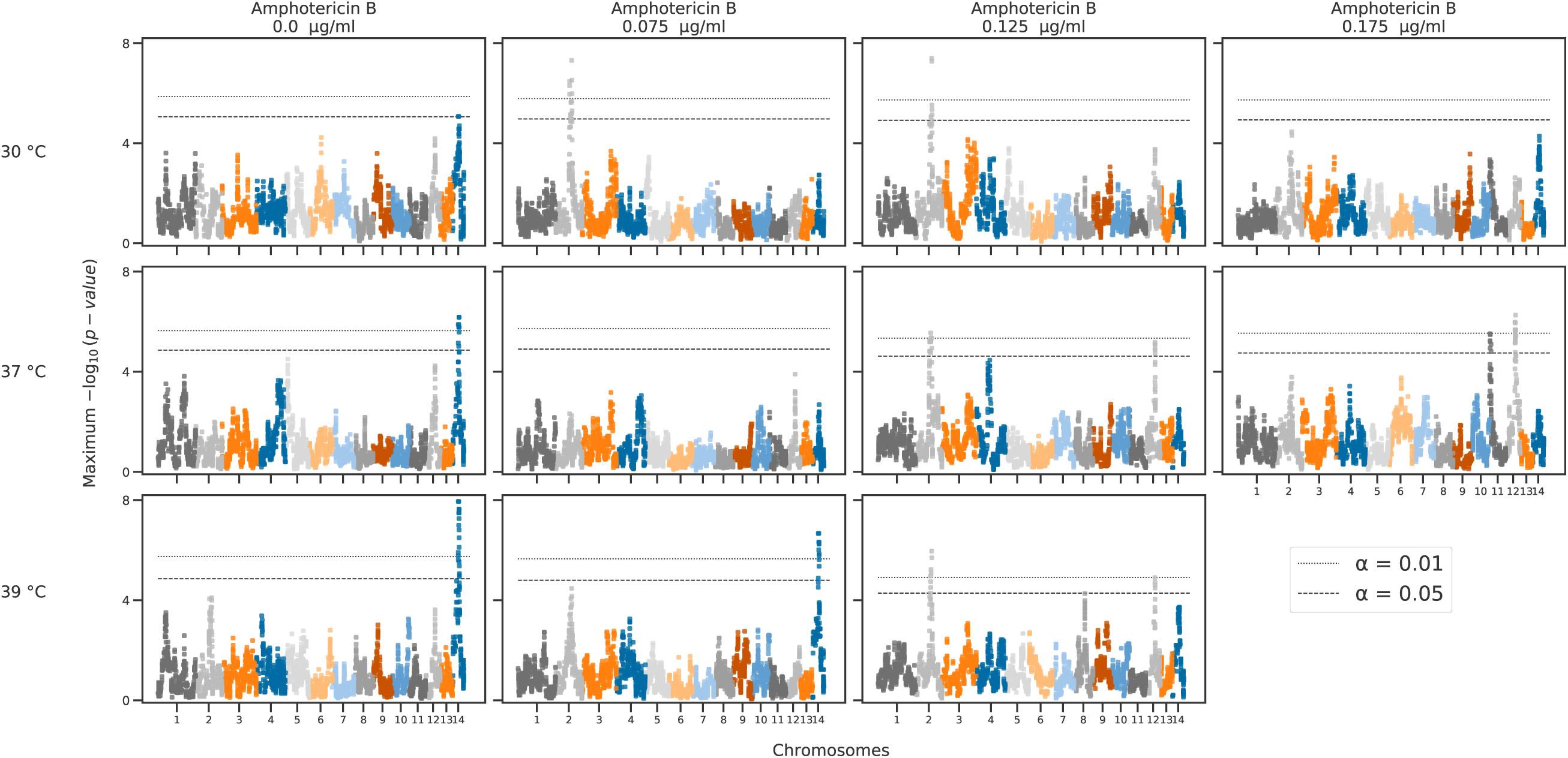
Genome-wide temporal maximum Manhattan plot. Genome-wide Manhattan plots of maximum association across time between genotype and phenotype for combinations of temperature (rows) and amphotericin B (columns) stress. For each experimental condition in Fig 3, the median growth AUC of segregants across the 72-hour time course was regressed onto the parental genotypes of XL280a and 431α. The x-axis represents positions along chromosomes (separated by colors) of 3,108, bi-allelic genetic variant sites, collapsed into haploblocks across segregants, and the y-axis is the maximum association across the 72-hour time course between genotype and the growth AUC values. Significance thresholds (horizontal dashed and dotted lines) were determined via permutation.

**S6 Fig.**
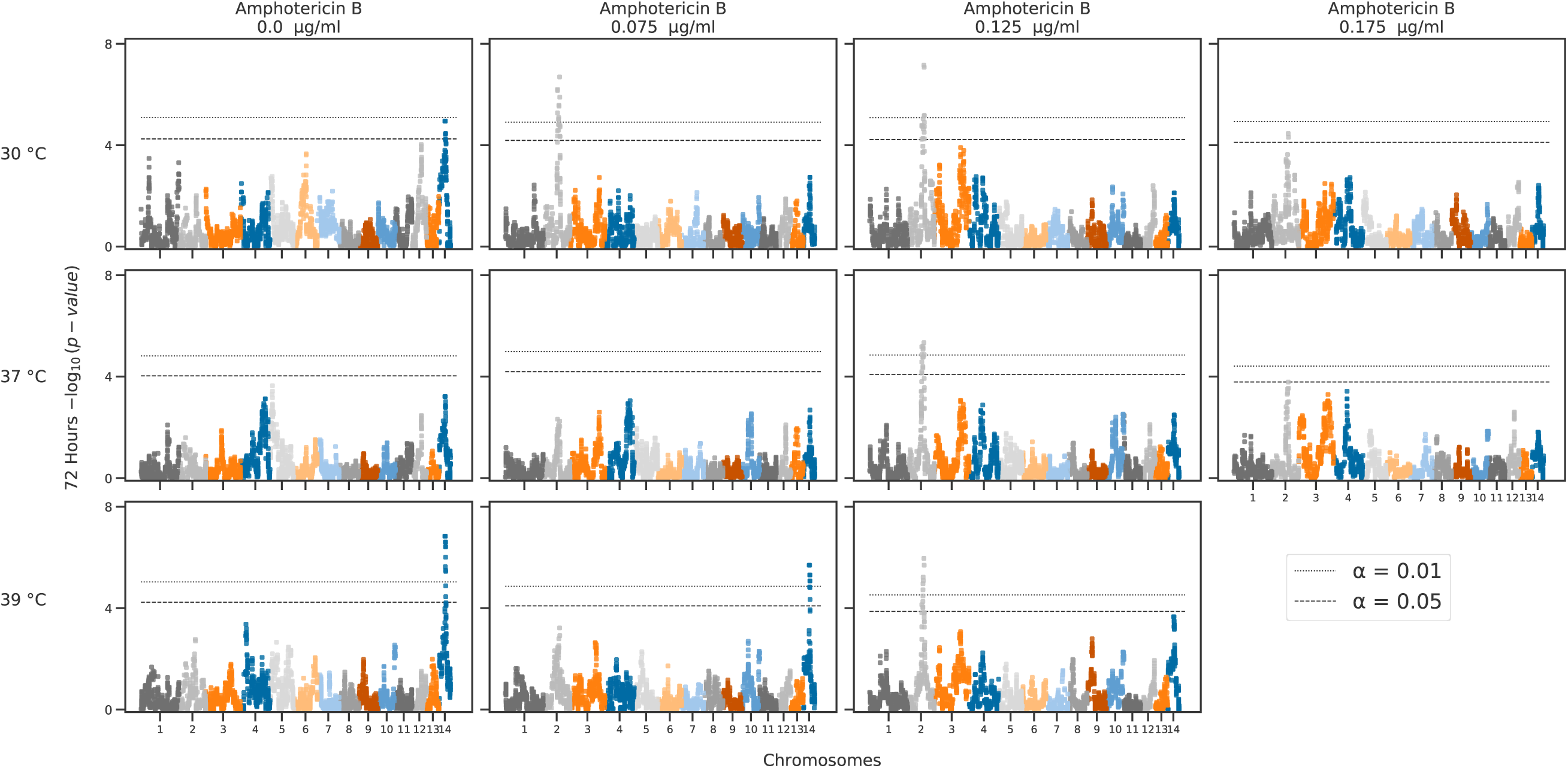
Genome-wide 72-hour Manhattan plot. Genome-wide Manhattan plots of association between genotype and phenotype for combinations of temperature (rows) and amphotericin B (columns) stress. For each experimental condition in Fig 3, the median growth AUC of segregants at 72 hours of segregants was regressed onto the parental genotypes of XL280a and 431α. The x-axis represents positions along chromosomes (separated by colors) of 3,108 bi-allelic genetic variant sites, collapsed into haploblocks across segregants and the y-axis is the association between genotype and the growth AUC values at 72 hour. Significance thresholds (horizontal dashed and dotted lines) were determined via permutation.

**S7 Fig.**
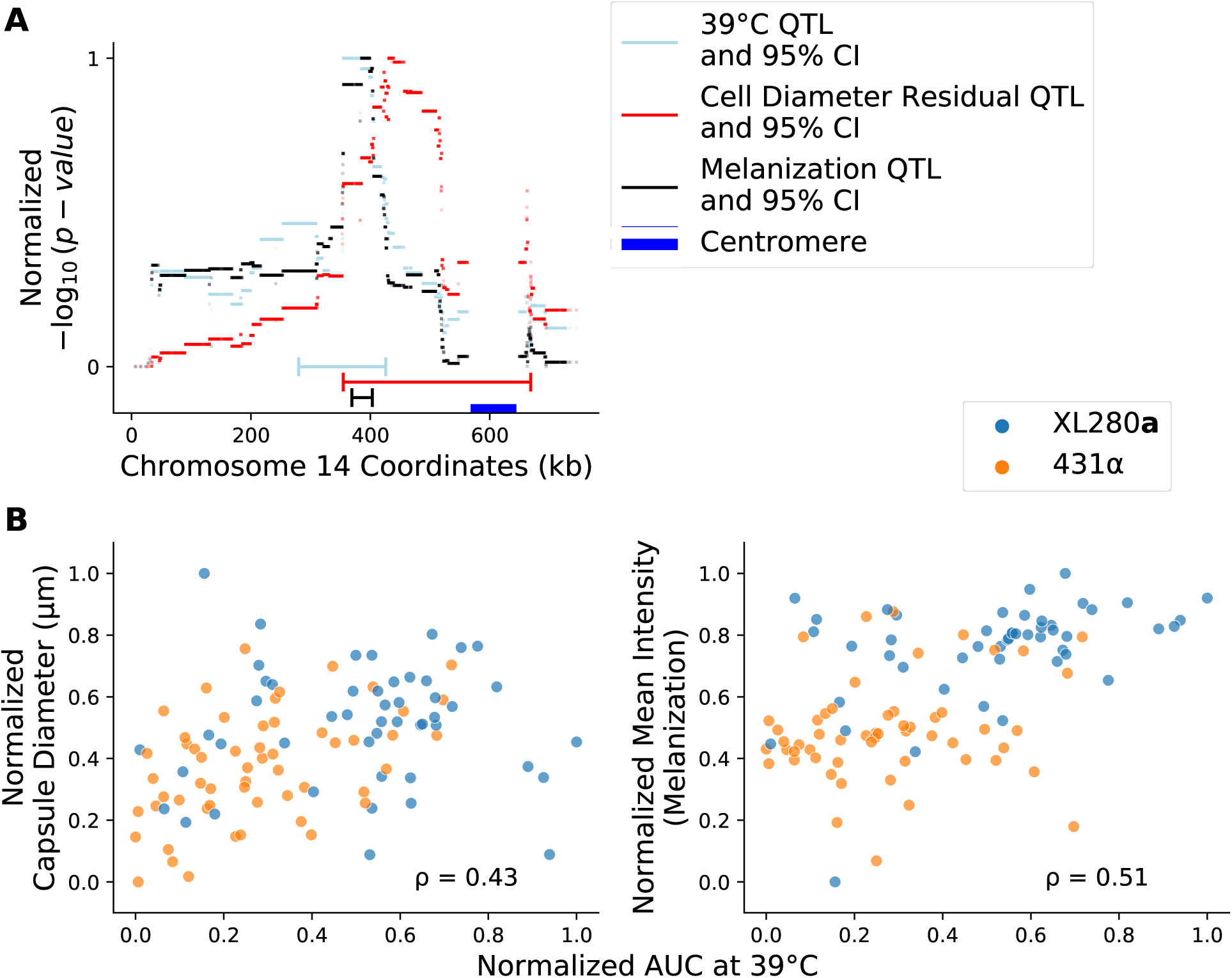
Pleiotropic QTL along chromosome 14. **A**) The three QTL and associated confidence intervals (CI) for area under the curve at 39*^◦^*C (light blue), cell diameter residuals (red), and melanization (black). Horizontal bar bells represent 95% confidence intervals. The location of the centromere on chromosome 14 is marked by a horizontal blue bar. **B**) Phenotypic relationships between capsule diameter and melanization (y-axis of left and right panels, respectively) as a function of growth at 39*^◦^*C (x-axis). Progeny values are colored by the allele at peak of the melanization QTL in **A**; blue for XL280a and orange for 431α. The Spearman rank correlation (*ρ*) between each pair of phenotypes is annotated within each plot. All QTL and phenotypic values in **A** and **B** (respectively) are re-scaled – in order to share the same scale – using max-min normalization.

**S8 Fig.**
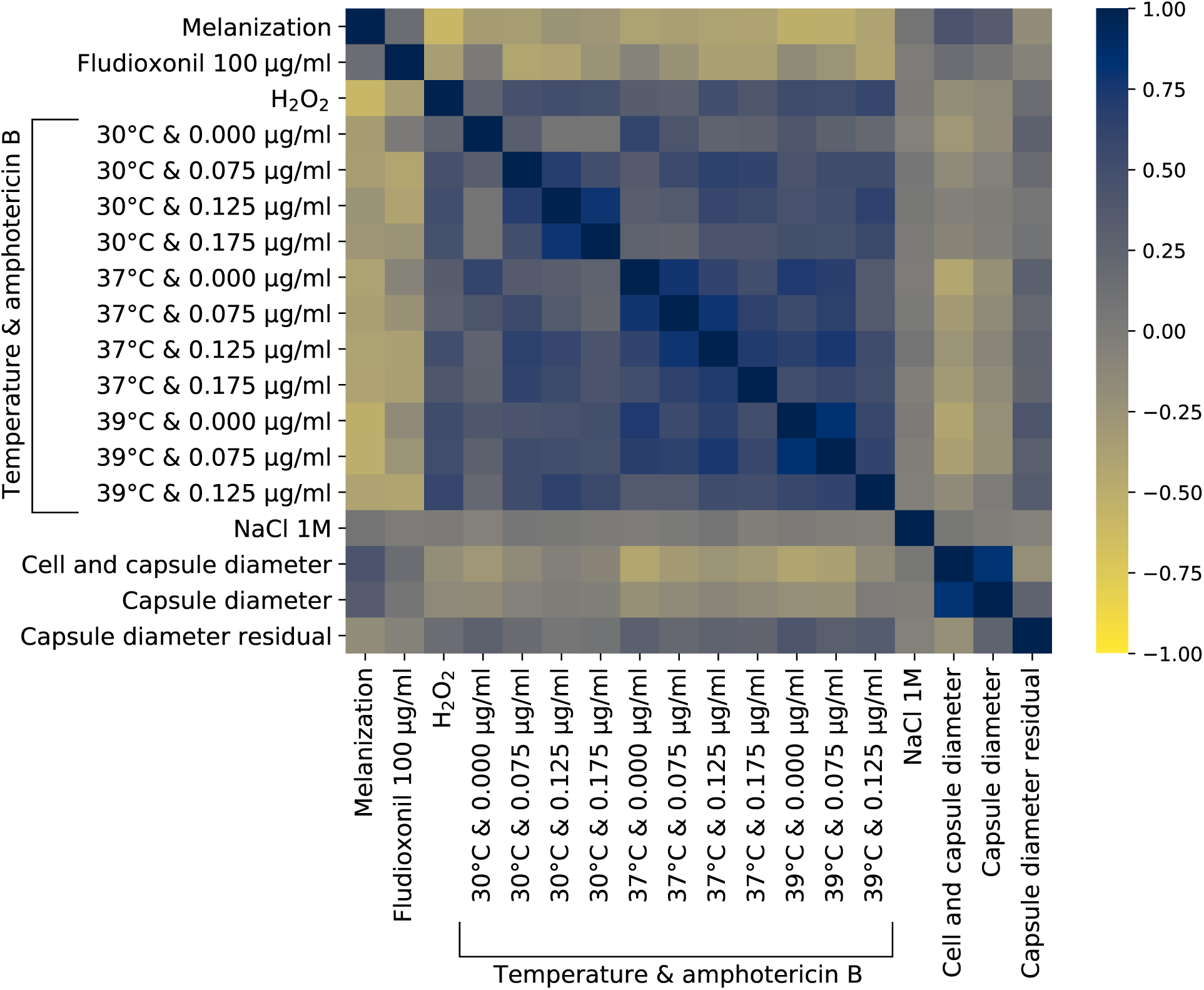
Phenotypic correlations. Spearman rank correlations between *C. deneoformans* phenotypes.

**S9 Fig.**
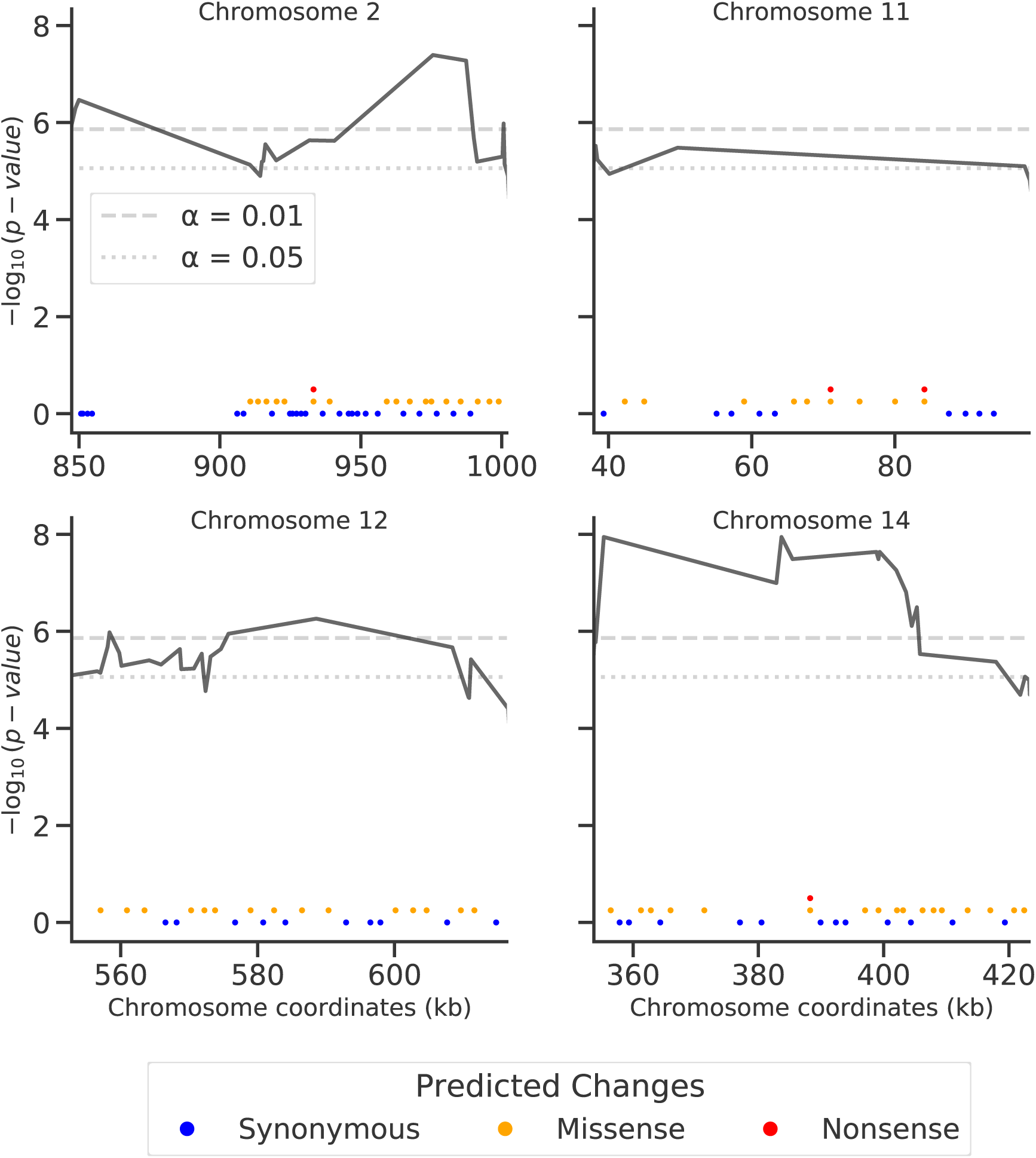
Changes in genes near QTL. Locations of genes, relative to the four identified QTL (black curves). From the JEC21α reference genome [208], features were aligned to the XL280α reference [137] and the changes and differences in protein sequence between XL280a, XL280α, and 431α were predicted. Dots along the x-axis represent location of mapped genes, colors indicate predicted change between the XL280a (or XL280α) and 431α parental strains.

**S10 Fig.**
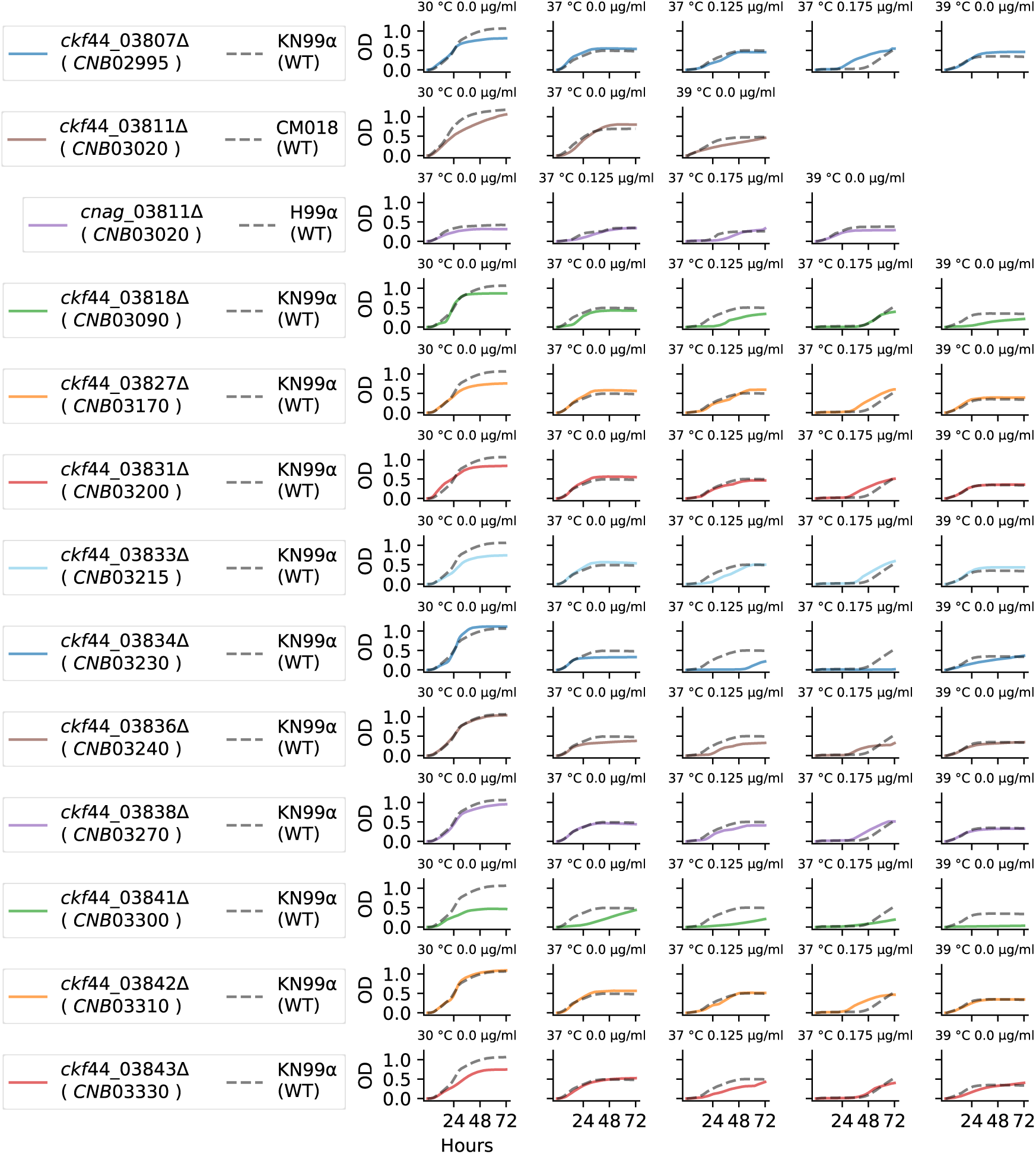
Phenotypes of chromosome 2 candidate deletion mutants. Growth of candidate deletion mutant strains for genes within the QTL along chromosome 2. The available deletion mutants (rows, solid curves) of genes within the QTL and the corresponding wild type, *C. neoformans* strain, were assayed for growth in liquid culture for 72 hour at high temperatures (30*^◦^*, 37*^◦^* and 39*^◦^*C) and in the presence of amphotericin B (at 0.125 and 0.175 *µ*g/ml). Legends on the far left show the gene names in the *C. neoformans* strain background with the corresponding *C. deneoformans* gene name.

**S11 Fig.**
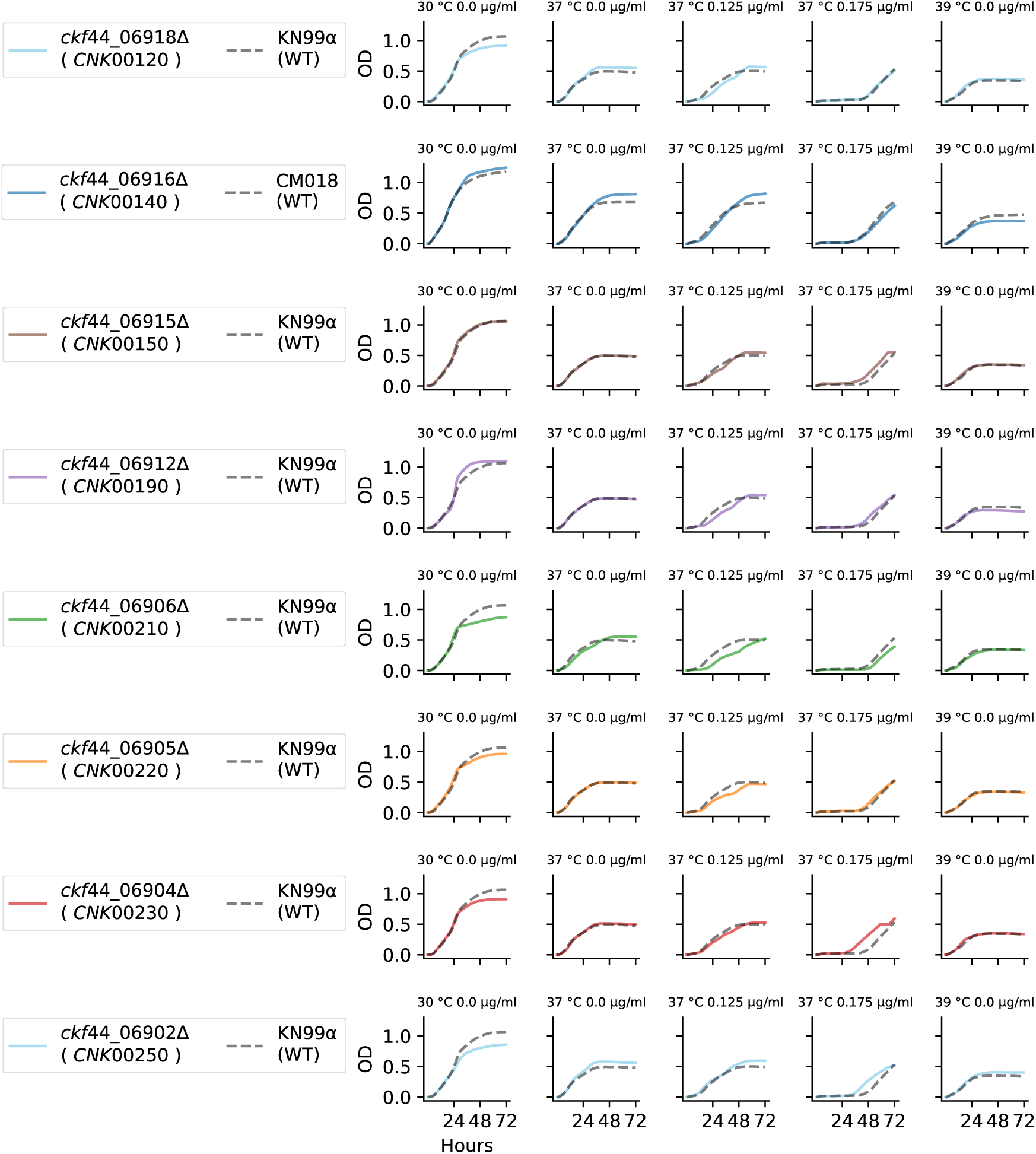
Phenotypes of chromosome 11 candidate deletion mutants. Growth of candidate deletion mutant strains for genes within the QTL along chromosome 11. The available deletion mutants (rows, solid curves) of genes within the QTL and the corresponding wild type, *C. neoformans* strain, were assayed for growth in liquid culture for 72 hour at high temperatures (30*^◦^*, 37*^◦^* and 39*^◦^*C) and in the presence of amphotericin B (at 0.125 and 0.175 *µ*g/ml). Legends on the far left show the gene names in the *C. neoformans* strain background with the corresponding *C. deneoformans* gene name.

**S12 Fig.**
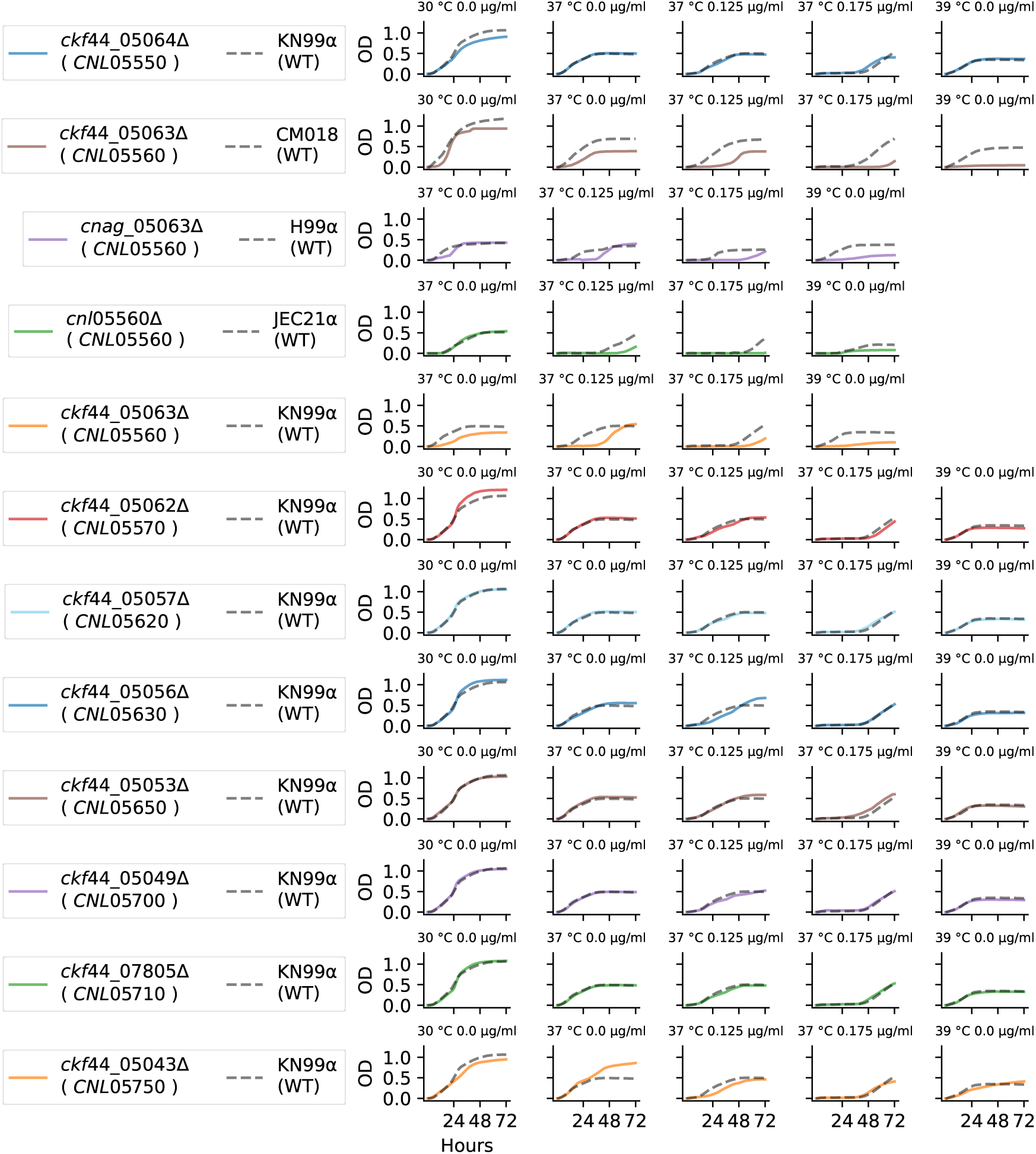
Phenotypes of chromosome 12 candidate deltion mutants. Growth of candidate deletion mutant strains for genes within the QTL along chromosome 12. The available deletion mutants (rows, solid curves) of genes within the QTL and the corresponding wild type, *C. neoformans* strain, were assayed for growth in liquid culture for 72 hour at high temperatures (30*^◦^*, 37*^◦^* and 39*^◦^*C) and in the presence of amphotericin B (at 0.125 and 0.175 *µ*g/ml). Legends on the far left show the gene names in the *C. neoformans* strain background with the corresponding *C. deneoformans* gene name.

**S13 Fig.**
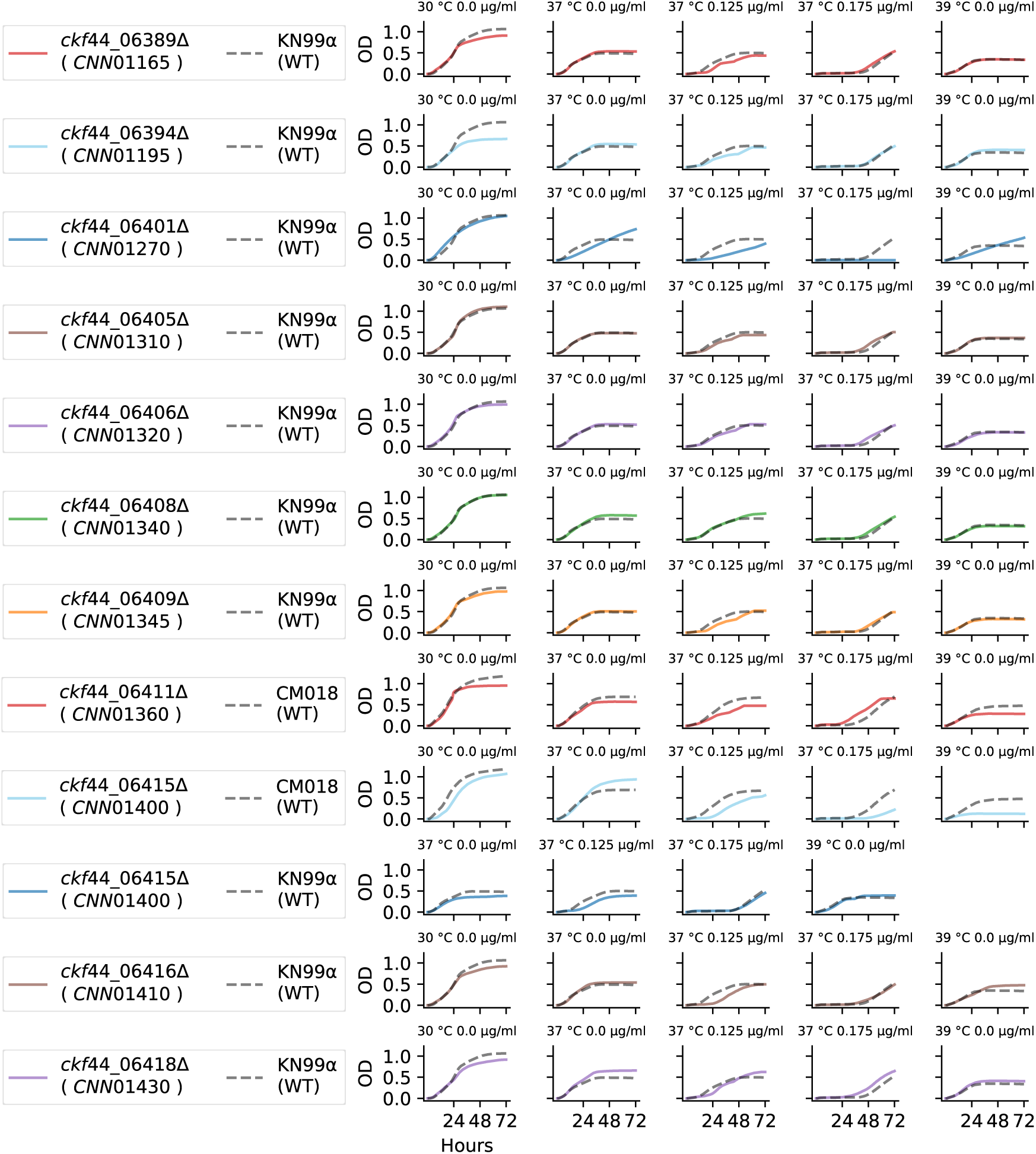
Phenotypes of chromosome 14 candidate deltion mutants. Growth of candidate deletion mutant strains for genes within the QTL along chromosome 14. The available deletion mutants (rows, solid curves) of genes within the QTL and the corresponding wild type, *C. neoformans* strain, were assayed for growth in liquid culture for 72 hour at high temperatures (30*^◦^*, 37*^◦^* and 39*^◦^*C) and in the presence of amphotericin B (at 0.125 and 0.175 *µ*g/ml). Legends on the far left show the gene names in the *C. neoformans* strain background with the corresponding *C. deneoformans* gene name.

**S14 Fig.**
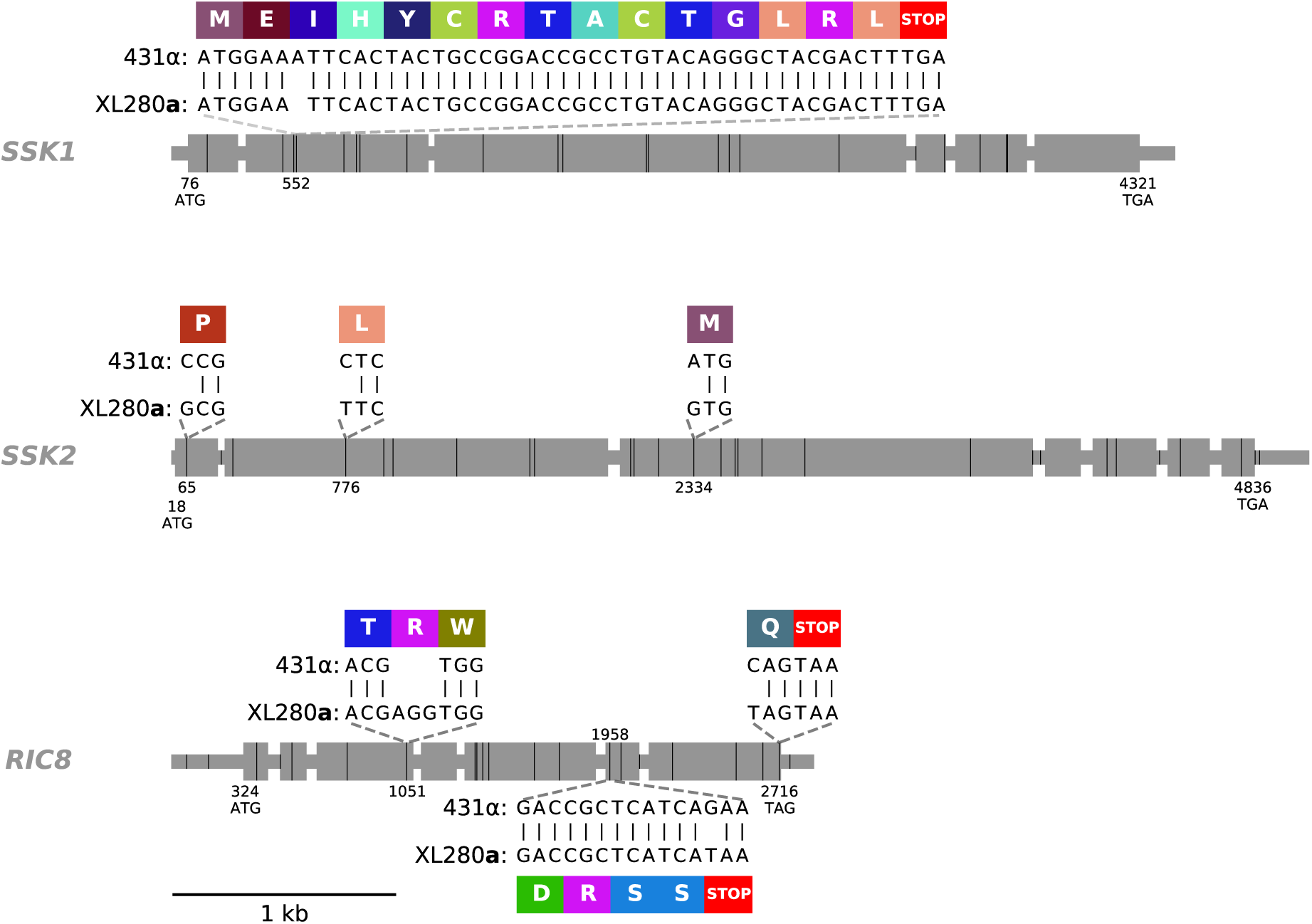
*SSK1*, *SSK2*, and *RIC8* gene models. Exons are shown as large grey rectangles, while the introns, 5’ UTR, and 3’ UTR are shown as grey, horizontal lines. The positions of bi-allelic genetic variants between the parental strains, 431α and XL280a are marked by black, vertical lines. The positions of the predicted start and stop codons are annotated along the bottom of the gene bodies. Within the second exon of *SSK1*, an insertion site of a single nucleotide, present in the 431α parental strain is annotated and this insertion is predicted to cause a frame shift that leads to a downstream early stop-gain. Within the first, second, and third exons of *SSK2*, three SNPs are annotated that lead to non-synonymous changes. The allelic states of the last two non-synonymous changes in *SSK2* have been previously identified by [143]. Within the third and last exon of *RIC8*, an in-frame codon deletion and shift in the predicted stop-codon (respectively) are seen in the 431α parental strain background. In the second to last exon of *RIC8*, a single-nucleotide polymorphism is present in the XL280a parental strain that is predicted to cause a premature stop. The local, predicted translations of the regions near these non-synonymous, genetic variants and associated amino acids are annotated in colored rectangles.

**S15 Fig.**
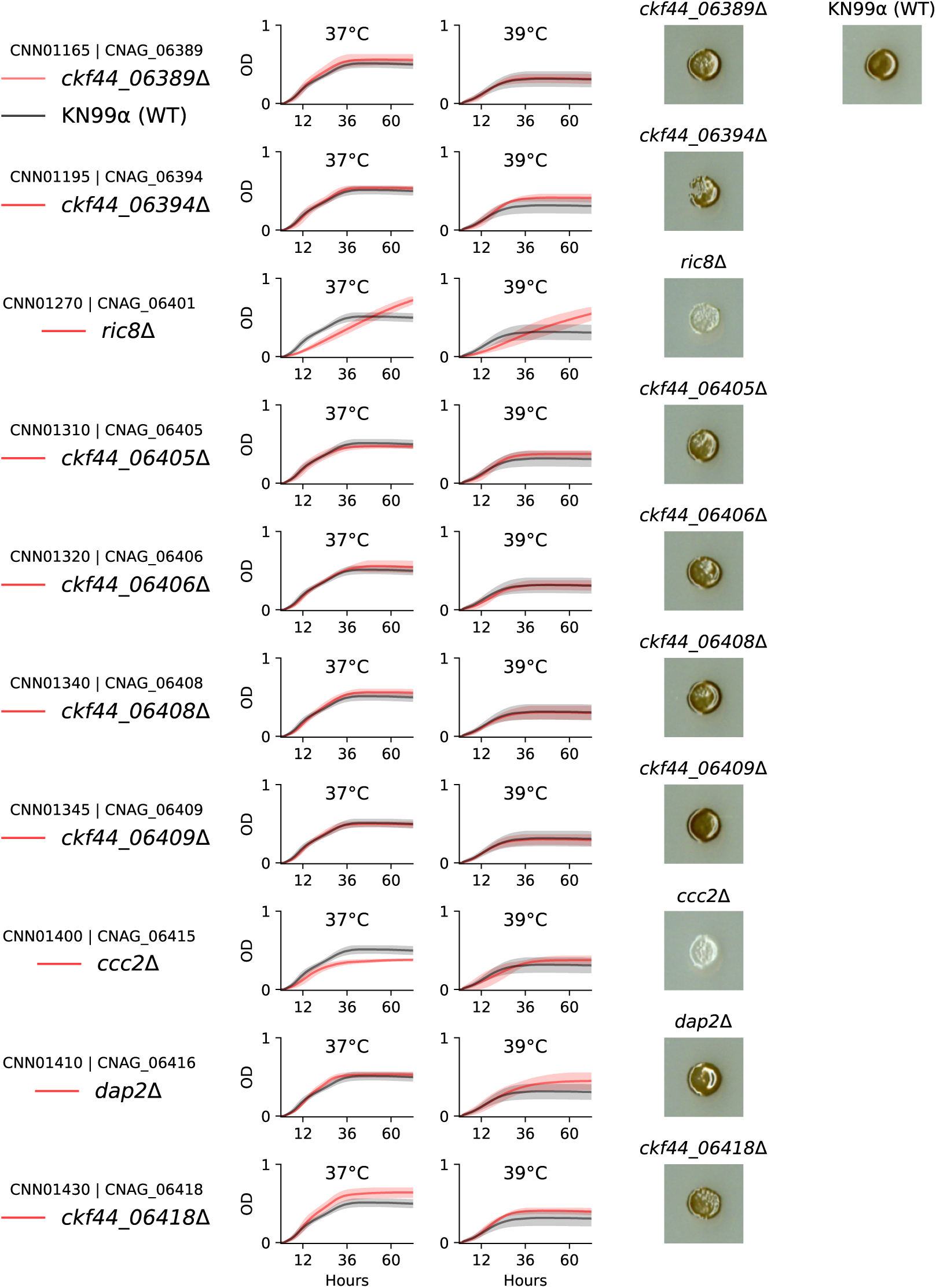
High temperature growth and melanization phenotypes. High temperature growth and melanization phenotypes of chromosome 14 candidate quantitative trait genes. The available deletion strains (rows) in the KN99α strain background of orthologous genes within the chromosome 14 QTL were assayed for high temperatures growth (37*^◦^* and 39*^◦^*C) in liquid culture and melanization on L-DOPA plates (columns, left to right respectively). Legends on the far left list the orthologous gene names in the *C. deneoformans* (JEC21α) and *C. neoformans* (H99α) background. Red and black curves display mean high temperature growth for the deletion strain and KN99α wild type (WT) strain (respectively) and shaded regions represent 95% confidence intervals.

**S16 Fig.**
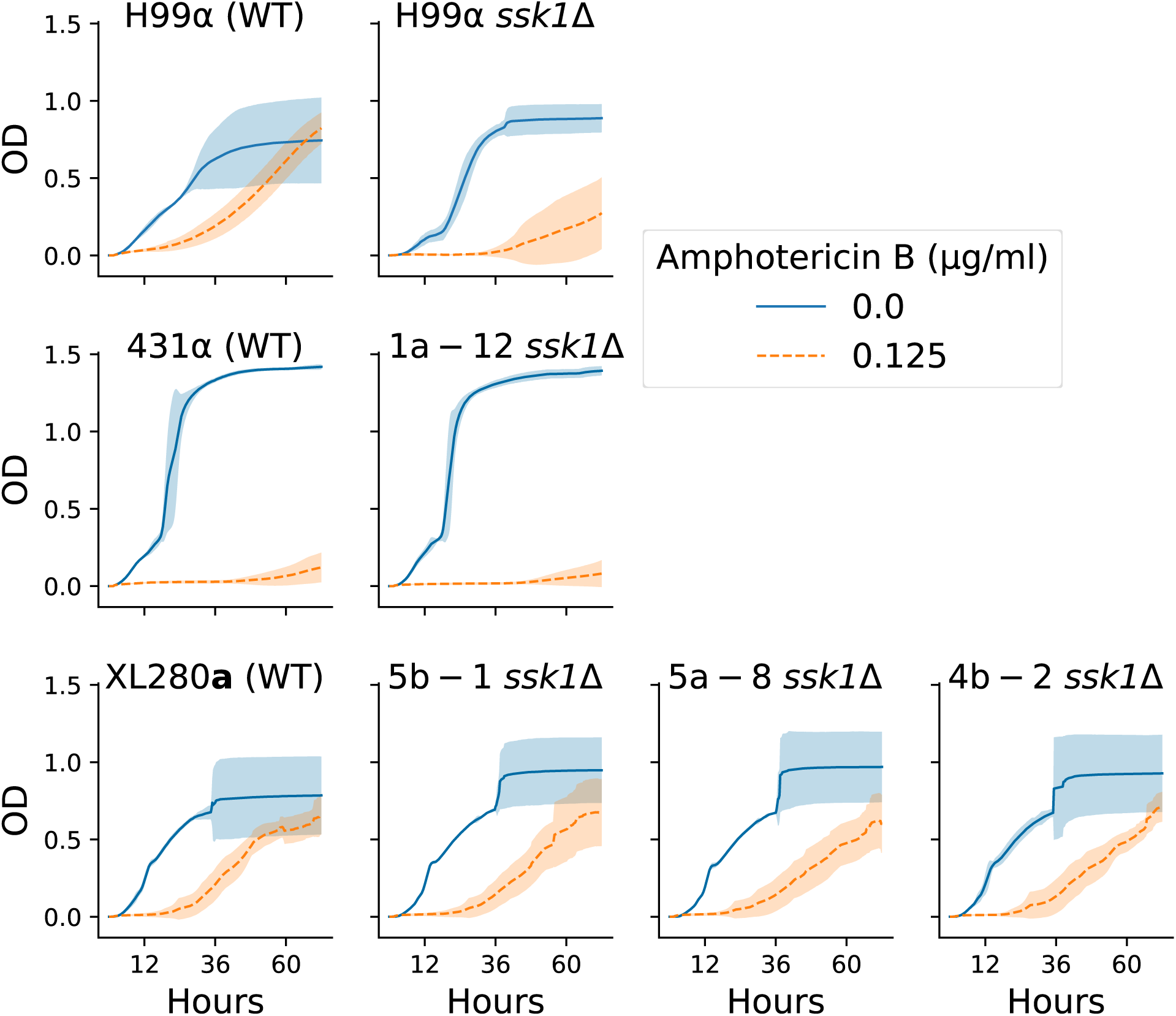
*C. neoformans* and *C. deneoformans ssk1* deletion mutant strain phenotypes. Growth of *C. neoformans* and *C. deneoformans ssk1* deletion mutant strains. Across the rows and columns, the growth in liquid culture of wild type (WT) and *ssk1*Δ strains (y-axis, optical density, 595 nm) incubated for 72 hour (x-axis) at 30*^◦^*C with and without amphotericin B (0.125 *µ*g/ml). Rows separate strain backgrounds. Growth curves are shown in the first column for the *C. neoformans* WT strain, H99α (first row) and the *C. deneoformans* strains XL280a and 431α (last two rows, respectively). The growth curves of *ssk1*Δ strains, per background, are depicted in the second, third, and fourth columns. Solid blue curves and dashed orange curves represent mean growth curves with 0.0 and 0.125 *µ*g/ml of amphotericin B (respectively) and shaded regions are 95% confidence intervals.

**S17 Fig.**
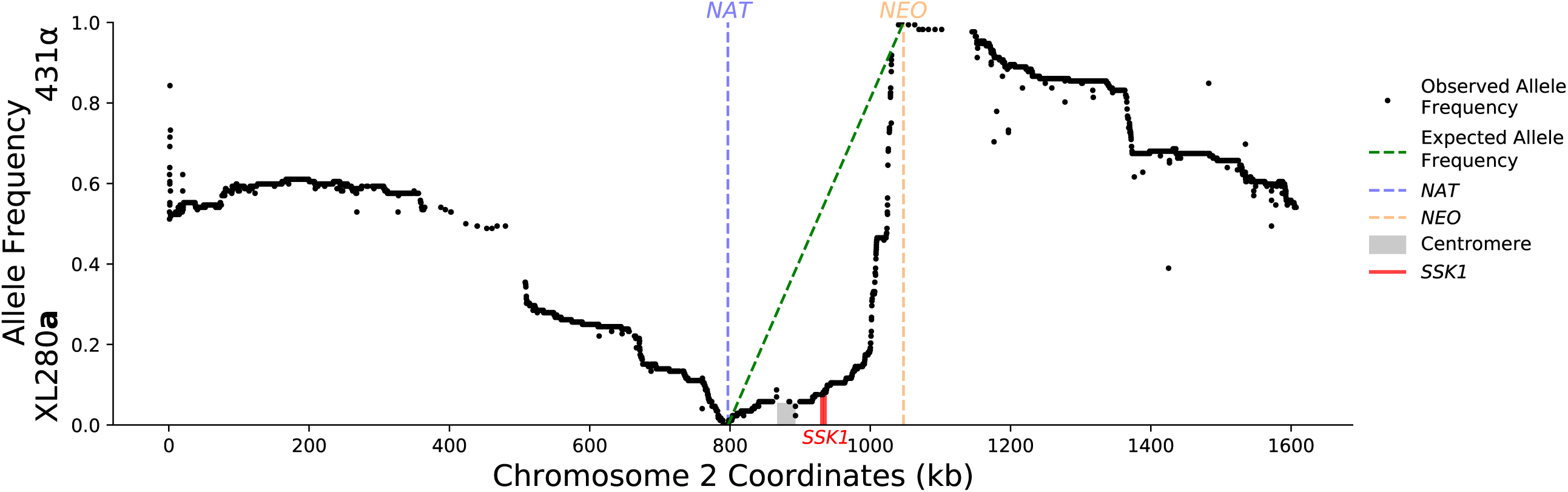
Chromosome 2 allele frequencies. Allele frequency across bi-allelic, genetic variant sites of chromosome 2. Across chromosome 2 (x-axis) for the progeny generated from fine mapping, the position and allele frequency (y-axis) of genetic variants between the parental strains XL280a and 431α are shown. The position of the selectable markers transformed within the parental backgrounds are shown by vertical, blue and orange, dashed lines. The expected allele frequency in this region given the marker locations is shown with a green, dashed line. The positions of the centromere and the *SSK1* gene are shown by grey and red rectangles, respectively.

**S18 Fig.**
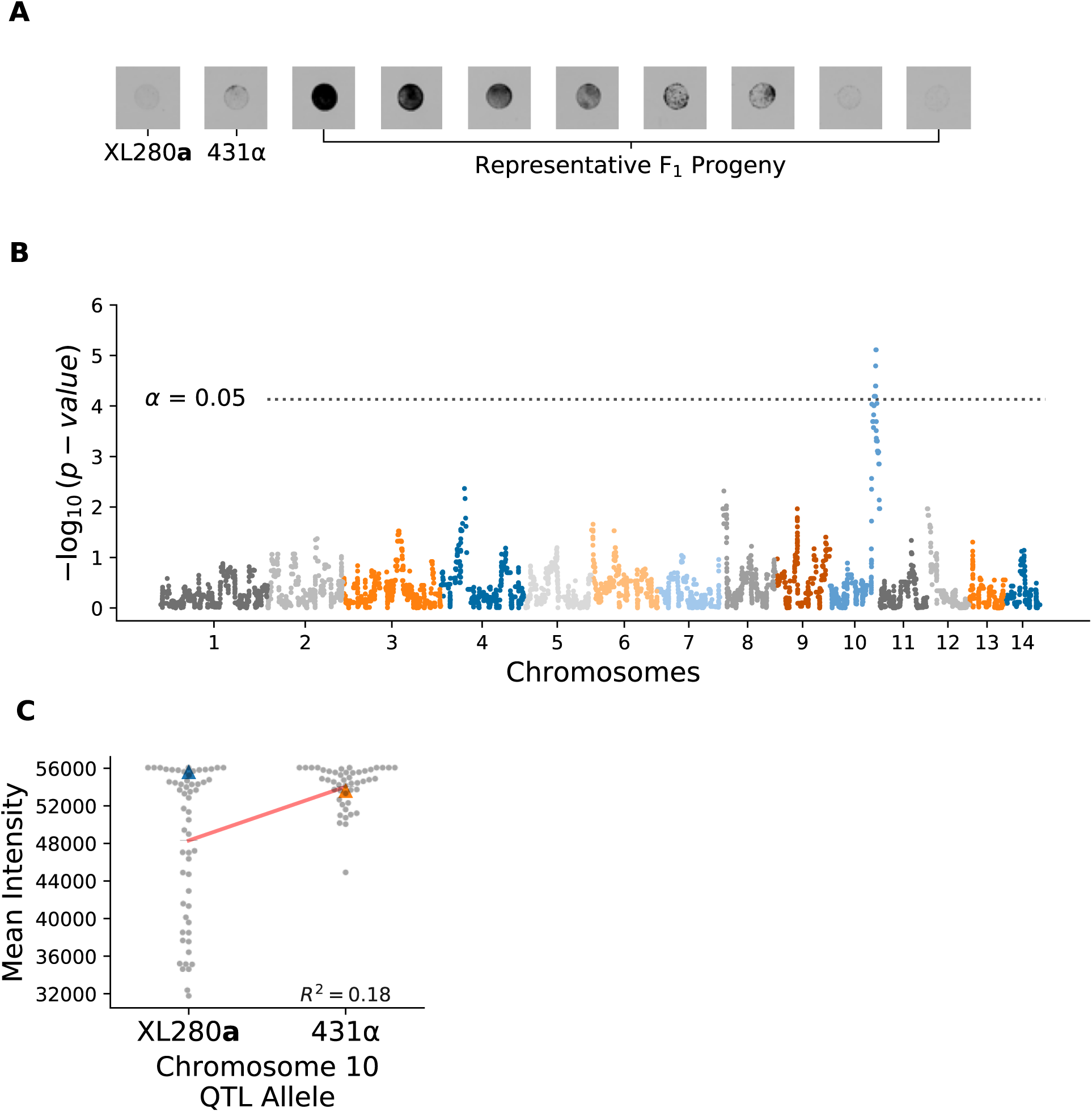
Osmotic shock QTL. QTL analysis of variation in response to osmotic shock. **A**) Growth phenotypes of parental strains grown on media with 1M NaCl and range of phenoytpes of their segregants. **B**) Manhattan plot of the association between genotype and growth in response to osmotic shock. The x-axis represents chromosomal locations of haploblocks and the y-axis represents the strength in association between genotype and variation in growth as measured by the mean intensity from translucent scans. **C**) Mean intensity (arbitrary units) of segregants (gray dots) from translucent scans as a function of allele at the peak of chromosome 10 QTL. The parental phenotypes are displayed by blue and orange triangles. Black horizontal lines denote the phenotypic means by allele and a red line represents a regression model relating genotype to phenotype. The heritablity – estimated from this regression model – is *∼*18 and annotated in black.

**S19 Fig.**
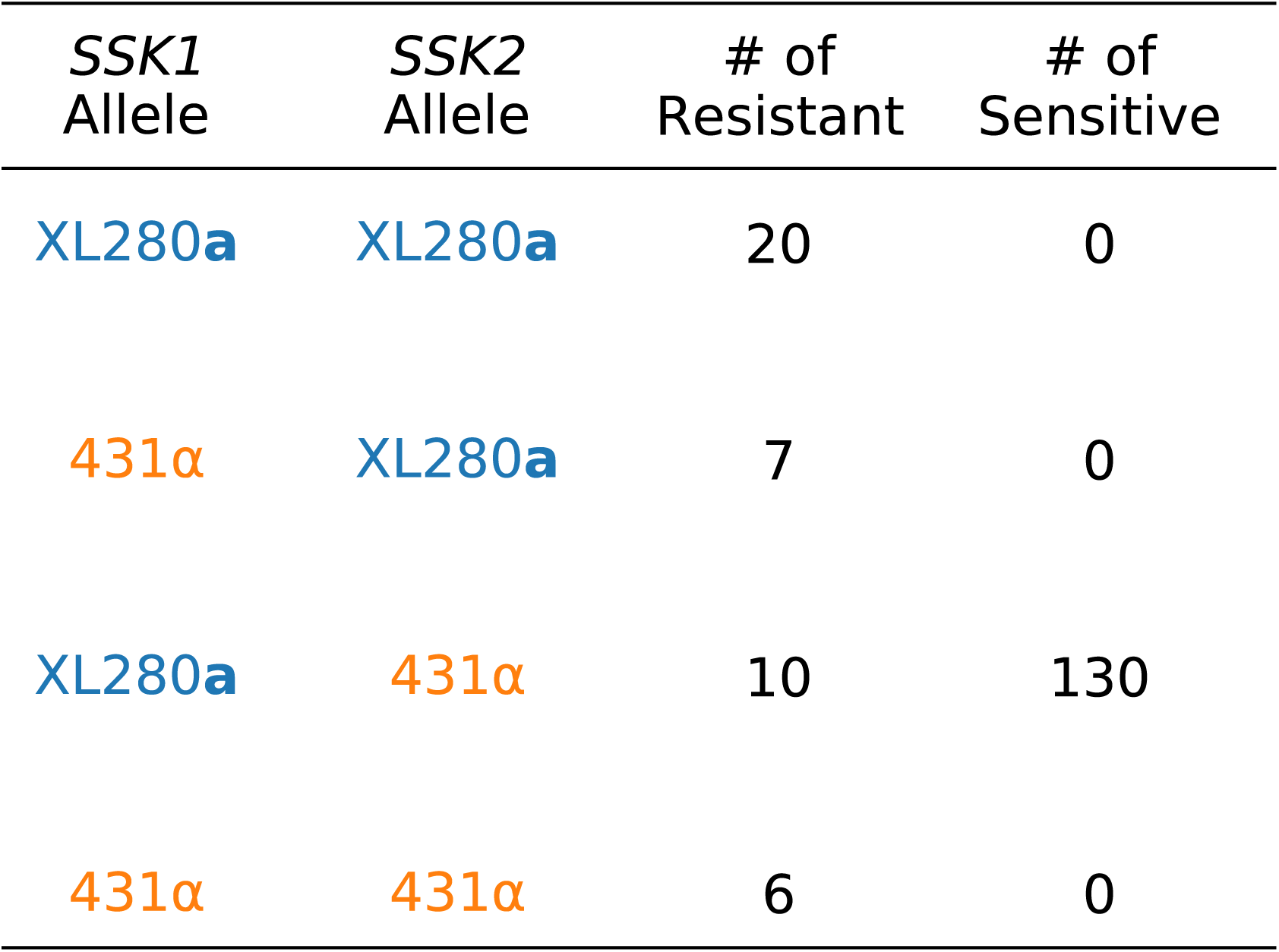
Fludioxonil phenotypes of additional *C. deneoformans* progeny. The segregants represent the possible combinations of the *SSK1* and *SSK2* alleles from the XL280a strain, CF1730, and the 431α strains, CF1705, CF1706, CF1707. Within these progeny, of the 140 progeny with the *SSK1* allele from XL280a parental strain and the *SSK2* allele from the 431α parental strains (second to last row), 130 (93%) demonstrated sensitivity to fludioxonil (100 *µ*g/ml). All other combinations of the parental alleles in the fine-mapping progeny demonstrated resistance to fludioxonil.

**S20 Fig.**
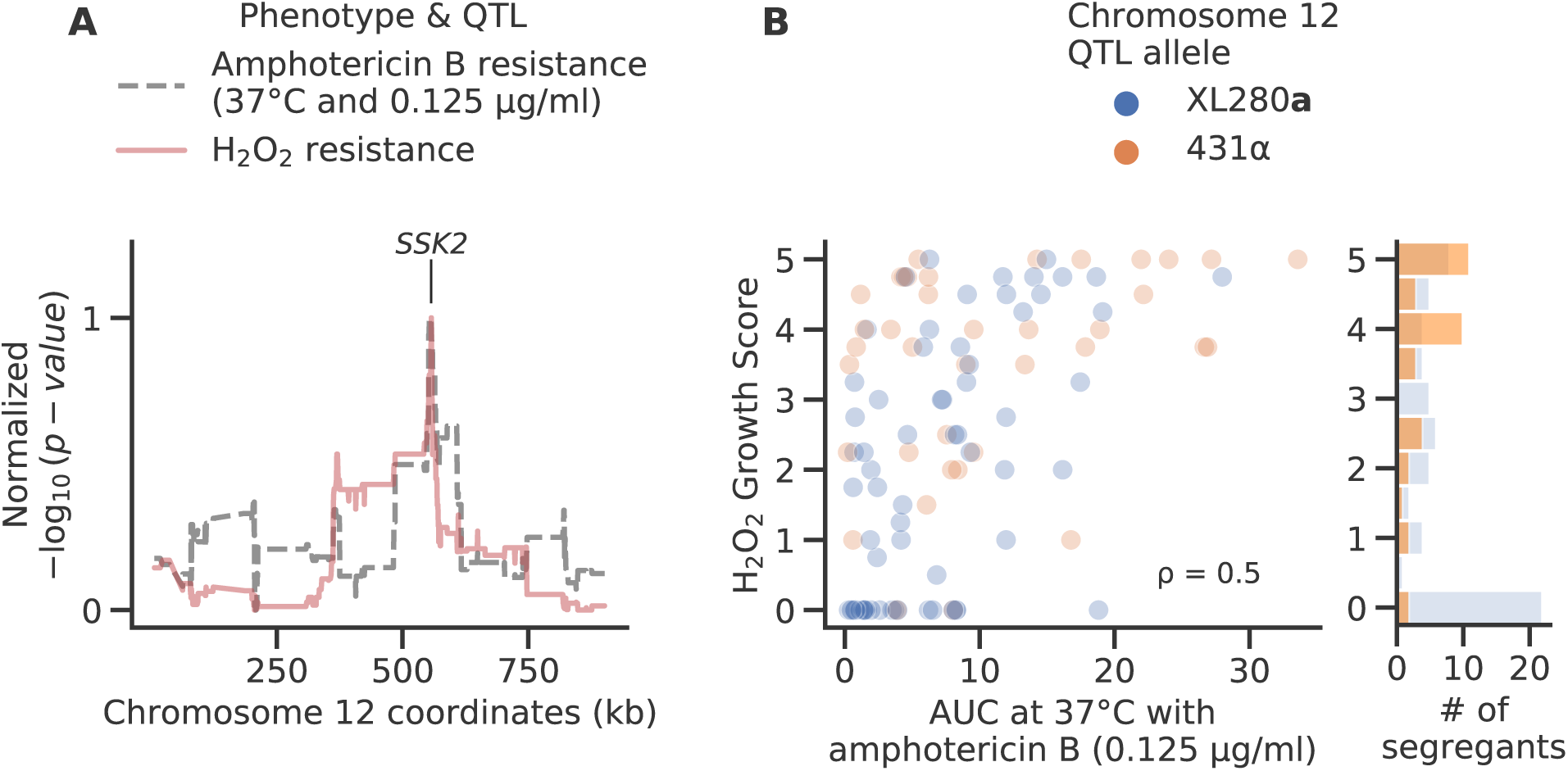
Pleiotropic chromosome 12 QTL. Chromosome 12 QTL and phenotypes for H_2_O_2_ and amphotericin B resistance. **A**) Normalized strength in association (y-axis) for the Chromosome 12 QTL for H_2_O_2_ (red) and amphotericin B resistance (black). The location of the candidate QTG, *SSK2* is annotated. **B**) Median H_2_O_2_ growth score (y-axis) as a function of *AUC* at 37*^◦^*C with 0.125 *µ*g/ml of amphotericin B (x-axis). The Spearman rank correlation is annotated within the plot. Segregant values are colored by their peak allele at chromosome 12 and a histogram in the right panel counts the number of segregants (x-axis) per H_2_O_2_ score.

**S21 Fig.**
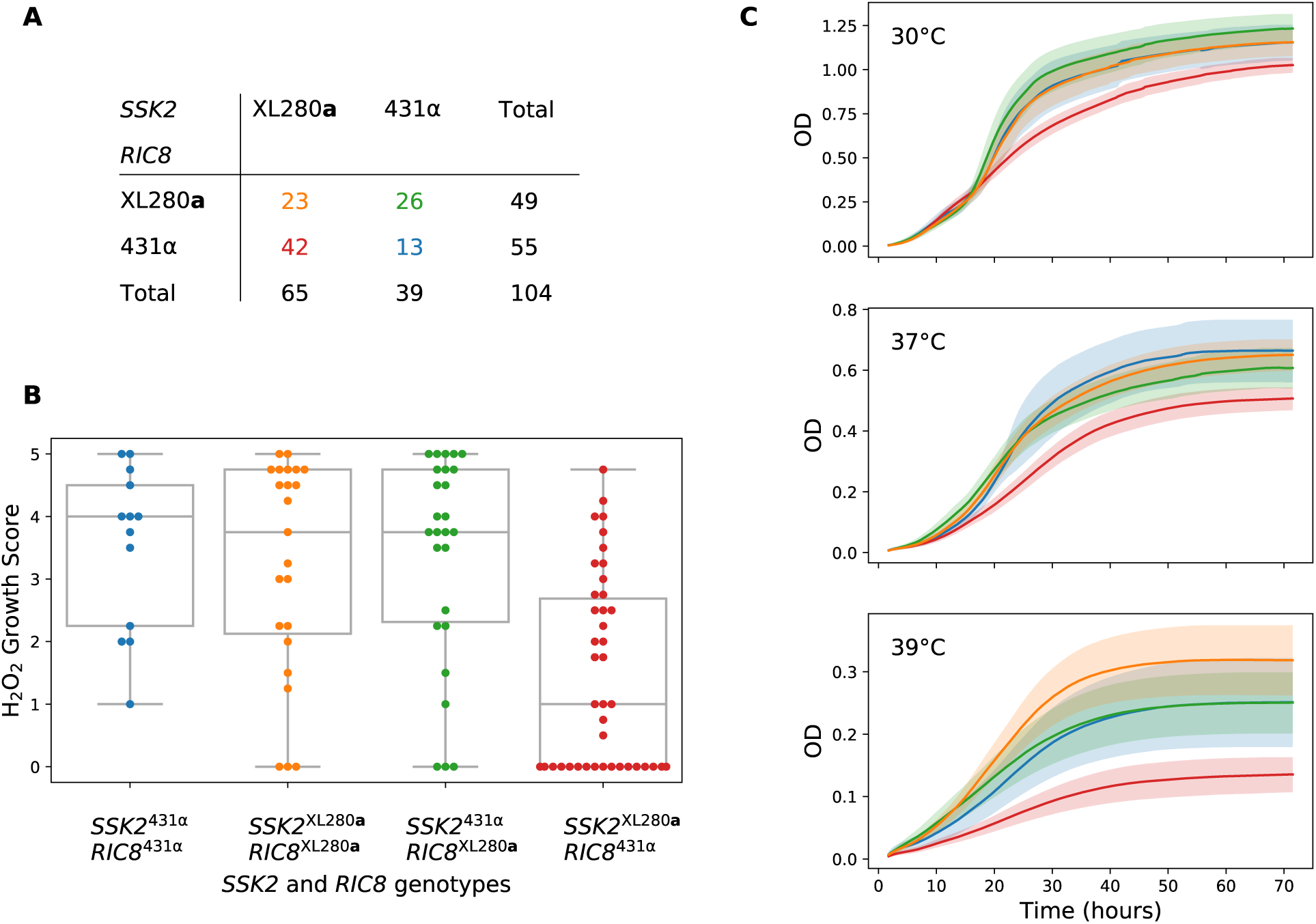
Effect of *SSK2* and *RIC8* alleles on H_2_O_2_ and thermal tolerance. **A**) Contingency table of *SSK2* (columns) and *RIC8* (rows) alleles across segregants. **B**) Box- and swarm-plots of H_2_O_2_ growth scores (y-axis) by allelic combinations of *SSK2* and *RIC8* (x-axis). **C**) The mean growth curves (solid lines) and 95% pointwise confidence intervals (shaded regions) per allelic combination of *SSK2* and *RIC8* across temperatures. OD is optical density sampled at 595nm. In panels **B** and **C**, phenotypes are color coded by the combinations of *SSK2* and *RIC8* alleles listed in **A**.

