## Supplemental Table 2 for "Pleiotropy and epistasis within and between signaling pathways defines the genetic architecture of fungal virulence"

| Name | Primer sequence 5' to 3' | Purpose | Citation |
|---|---|---|---|
| ***SSK1* gene disruption** | | | |
| CRISPR- Cas9 | | | |
| Ssk1 P1 1000 | CCCAAAAGAGGGTGGATGCT | construction of deletion cassette | |
| Ssk1 P2 1000 | CTGGCCGTCGTTTTACGCCCCATTGTAACACCTCCA | construction of deletion cassette | |
| Ssk1 P3 1000 | GTCATAGCTGTTTCCTGTGATGATCGAAGCCCGACAA | construction of deletion cassette | |
| Ssk1 P4 1000 | TTATCCATTCTGCCGGTCGC | construction of deletion cassette | |
| Ssk1 P5 1000 | GGCCTCAATGGCGTAGACTT | construction of deletion cassette | |
| Ssk1 P6 1000 | CTGTTCAGGAGCAGAGCCAA | construction of deletion cassette | |
| Ssk1 sgF Cd1 | GTCCCGTATCCTCGAGCTCAGGTTTTAGAGCTAGAAATAGCAAGTT | construction of gRNA cassette | |
| Ssk1 sgR Cd1 | CTGAGCTCGAGGATACGGGACAACAGTATACCCTGCCGGTG | construction of gRNA cassette | |
| U6 F | TTTGCATTAGAACTAAAAACAAAGCA | construction of gRNA cassette | Fan and Lin 2018 |
| gRNA R | TAAAACAAAAAAGCACCGACTCGGTGCC | construction of gRNA cassette | Fan and Lin 2018 |
| U6 FLF | GGCTCAAAGAGCAGATCAATG | construction of gRNA cassette | Fan and Lin 2018 |
| sgRNA FR | CCTCTGACACATGCAGCTCC | construction of gRNA cassette | Fan and Lin 2018 |
| GPD1-P-F | CATGCATCTAGGTCTAGAAACC | amplifying Cas9 from pXL1-Cas9-HygB | |
| GPD1-T-R | CCTCTTCACGTGGACGCTCC | amplifying Cas9 from pXL1-Cas9-HygB | |
| Confirmation of transformants | | | |
| *SSK1* F3 | CGTCGGACTTCCCATAGCAT | detecting presence of *SSK1* | |
| *SSK1* R3 | ATCCTTCGTTTTGTCGGCCT | detecting presence of *SSK1* | |
| Cas9 intF | CGACCTCCGTCTCATCTACC | detecting presence of Cas9 | |
| Cas9 intR | TTGAGGAGGGTGAGGTCTTG | detecting presence of Cas9 | |
| *SSK1* A | TAGTTCGAGGAAACGCGGAG | confirmation of boundaries, length | |
| *SSK1* B | TGTTACGGAGCGATGGTGAC | confirmation of boundaries, length | |
| *SSK1 NAT* B | CTGGCGGAGGATAGAAGCTG | confirmation of boundaries, length | |
| *SSK1* C | TCGCTTGCTAGCAGACTACG | confirmation of boundaries, length | |
| *SSK1* D | ACTTGGATTAGGCATTGCATGT | confirmation of boundaries, length | |
| *SSK1 NAT* C | TCGGGTCAATTGTCTCAGTCG | confirmation of boundaries, length | |
| Sequencing | | | |
| *SSK1* 1F | CCAAGCATCAGTCATGCACC | sequencing of transformants | |
| *SSK1* 1R | CGCGTTACTTTTCTTCGCGT | sequencing of transformants | |
| *SSK1* 2F | AATGGAAAGACTCGGCTGGC | sequencing of transformants | |
| *SSK1* 2R | GGAAGCATTTGAGCCCCACA | sequencing of transformants | |
| *SSK1* 2R *NAT* | ATGTAAGTCCGCTCCTTCCC | sequencing of transformants | |
| *SSK1* 3F *NAT* | AGAATTCGCCCTTAGGCTGC | sequencing of transformants | |
| *SSK1* 3R *NAT* | CTCTGTCCAACGCACATCCA | sequencing of transformants | |
| *SSK1* 4F *NAT* | GCAACAGCCCATCCTTGTTG | sequencing of transformants | |
| *SSK1* 4R *NAT* | AAGAGCTTGCTCTCCGTCAG | sequencing of transformants | |
| *SSK1* 5F *NAT* | TTGTGGACTGGATACCGCAC | sequencing of transformants | |
| *SSK1* 5R | TAGGAGAGCTGGTCGACTCC | sequencing of transformants | |
| *SSK1* 6F | ACATGCAATGCCTAATCCAAGT | sequencing of transformants | |
| *SSK1* 6R | CAACCCCAACGGAGGAGAAA | sequencing of transformants | |
| *SSK1* 7F | GAGTTTGTTCATGGCAGAGGC | sequencing of transformants | |
| *SSK1* 7R | AGCCATGGTTTCTTTCCAGTAA | sequencing of transformants | |
| **Chr2 QTL fine mapping** | | | |
| CRISPR- Cas9 | | | |
| QTL-L-5F | TGACTTCCCCTGGTTCCTAT | QTL left border *NAT* cassette | |
| QTL-L-5R | ACTGGCCGTCGTTTTACACCCGATGATTTTCCGAC | QTL left border *NAT* cassette | |
| QTL-L-3F | GTCATAGCTGTTTCCTGAAAGGAAGACGACGGGAAGT | QTL left border *NAT* cassette | |
| QTL-L-3R | AAAGCCAAGGACGAGGTCA | QTL left border *NAT* cassette | |
| QTL-R-5F | TGAAAACCGAAAACCCTGAC | QTL right border *NEO* cassette | |
| QTL-R-5R | ACTGGCCGTCGTTTTACACGCCTTCGTCACTCAAAAC | QTL right border *NEO* cassette | |
| QTL-R-3F | GTCATAGCTGTTTCCTGAGCAGTTGGCGTTAGCAGTT | QTL right border *NEO* cassette | |
| QTL-R-3R | GGACCTCTTCGTATTTTCAAGGAC | QTL right border *NEO* cassette | |
| QTL-L-F | ATGAGAGGTGGAACCGAGAG | QTL left border genotyping | |
| QTL-L-R | TCAAGTCCAACAAGCAGTGG | QTL left border genotyping | |
| QTL-R-F | AACTGGGCTGGGTCCATACA | QTL right border genotyping | |
| QTL-R-R | GTTCGGGAGCGTTCGTTAGA | QTL right border genotyping | |
| *NAT*-Southern-F | CGATACGGCTTACCGTTACAG | *NAT* southern probe | |
| *NAT*-Southern-R | GAGCTTGCTCTCCGTCAGATG | *NAT* southern probe | |
| *NEO*-Southern-F | GAAGGGACTGGCTGCTATTG | *NEO* southern probe | |
| *NEO*-Southern-R | GAACTCGTCAAGAAGGCGATA | *NEO* southern probe | |
